# Comparison Between Gradients and Parcellations for Functional Connectivity Prediction of Behavior

**DOI:** 10.1101/2022.09.22.509045

**Authors:** Ru Kong, Yan Rui Tan, Naren Wulan, Leon Qi Rong Ooi, Seyedeh-Rezvan Farahibozorg, Samuel Harrison, Janine D. Bijsterbosch, Boris C. Bernhardt, Simon Eickhoff, B.T. Thomas Yeo

## Abstract

Resting-state functional connectivity (RSFC) is widely used to predict behavioral measures. To predict behavioral measures, representing RSFC with parcellations and gradients are the two most popular approaches. Here, we compare parcellation and gradient approaches for RSFC-based prediction of a broad range of behavioral measures in the Human Connectome Project (HCP) and Adolescent Brain Cognitive Development (ABCD) datasets. Among the parcellation approaches, we consider group-average “hard” parcellations (Schaefer et al., 2018), individual-specific “hard” parcellations (Kong et al., 2021a), and an individual-specific “soft” parcellation (spatial independent component analysis with dual regression; Beckmann et al., 2009). For gradient approaches, we consider the well-known principal gradients (Margulies et al., 2016) and the local gradient approach that detects local RSFC changes (Laumann et al., 2015). Across two regression algorithms, individual-specific hard-parcellation performs the best in the HCP dataset, while the principal gradients, spatial independent component analysis and group-average “hard” parcellations exhibit similar performance. On the other hand, principal gradients and all parcellation approaches perform similarly in the ABCD dataset. Across both datasets, local gradients perform the worst. Finally, we find that the principal gradient approach requires at least 40 to 60 gradients to perform as well as parcellation approaches. While most principal gradient studies utilize a single gradient, our results suggest that incorporating higher order gradients can provide significant behaviorally relevant information. Future work will consider the inclusion of additional parcellation and gradient approaches for comparison.

## Introduction

Resting-state functional connectivity (RSFC) reflects the synchrony of fMRI signals between brain regions, while a subject is lying at rest without performing any explicit task (Biswal et al., 1995; Greicius et al., 2003; Fox and Raichle, 2007). There is significant interest in using RSFC for predicting individual differences in behavior (van den Heuvel et al., 2009; Finn et al., 2015; Dubois et al., 2018a; Rosenberg et al., 2020). For example, machine learning techniques have been used to learn the relationship between RSFC patterns and fluid intelligence in a large sample of participants. Given the RSFC pattern of a new participant, the goal is to use the learned relationship to predict the fluid intelligence of the new participant.

To predict behavioral measures, there are different approaches for representing RSFC data (Bijsterbosch et al., 2020) with the two most popular approaches being parcellations (Smith et al., 2009; Power et al., 2011; Yeo et al., 2011; Glasser et al., 2016) and gradients (Cohen et al., 2008; Margulies et al., 2016; Haak et al., 2018; Tian et al., 2020). Since different parcellation and gradient approaches might capture different aspects of brain organization, we compared different parcellation and gradient approaches for RSFC-based behavioral prediction.

Example parcellation approaches include hard-parcellation approaches that estimate non-overlapping regions of interest (ROIs) (Shen et al., 2013; Glasser et al., 2016; Schaefer et al., 2018), and soft-parcellation approaches that estimate overlapping ROIs (Calhoun et al., 2001; Beckmann et al., 2005; Smith et al., 2009; Zuo et al., 2010; Lee et al., 2012; Harrison et al., 2015; Farahibozorg et al., 2021). Most studies have utilized RSFC from population-average brain parcellations to predict behavior measures. Recent studies have shown that individual-specific parcellation topography is behaviorally relevant (Bijsterbosch et al., 2018; Kong et al., 2019; Cui et al., 2020). Functional connectivity derived from individual-specific parcellations could further improve the prediction performance compared with population-average parcellations (Li et al., 2019b; Farahibozorg et al., 2021; Kong et al., 2021a).

Besides parcellation approaches, many studies have also utilized gradient techniques to characterize brain organization(Huntenburg et al., 2018; Bernhardt et al., 2022). For example, the local gradient approach detects local changes (i.e., gradients) in RSFC across the cortex (Cohen et al., 2008; Wig et al., 2014; Laumann et al., 2015; Gordon et al., 2016), which has been widely used for estimating hard-parcellations. Recent studies (Bijsterbosch et al., 2018; Kong et al., 2019) have suggested that network topography is behaviorally meaningful, so we hypothesize that local gradient maps can also be used to predict behavior. On the other hand, gradients have also been derived using manifold learning algorithms, such as diffusion embedding (also referred to as principal gradients; Margulies et al., 2016), principal component analysis (PCA; Hong et al. 2020) and Laplacian eigenmaps (LE) (Haak et al., 2018; Tian et al., 2020). Therefore, in this study, we considered both local and principal gradients for predicting behavior.

There have been previous comparisons of various parcellations approaches for predicting behavioral measures (Dadi et al., 2019; Pervaiz et al., 2020; Farahibozorg et al., 2021). These studies typically found that soft parcellations (e.g., ICA dual regression) performed better than group-level hard parcellations. However, these studies did not consider the use of individual-specific hard parcellation approaches (e.g., Kong et al., 2021). Furthermore, these studies found that RSFC computed using full correlation (Pearson’s correlation) performed worse than partial correlation. Given the increased popularity of gradient approaches, there is a need to compare prediction performance across various parcellation and gradient approaches.

One recent study has suggested that RSFC gradients (based on PCA, diffusion embedding and Laplacian eigenmaps) resulted in better prediction performance than parcellation-based RSFC (Hong et al., 2020). However, prediction with parcellation-based RSFC was performed using connectome predictive modeling (Shen et al., 2017), while prediction with gradient approaches was performed using canonical correlation analysis (Smith et al., 2015), so it is somewhat challenging to directly compare the two results. Furthermore, their prediction analyses were performed only in the Human Connectome Project (HCP) dataset, which is one of the most widely used dataset for investigating individual differences in behaviors. Repeated reuse of the same dataset by multiple researchers can lead to inflated error rates (Thompson et al., 2020).

Additionally, repeatedly using the same dataset for training and testing can cause the model overfit to that dataset, resulting in overly optimistic prediction results and less generalizable models to new datasets (Recht et al., 2019; Beyer et al., 2020). This emphasizes the importance of replicating analyses using additional less commonly used datasets. In the current study, in addition to the widely used HCP dataset, we replicate our analyses using the adolescent brain cognitive development (ABCD) dataset.

In this study, we compared different parcellation and gradient approaches for RSFC prediction of behavioral measures across a wide range of behavioral measures in two different datasets using two different prediction models. We considered a group-level hard-parcellation approach (Schaefer et al., 2018), an individual-specific hard-parcellation approach (Kong et al., 2021a), an individual-specific soft-parcellation approach based on ICA dual regression (Calhoun et al., 2001; Beckmann et al., 2005; Smith et al., 2009; Zuo et al., 2010; Nickerson et al., 2017), the principal gradients (Margulies et al., 2016), and the local gradient approach (Wig et al., 2014; Laumann et al., 2015; Gordon et al., 2016). Furthermore, we considered different resolutions (i.e., number of ROIs or gradients) for each approach. To compare the prediction performance across different approaches, the resolution was optimized as a hyperparameter in the prediction model. In a separate analysis, we investigated prediction performance as a function of the number of ROIs or gradients.

## Methods

### Datasets

We considered two publicly available datasets: the Human Connectome Project (HCP) S1200 release (Van Essen et al., 2012a; Smith et al., 2013) and the Adolescent Brain Cognitive Development (ABCD) 2.0.1 release. Both datasets contained structural MRI, resting-state fMRI (rs-fMRI), and multiple behavioral measures for each subject. After strict pre-processing quality control of the HCP and ABCD datasets based on our previous studies (Li et al., 2019a; Kong et al., 2021a; Chen et al., 2022), we considered participants with all four rs-fMRI scans remaining as well as all behavioral scores of interest. Our main analysis comprised 746 participants from HCP and 1476 participants from ABCD.

### Preprocessing

Details of the HCP preprocessing can be found elsewhere (Van Essen et al., 2012a; Glasser et al., 2013; Smith et al., 2013). The HCP rs-fMRI data has been projected to the fs_LR32k space (Van Essen et al., 2012b), denoised with ICA-FIX (Griffanti et al., 2014; Salimi-Khorshidi et al., 2014) and aligned with MSMAll (Robinson et al., 2014). Consistent with our previous studies (Li et al., 2019a; He et al., 2020), we further applied global signal regression (GSR) and censoring to eliminate global and head motion related artifacts. More details of the processing can be found elsewhere (Li et al., 2019a). Runs with more than 50% censored frames were removed. Participants with all four rs-fMRI runs remaining (N = 835) were considered.

Details of the ABCD preprocessing can be found elsewhere (Casey et al., 2018; Hagler et al., 2019). We utilized the minimally preprocessed functional data with additional processing steps including T1-T2* registration, respiratory pseudomotion motion filtering, nuisance regression, censoring and bandpass filtering. Nuisance regressors comprised global signal, ventricular signal, white matter signal, and six motion parameters, as well as their temporal derivatives. The data was then projected to the FreeSurfer fsaverage6 surface space and smoothed using a 6 mm full-width half maximum kernel. More details of the processing can be found elsewhere (Chen et al., 2022). Of the 2264 unrelated participants considered in our previous study (Chen et al., 2022), 1476 subjects had four runs of rs-fMRI data.

### Functional connectivity features for behavioral prediction

Here, we compared functional connectivity behavioral prediction across different parcellation and gradient approaches, including a group-level hard-parcellation approach (Schaefer et al., 2018), an individual-specific hard-parcellation approach (Kong et al., 2021a), a individual-specific soft-parcellation approach (Calhoun et al., 2001; Beckmann et al., 2005; Smith et al., 2009; Zuo et al., 2010; Nickerson et al., 2017), the principal gradient (Margulies et al., 2016), and the local gradient (Wig et al., 2014; Laumann et al., 2015; Gordon et al., 2016).

The different parcellation and gradient approaches were applied to each participant from the HCP and ABCD datasets using all rs-fMRI scans. We then estimated the functional connectivity features for each participant based on the derived parcellations and gradients (Figure 1):

**Figure 1.**
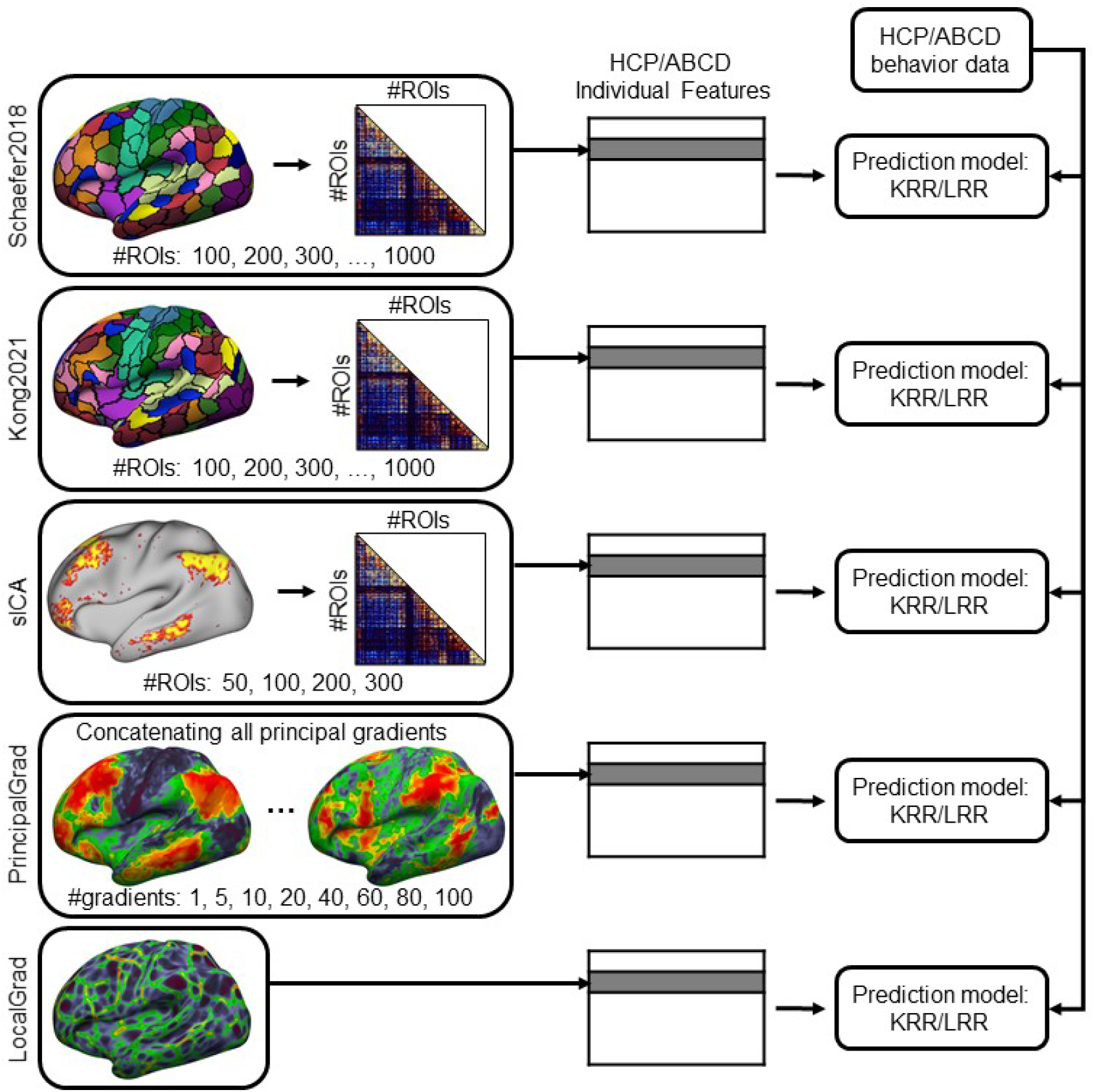
Flowchart across different parcellation and gradient approaches for RSFC-based behavioral prediction. For three parcellation approaches Schaefer 2018, Kong 2021 and sICA, the lower triangular part of the RSFC matrix was vectorized for each participant to serve as the individual-level RSFC features. A 50 × 50 RSFC matrix correspond to 1225 features, while a 1000 × 1000 RSFC matrix correspond to 499500 features. For the principal gradient approach PrincipalGrad, the principal gradients for each participant were concatenated together to serve as the individual-level RSFC features. At one extreme, 1 principal gradient comprised 59412 features for the HCP dataset (fs_LR32k space) and 74947 features for the ABCD dataset (fsaverage6 space). At the other extreme, 100 principal gradients utilized 5941200 features for the HCP dataset and 7494700 features for the ABCD dataset. For the local gradient approach LocalGrad, the local gradient map was used as the individual-level RSFC features. A local gradient map comprised 59412 features for the HCP dataset and 74947 features for the ABCD dataset. For each approach, we performed behavioral prediction for HCP and ABCD datasets separately using two different prediction models KRR and LRR.

1. *Group-level hard-parcellation Schaefer 2018*. Group-level hard-parcellations are estimated by averaging or concatenating data across many individuals, where each vertex is assigned to one region of interest (ROI). In our previous work, we developed a set of high-quality group-level hard-parcellations of the cerebral cortex with multiple resolutions from 100 to 1000 ROIs (Schaefer et al., 2018), which we will refer to as “Schaefer 2018”. For each parcellation resolution, the Schaefer 2018 parcellation was applied to all rs-fMRI scans of each participant to generate a resting-state functional connectivity (RSFC) matrix. The RSFC matrix was generated by Tikhonov-regularized partial correlation using nets_netmats.m from FSLNets (Smith et al., 2011; Pervaiz et al., 2020). Since the correlation between two ROIs A and B is the same as the correlation between ROI B and ROI A in the RSFC matrix, the lower triangular part of the RSFC contains identical information as the upper triangular part. Therefore, only the lower triangular portion of each RSFC matrix was vectorized for each participant to serve as the individual functional connectivity features (Figure 1 top row).
2. *Individual-specific hard-parcellation Kong 2021*. Individual-specific hard-parcellations are estimated for each participant, where each vertex is only assigned to one ROI. We have previously developed a multi-session hierarchical Bayesian model (MS-HBM) of individual-specific hard-parcellation that accounted for both between-subject and within-subject variability (Kong et al., 2021a), which we will refer to as “Kong2021”. For each participant, we utilized the MS-HBM model with pre-trained group priors (https://github.com/ThomasYeoLab/CBIG/tree/master/stable_projects/brain_parcellation/Kong2022_ArealMSHBM/lib/group_priors) to generate individual-specific Kong2021 parcellations with 100 to 1000 ROIs using all rs-fMRI scans. For each parcellation resolution, the Kong 2021 parcellation was applied to all rs-fMRI scans of each participant to generate a RSFC matrix. The RSFC matrix was generated by Tikhonov-regularized partial correlation using nets_netmats.m from FSLNets (Smith et al., 2011; Pervaiz et al., 2020). The lower triangular portion of each RSFC matrix was vectorized for each participant to serve as the individual functional connectivity features (Figure 1 second row).
3. *Individual-specific soft-parcellation sICA*. Individual-specific soft-parcellations are estimated in the cerebral cortex of each participant, where each vertex could be involved in multiple ROIs. Spatial independent component analysis (ICA) is one of the most popular soft-parcellation approaches (Calhoun et al., 2001; Beckmann et al., 2005; Smith et al., 2009; Zuo et al., 2010; Nickerson et al., 2017). The individual-specific soft-parcellations could be estimated by spatial ICA (melodic tool from FSL) followed by dual regression (dual_regression tool from FSL) (Beckmann et al., 2009; Nickerson et al., 2017), which we will refer to as “sICA”. For each HCP and ABCD participant, we obtained the individual-specific sICA parcellations with 50, 100, 200, 300 components using all rs-fMRI scans. For each resolution and each participant, the RSFC matrix was generated by Tikhonov-regularized partial correlation using nets_netmats.m from FSLNets (Smith et al., 2011; Pervaiz et al., 2020). The lower triangular part of each RSFC matrix was vectorized for each participant to serve as the individual functional connectivity features (Figure 1 third row).
4. *Principal gradients (PrincipalGrad)*. Resting-state functional connectivity can be decomposed into multiple principal gradients using dimension reduction techniques, such as the non-linear diffusion embedding (Margulies et al., 2016), which we will refer to as “PrincipalGrad”. We first generated group-level principal gradients for the HCP and ABCD datasets separately using rs-fMRI scans of all participants (Margulies et al., 2016). For each dataset, we then generated the principal gradients separately for each participant using all rs-fMRI scans of this participant. The Procrustes alignment was used to align individual principal gradient maps to the group-level principal gradients (Hong et al., 2020; Vos de Wael et al., 2020). We considered different number of principal gradients. The top 1, 5, 10, 20, 40, 60, 80 or 100 principal gradients were concatenated as the individual functional connectivity features (Figure 1 fourth row).
5. *Local gradient (LocalGrad)*. The local gradient approach detects local abrupt changes in resting-state functional connectivity across the cortex (Cohen et al., 2008; Wig et al., 2014; Laumann et al., 2015; Gordon et al., 2016), which we will refer to as “LocalGrad”. For each participant in the HCP and ABCD datasets, we estimated the local gradient map using all rs-fMRI scans (Laumann et al., 2015; Gordon et al., 2016). Unlike the principal gradient approach (Margulies et al., 2016), the local gradient approach has a single gradient map, which was used as the individual functional connectivity features (Figure 1 last row).

### Behavioral data

Consistent with our previous work, we considered 58 behavioral measures from the HCP dataset (Kong et al., 2019; Li et al., 2019a; Kong et al., 2021a), and 36 behavioral measures from the ABCD dataset (Chen et al., 2022). We also included three behavioral components derived by a factor analysis from our previous work (Ooi et al., 2022). These three components for HCP dataset were interpreted to be related to cognition, life dissatisfaction, and emotional recognition, which we will refer to as “cognition”, “dissatisfaction”, and “emotion”. Across the 58 behavioral measures, the variance explained by “cognition” was 9.3%, the variance explained by “dissatisfaction” was 14.7%, and the variance explained by “emotion” was 3.8%. These three components for ABCD dataset were interpreted to be related to cognition, mental health, and personality, which we will refer to as “cognition”, “mental”, and “personality”. Across the 36 behavioral measures, the variance explained by “cognition” was 20.7%, the variance explained by “mental” was 13.2%, and the variance explained by “personality” was 7.6%.

Of the 835 HCP participants with 4 runs, only 746 have all 58 behavioral measures, who were used in the current study. Of the 1476 ABCD participants with 4 runs, all have the 36 behavioral measures, who were used in the current study.

### RSFC-Based Behavioral Prediction

Consistent with our previous work (Kong et al., 2019; Li et al., 2019a; He et al., 2020; Kong et al., 2021a), kernel ridge regression (KRR) was used to predict behavioral measures in our main analysis. Given parcellations and gradients derived from different approaches (Schaefer 2018, Kong2021, sICA, PrincipalGrad, LocalGrad), KRR performs predictions based on the similarity between functional connectivity features. Suppose *y* is the behavioral measure (e.g., fluid intelligence) and FC is the functional connectivity features of a test participant. In addition, suppose *y*_*i*_ is the behavioral measure (e.g., fluid intelligence) and FC_*i*_ is the individual-specific functional connectivity matrix of the *i*-th training participant. Then kernel regression would predict the behavior of the test participant as the weighted average of the behavioral measures of the training participants: *y* ≈ ∑_*i*∈training set_ Similarity(FC_*i*_, FC)*y*_*i*_. Here, Similarity(FC_*i*_, FC) is the Pearson’s correlation between the functional connectivity features of the *i*-th training participant and the test participant. Therefore, a training participant is weighted more if the training participant’s functional connectivity features are more similar to the test participant. For example, let’s assume there are two training participants and one test participant. The RSFC of the test participant is highly similar to one of the training participants (e.g., RSFC similarity is 0.8), but is very different from the other training participant (e.g., RSFC similarity is 0.1). Suppose the behavioral scores of these two training participants are 10 and 100, respectively. The behavioral score of the test participant will be predicted as y ≈ 10 × 0.8 + 100 × 0.1 = 18. A L2-regularization term was used in the model to reduce overfitting.

To compare the prediction performance across different parcellation and gradient approaches, we treated the resolution (i.e., number of parcels or gradients) as a hyperparameter for approaches with multiple resolutions (i.e. Schaefer2018, Kong2021, sICA, and PrincipalGrad). This hyperparameter is estimated via a nested cross-validation procedure (see below). As a separate analysis, we also compared the prediction performance across different resolutions for Schaefer2018, Kong2021, sICA, and PrincipalGrad, where the prediction was performed using RSFC features from different number of ROIs/gradients.

A nested cross-validation procedure was performed to train predictive models. In the HCP dataset, we performed 100 random replications of 20-fold nested cross-validation. Family structure was taken into account by ensuring participants from the same family were kept within the same fold and not split across folds. In the ABCD dataset, as before (Chen et al., 2022), we combined participants across the 19 imaging sites to reduce sample-size variability across sites, yielding 9 “site-clusters”. Each site-cluster comprised at least 124 participants (see Table S1).

We performed a leave-3-site-clusters out nested cross-validation. For each fold, 6 random site-clusters were used for training while the remaining 3 site-clusters were used for testing. The prediction was performed for every possible split of the site clusters, resulting in 84 replications (9 choose 3 = 84).

The resolution parameter searching ranges were [100, 200, 300, …, 1000], [100, 200, 300, …, 1000], [50, 100, 200, 300], and [1, 5, 10, 20, 40, 60, 80 or 100] for Schaefer2018, Kong2011, sICA, and PrincipalGrad respectively. The regularization parameter searching range for KRR was [0, 0.00001, 0.0001, 0.001, 0.004, 0.007, …, 1, 1.5, 2, …, 4, 5, 10, 15, 20]. The resolution parameter and the regularization parameter were estimated within the “inner-loop” of the inner-loop (nested) cross-validation procedure. The optimal parameters from the inner-loop cross-validation were then used to predict the behavioral measures in the test fold or test site clusters.

As certain behavioral measures are known to correlate with motion (Siegel et al., 2017), we regressed out age, sex, framewise displacement, and DVARS from the behavioral data before kernel ridge regression for both HCP and ABCD datasets. To prevent any information leak from the training data to test data, the regression was performed on the training data and the estimated nuisance regression coefficients were applied to the test fold.

Accuracy was measured by correlating the predicted and actual behavioral measure across all participants within the test fold (Finn et al., 2015; Kong et al., 2019; Li et al., 2019a; Kong et al., 2021a), and then averaged across test folds and replications. When comparing between approaches, a corrected resampled t-test for repeated k-fold cross-validation was performed (Bouckaert and Frank, 2004). To control for multiple comparisons, we performed a false discovery rate (FDR) (Benjamini and Hochberg, 1995) correction with q < 0.05 for all p-values reported in this paper.

To ensure our conclusions are robust across different regression approaches, we also considered linear ridge regression (LRR) as the predictive model, which has been widely used in many studies (Siegel et al., 2016; Cui et al., 2020; Rapuano et al., 2020). The regularization parameter searching range for LRR was [0.05, 0.1, 0.15, …,1].

### Code and data availability

Code for this work is freely available at the GitHub repository maintained by the Computational Brain Imaging Group (https://github.com/ThomasYeoLab/CBIG). The Schaefer2018 group-level parcellations with 100 to 1000 ROIs can be found here (https://github.com/ThomasYeoLab/CBIG/tree/master/stable_projects/brain_parcellation/Schaefer2018_LocalGlobal/Parcellations). The individual-specific Kong2021 parcellations with 100 to 1000 ROIs can be found here (https://github.com/ThomasYeoLab/Kong2022_ArealMSHBM/tree/main/Parcellations). The kernel ridge regression and linear ridge regression model used in this paper are available in this Github repository (GITHUB_LINK). Code specific to the regression models and analyses in this study can be found here (GITHUB_LINK). The HCP data are publicly available in this Github repository (GITHUB_LINK). The ABCD data are publicly available via the NIMH Data Archive (NDA) website (NDA_LINK).

## Results

### Kong2021 compared favorably with other approaches in the HCP dataset

The RSFC features of different gradient and parcellation approaches with optimal resolutions (estimated from inner-loop nested cross-validation) were used for predicting behavioral measures in the HCP dataset. For the HCP dataset, we trained a separate KRR model for each approach to predict three behavioral components (“cognition”, “dissatisfaction”, and “emotion”) and 58 behavioral measures.

Figure 2A shows the average prediction accuracies of all 58 behavioral measures, task performance measures, and self-reported measures from the HCP dataset across different gradient and parcellation approaches. Figure 2B shows the prediction accuracies of three behavioral components from the HCP dataset across different gradient and parcellation approaches.

**Figure 2.**
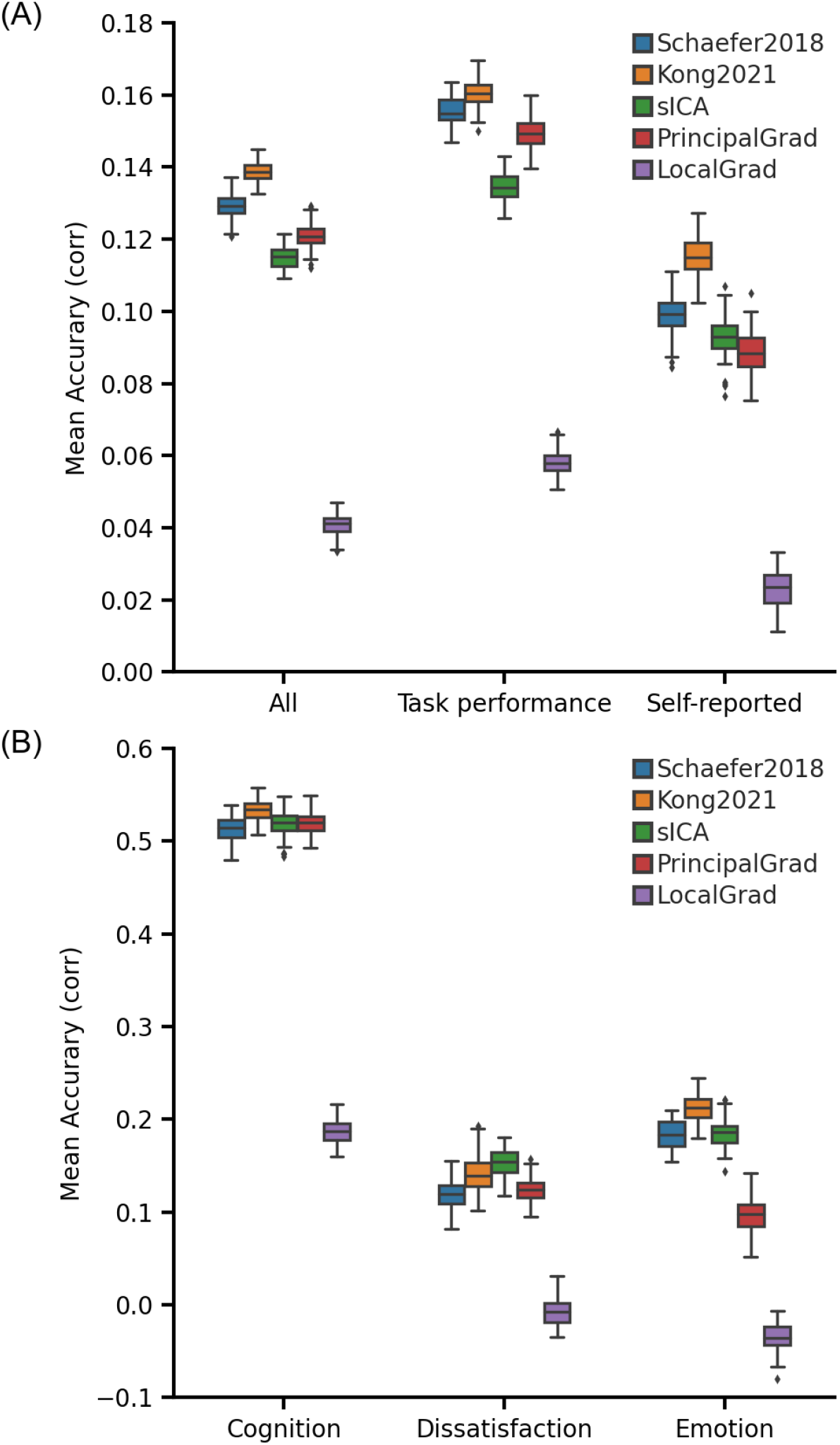
Individual-specific hard-parcellation approach Kong2021 compared favorably with other approaches for kernel ridge regression (KRR) in the HCP dataset. (A) Average prediction accuracies (Pearson’s correlation) of all 58 behavioral measures, task performance measures, and self-reported measures. (B) Prediction accuracies (Pearson’s correlation) of three behavioral components: cognition, dissatisfaction, and emotion. Boxplots utilized default Python seaborn parameters, that is, box shows median and interquartile range (IQR). Whiskers indicate 1.5 IQR. Designation of behavioral measures into “self-reported” and “task-performance” measures followed previous studies (Li et al., 2019a; Liégeois et al., 2019; Kong et al., 2021a). The RSFC features of different gradient and parcellation approaches with optimal resolutions (estimated from inner-loop nested cross-validation) were used for predicting behavioral measures. LRR results are shown in Figure S1.

To compare the prediction accuracies across different approaches, p values were computed between each pair of approaches. Figure 3 shows the p values of comparing prediction accuracies between each pair of approaches in the HCP dataset. Since there were 5 different approaches, the p values were shown as 5×5 matrices. The i-th row and j-th column of each matrix represents the p value of comparing prediction accuracies between i-th approach and j-th approach. P values that remained significant after correcting for multiple comparisons (FDR q < 0.05) were colored based on -log10(p). Therefore, bright color indicates small p values, while dark color indicates large p values. The black color indicates non-significant p values after FDR correction. The warm colors represent higher prediction accuracies of the “row” approach compared with the “column” approach. For example, the average prediction accuracy across 58 behavioral measures of Kong2021 (2nd approach) was significantly better than the sICA (4th approach). Therefore, the 2nd row and 4th column of the “All” panel (in Figure 3) is yellow, while the 4th row and 2nd column of the “All” panel (in Figure 3) is blue.

**Figure 3.**
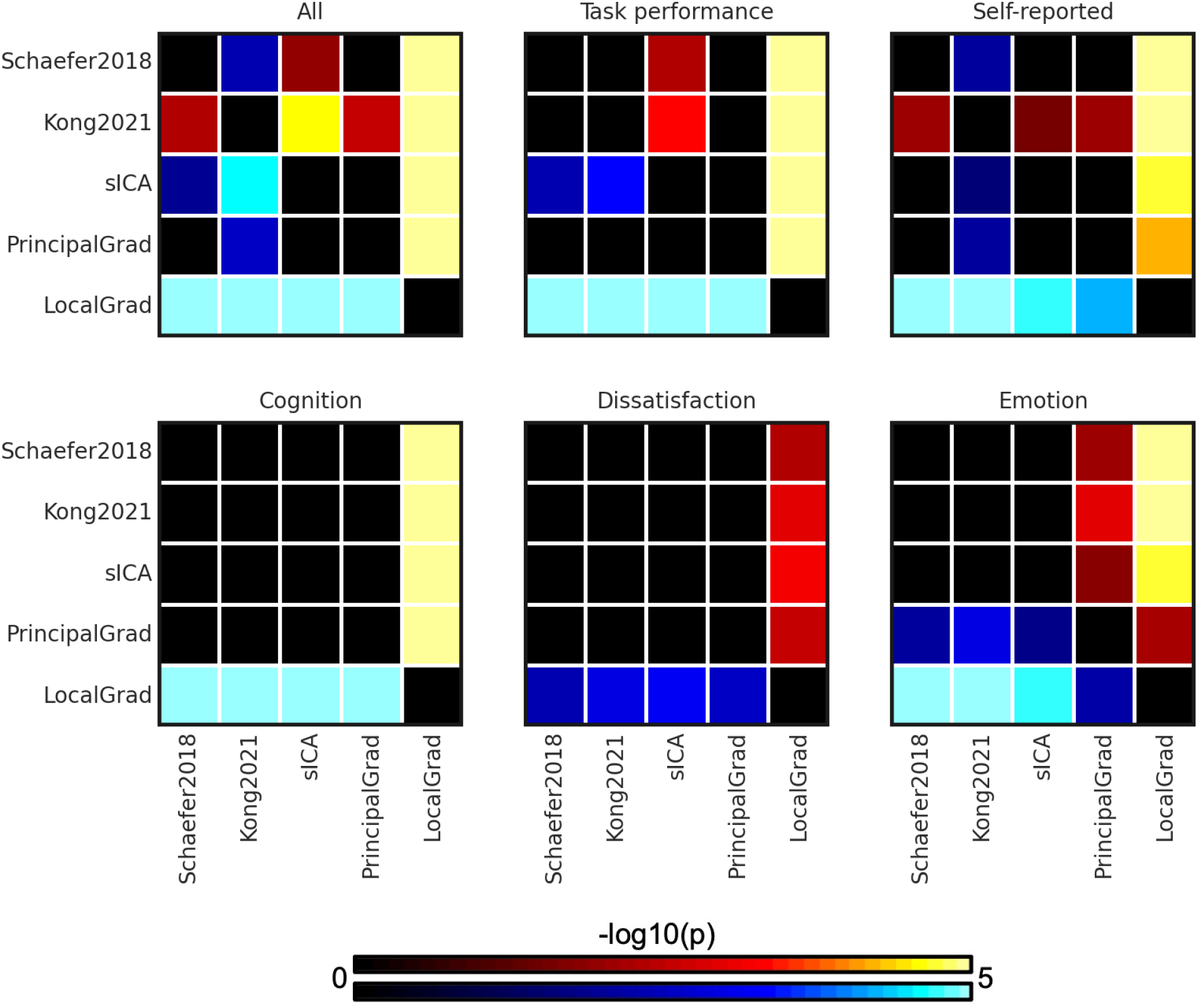
P values (-log10(p)) of comparing prediction accuracies between each pair of approaches for kernel ridge regression (KRR) in the HCP dataset. Non-black colors denote significantly different prediction performances after correcting for multiple comparisons with FDR q < 0.05. Bright colors indicate small p values, dark colors indicate large p values. For each pair of comparisons, warm colors represent higher prediction accuracies of the “row” approach than the “column” approach. Individual-specific hard-parcellation approach Kong2021 compared favorably with the other approaches, as can be seen from warm colors along the rows corresponding to Kong2021. LRR results are shown in Figure S2.

The individual-specific hard-parcellation approach Kong2021 compared favorably with the other approaches, as can be seen from warm colors along the rows corresponding to Kong2021 in Figure 3. This is especially the case for average prediction accuracies across all 58 behavioral measures (p=4.4e-3, p=2.9e-5, p=2.4e-3, and p=5.3e-25 with respect to Schaefer2018, sICA, PrincipalGrad, and LocalGrad, respectively) and self-reported measures (p=7.7e-3, p=2.9e-2, p=7.6e-3, and p=1.5e-8 with respect to Schaefer2018, sICA, PrincipalGrad, and LocalGrad, respectively). The principal gradient approach PrincipalGrad, sICA, and Schaefer2018 performed similarly. The local gradient approach LocalGrad performed the worst, as can be seen by cool colors along the rows corresponding to LocalGrad in Figure 3. Similar results were obtained with LRR (Figures S1 and S2).

We repeated the comparison using full correlation RSFC instead of partial correlation RSFC for Schaefer2018, Kong2021, and sICA. Consistent with previous work (Dadi et al., 2019; Pervaiz et al., 2020; Farahibozorg et al., 2021), we found that full correlation RSFC performed worse than partial correlation RSFC (Figure S3). Because of the lower prediction performance for full correlation RSFC, the principal gradient approach PrincipalGrad achieved statistically better performance than full correlation RSFC for certain behavioral measures (e.g., task performance), while achieving similar results in other behavioral measures (Figures S4-S7).

### Parcellation and principal gradients exhibit similar performance in the ABCD dataset

The RSFC features of different gradient and parcellation approaches with optimal resolutions (estimated from inner-loop nested cross-validation) were used for predicting behavioral measures in the ABCD dataset. For the ABCD dataset, we trained a separate KRR model for each approach to predict three behavioral components (“cognition”, “mental”, and “personality”) and 36 behavioral measures. For both datasets, we categorized the behavioral measures into “task performance” and “self-reported” measures.

Figure 4A shows the average prediction accuracies of all 36 behavioral measures, task performance measures, and self-reported measures from the ABCD dataset across different gradient and parcellation approaches. Figure 4B shows the prediction accuracies of three behavioral components from the ABCD dataset across different gradient and parcellation approaches.

**Figure 4.**
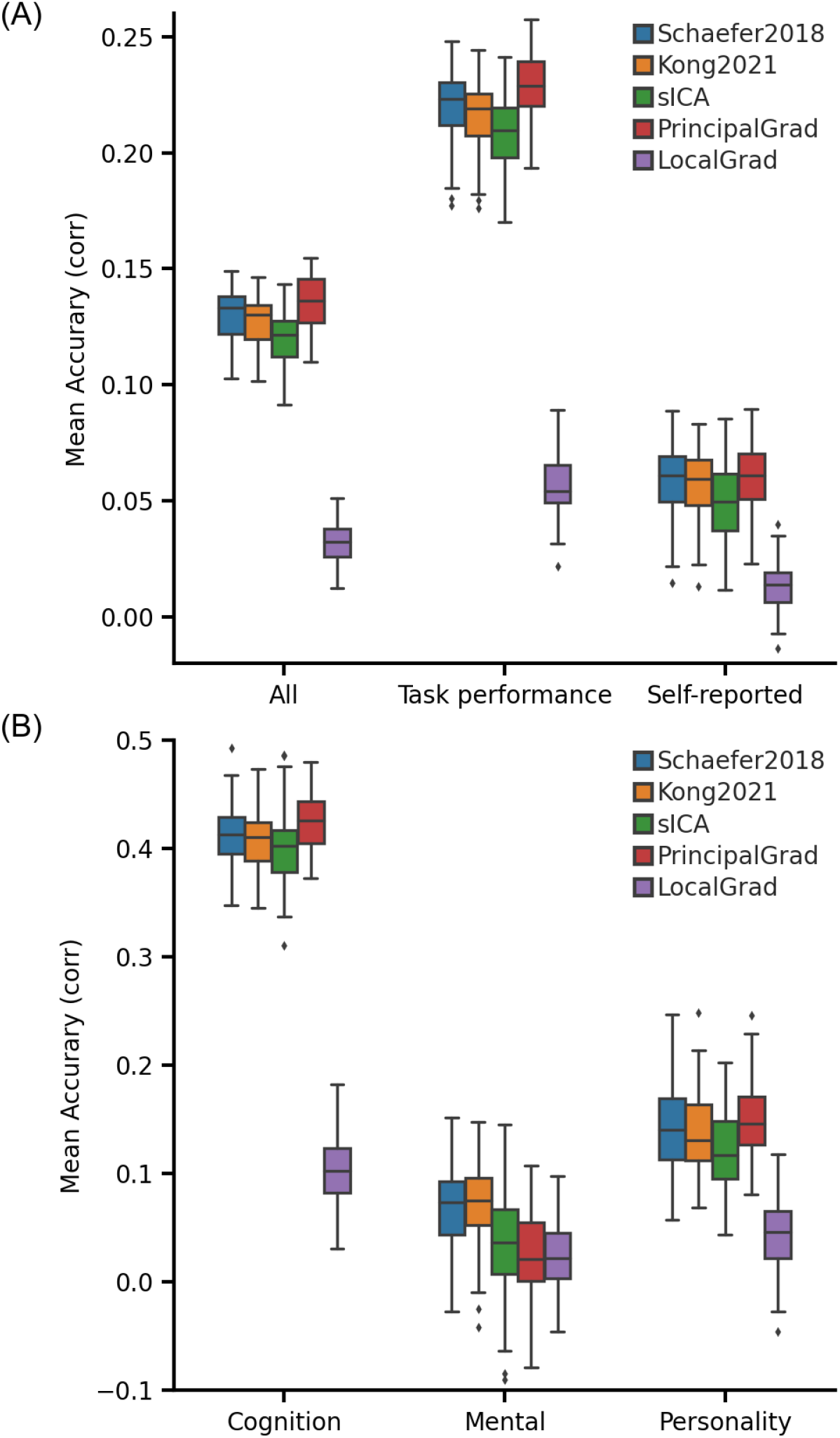
Principal gradient approach achieves comparable behavioral prediction performance as parcellation approaches for kernel ridge regression (KRR) in the ABCD dataset. (A) Average prediction accuracies (Pearson’s correlation) of all 36 behavioral measures, task performance measures, and self-reported measures. (B) Prediction accuracies (Pearson’s correlation) of three behavioral components: cognition, mental health, and personality. Boxplots utilized default Python seaborn parameters, that is, box shows median and interquartile range (IQR). Whiskers indicate 1.5 IQR. Designation of behavioral measures into “self-reported” and “task-performance” measures followed previous studies (Li et al., 2019a; Liégeois et al., 2019; Kong et al., 2021a). The RSFC features of different gradient and parcellation approaches with optimal resolutions (estimated from inner-loop nested cross-validation) were used for predicting behavioral measures. LRR results are shown in Figure S8. The principal gradient approach PrincipalGrad was numerically the best for most cases, but there was largely no statistical difference among the approaches.

**Figure 3.**
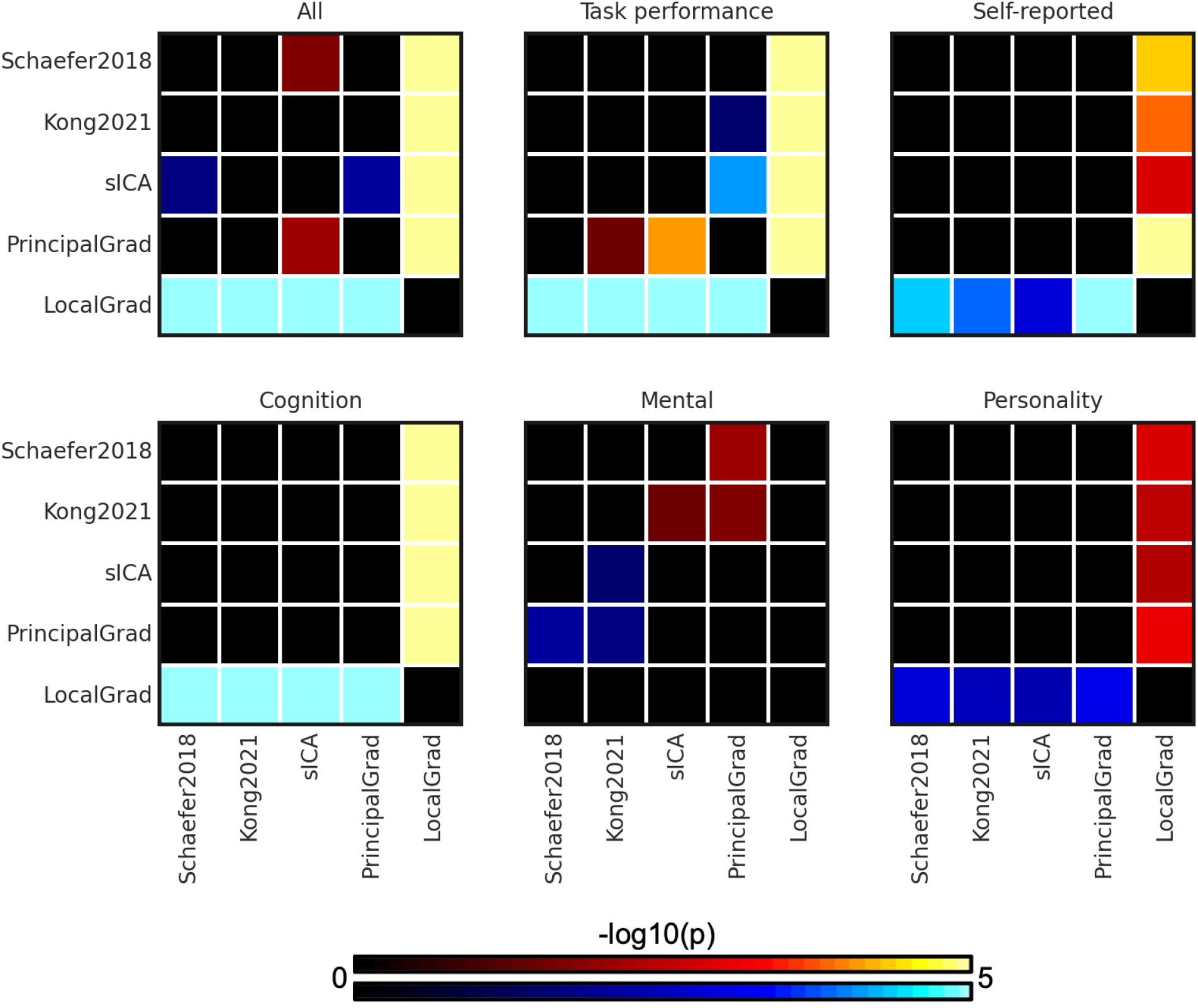
P values (-log10(p)) of comparing prediction accuracies between each pair of approaches for kernel ridge regression (KRR) in the ABCD dataset. Non-black colors denote significantly different prediction performances after correcting for multiple comparisons with FDR q < 0.05. Bright colors indicate small p values, dark colors indicate large p values. For each pair of comparisons, warm colors represent higher prediction accuracies of the “row” approach than the “column” approach. There was no statistical difference among most approaches. LRR results are shown in Figure S9.

In the ABCD dataset, the principal gradient approach PrincipalGrad was numerically the best for most cases, but there was largely no statistical difference among the approaches (Figure 5). More specifically, PrincipalGrad was significantly better than Kong2021, sICA and LocalGrad in the case of the average prediction accuracies across task performance measures (p=3.3e-2, p=7.6e-5, and p=5.8e-29 with respect to Kong2021, sICA, and LocalGrad, respectively), while Kong2021 was significantly better than sICA and PrincipalGrad in the case of the mental health component (p=3.7e-2, p=2.0e-2 with respect to sICA and PrincipalGrad respectively). Similar results were obtained with LRR (Figures S8 and S9).

**Figure 4.**
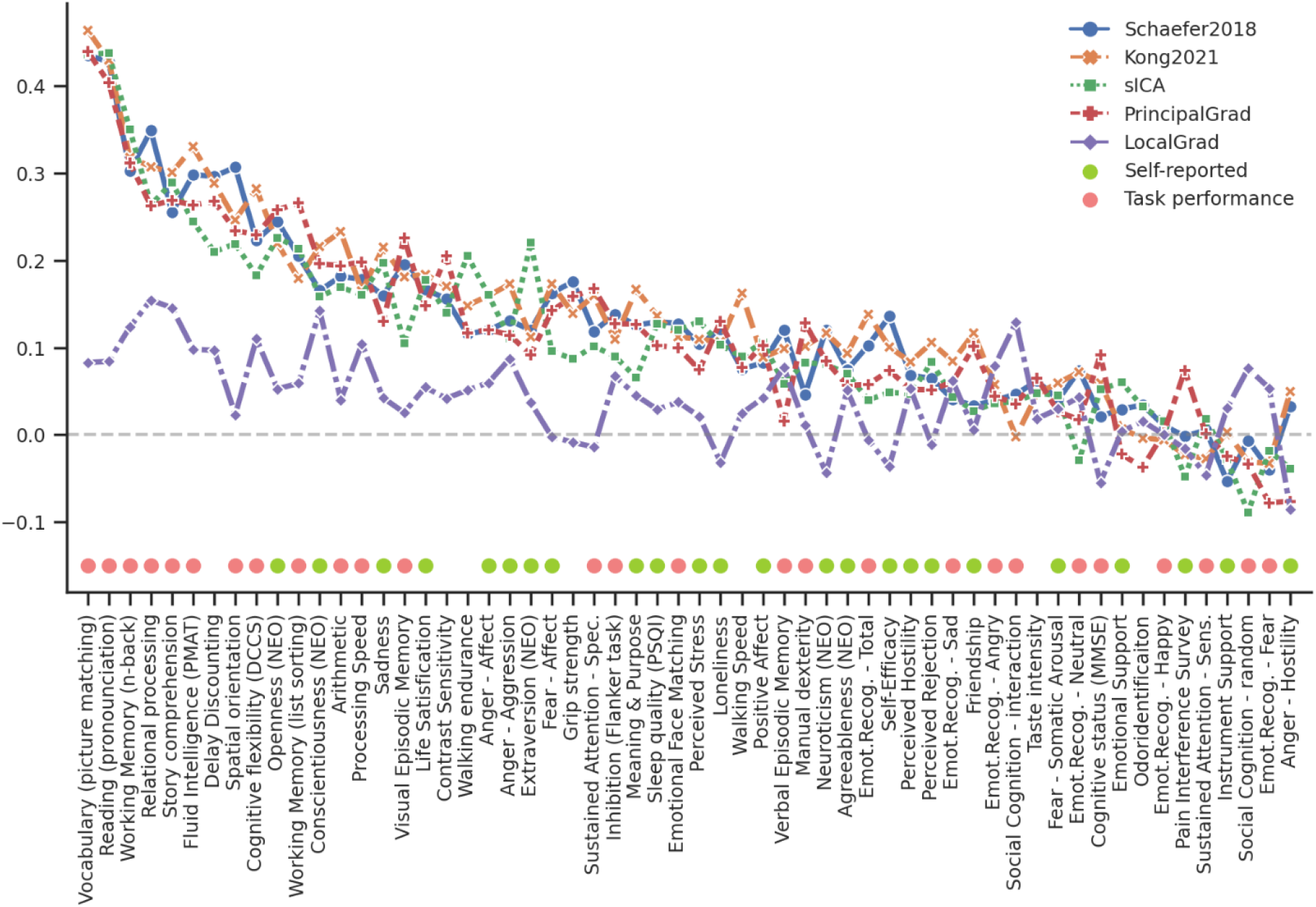
Task performance measures were predicted better than self-reported measures across different gradient and parcellation approaches with optimized resolutions for KRR in the HCP dataset. 58 behavioral measures were ordered based on average prediction accuracies across Schaefer2018, Kong2021, sICA, PrincipalGrad, and LocalGrad. Pink circles indicate task performance measures. Green circles indicate self-reported measures. Boxplots utilized default Python seaborn parameters, that is, box shows median and interquartile range (IQR). Whiskers indicate 1.5 IQR. Designation of behavioral measures into “self-reported” and “task-performance” measures followed previous studies (Li et al., 2019a; Liégeois et al., 2019; Kong et al., 2021a). LRR results are shown in Figure S15.

**Figure 5.**
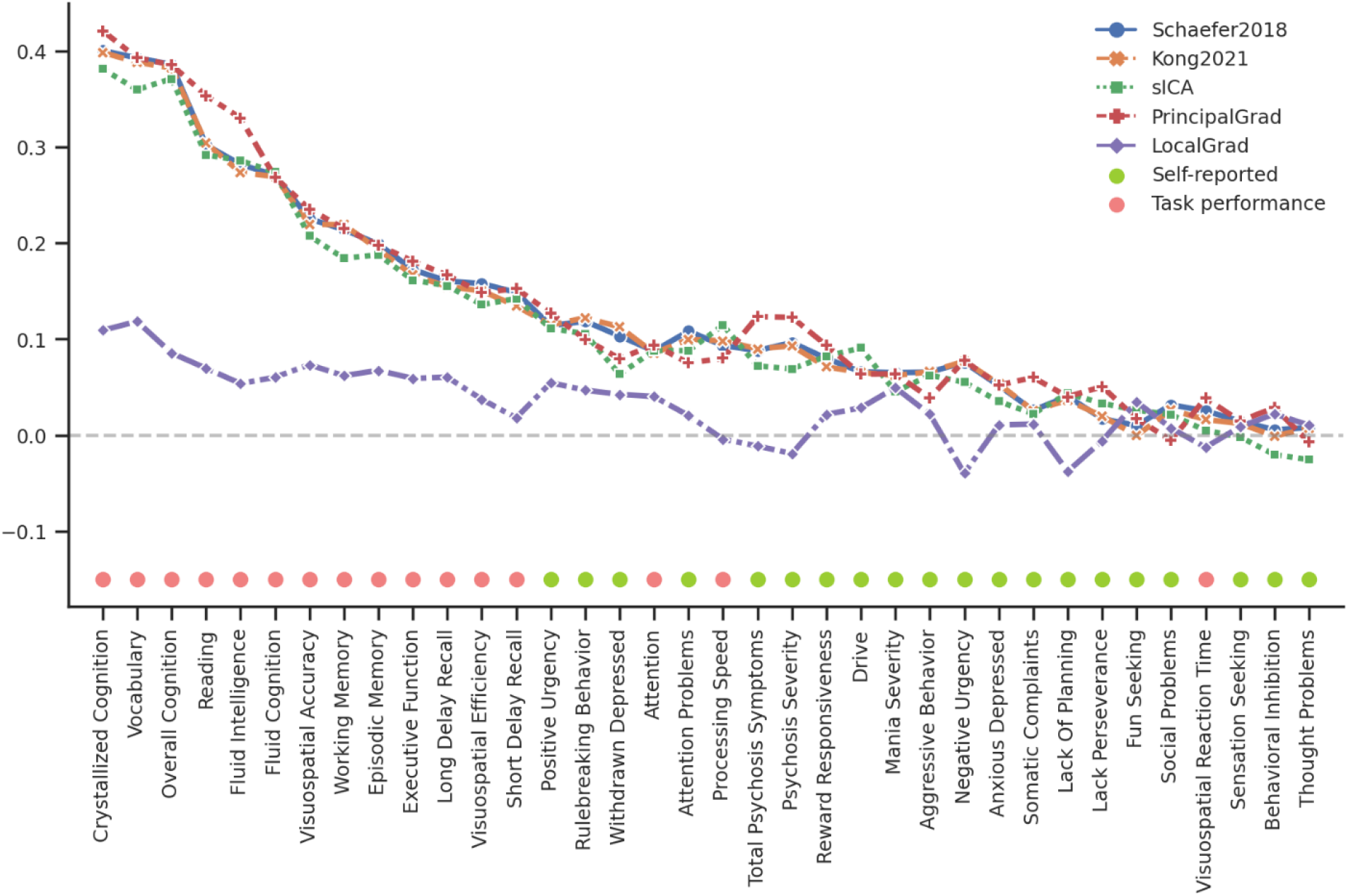
Task performance measures were predicted better than self-reported measures across different gradient and parcellation approaches with optimized resolutions for KRR in the ABCD dataset. 36 behavioral measures were ordered based on average prediction accuracies across Schaefer2018, Kong2021, sICA, PrincipalGrad, and LocalGrad. Pink circles indicate task performance measures. Green circles indicate self-reported measures. Boxplots utilized default Python seaborn parameters, that is, box shows median and interquartile range (IQR). Whiskers indicate 1.5 IQR. Designation of behavioral measures into “self-reported” and “task-performance” measures based on ABCD behavioral measures description (Li et al., 2019a; Liégeois et al., 2019; Kong et al., 2021a). LRR results are shown in Figure S16.

We repeated the comparison using full correlation RSFC instead of partial correlation RSFC for Schaefer2018, Kong2021, and sICA. The prediction performance of full correlation RSFC was numerically (but not significantly) worse than using partial correlation RSFC for most cases in the ABCD dataset (Figure S10). Because of the lower prediction performance for full correlation RSFC, the principal gradient approach now achieved statistically better performance than full correlation RSFC for certain behavioral measures (e.g., task performance and cognition), while achieving similar results in other behavioral measures (Figures S11-S14).

### Task performance measures are more predictable than self-reported measures for all approaches

To explore which behavioral measures can be consistently predicted well regardless of gradient and parcellation approaches, we ordered the behavioral measures based on averaged prediction accuracies (Pearson’s correlation) across different approaches with the optimized resolutions for KRR in the HCP (Figure 6) and ABCD (Figure 7) datasets. Our previous studies have suggested that self-reported and task performance measures might be differentially predicted under different conditions (Li et al., 2019a; Liégeois et al., 2019; Kong et al., 2021a). For example, the task performance measures were more predictable than self-reported measures based on functional connectivity of hard-parcellation approaches (Kong et al., 2021a).

**Figure 6.**
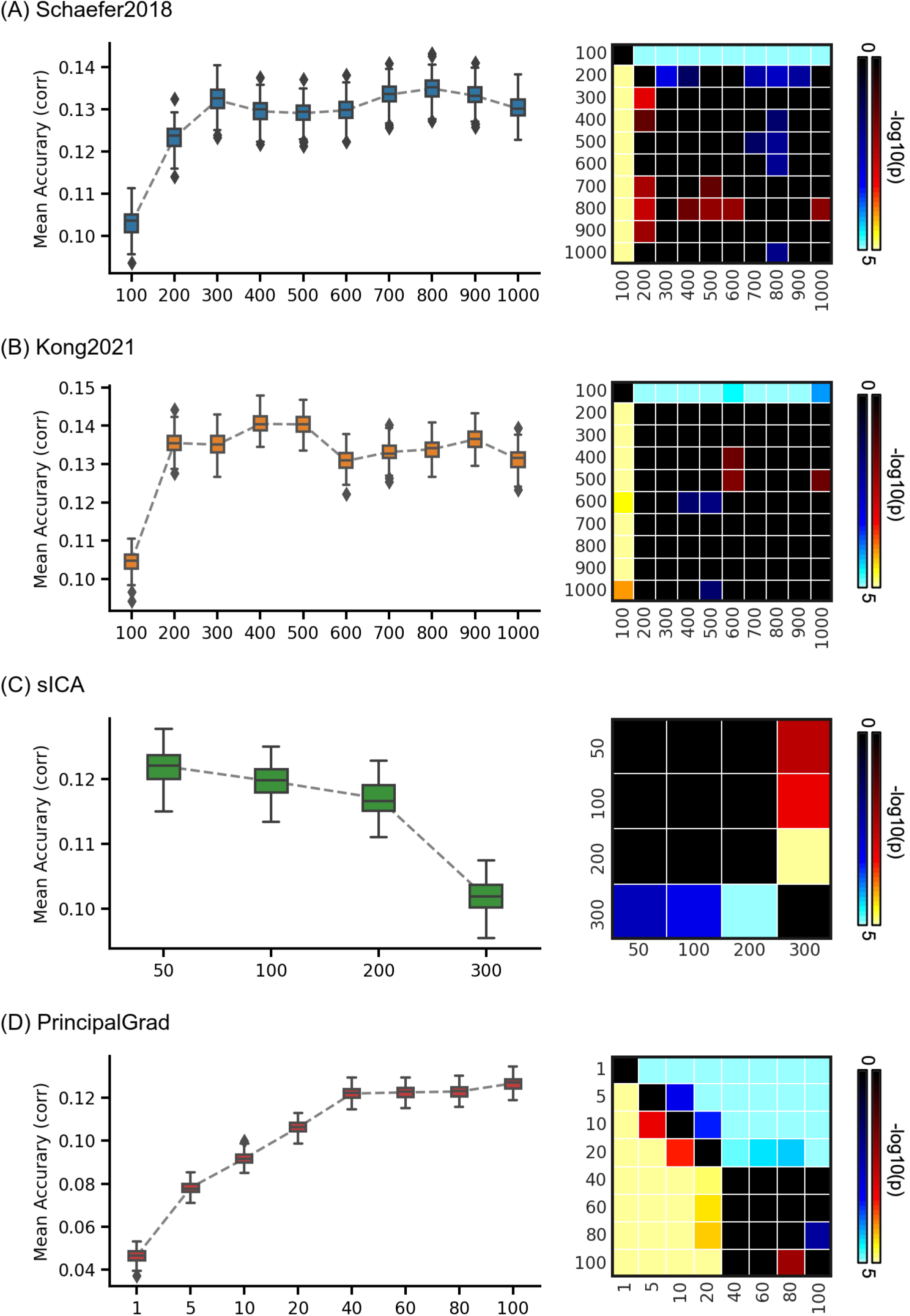
Average prediction accuracies (Pearson’s correlation) of all 58 behavioral measures vary across resolutions for gradient and parcellation approaches using KRR in the HCP dataset. (A) Prediction accuracies and p values of the hard-parcellation Schaefer2018 with 100 to 1000 ROIs. (B) Prediction accuracies and p values of the hard-parcellation Kong2021 with 100 to 1000 ROIs. (C) Prediction accuracies and p values of the soft-parcellation sICA with 50 to 300 components. (D) Prediction accuracies and p values of the principal gradient PrincipalGrad with 1 to 100 gradients. Boxplots utilized default Python seaborn parameters, that is, box shows median and interquartile range (IQR). Whiskers indicate 1.5 IQR. P values (-log10(p)) were computed between prediction accuracies of each pair of resolutions. Non-black colors denote significantly different prediction performances after correcting for multiple comparisons with FDR q < 0.05. Bright colors indicate small p values, dark colors indicate large p values. For each pair of comparisons, warm colors represent higher prediction accuracies of the “row” resolution than the “column” resolution.

**Figure 7.**
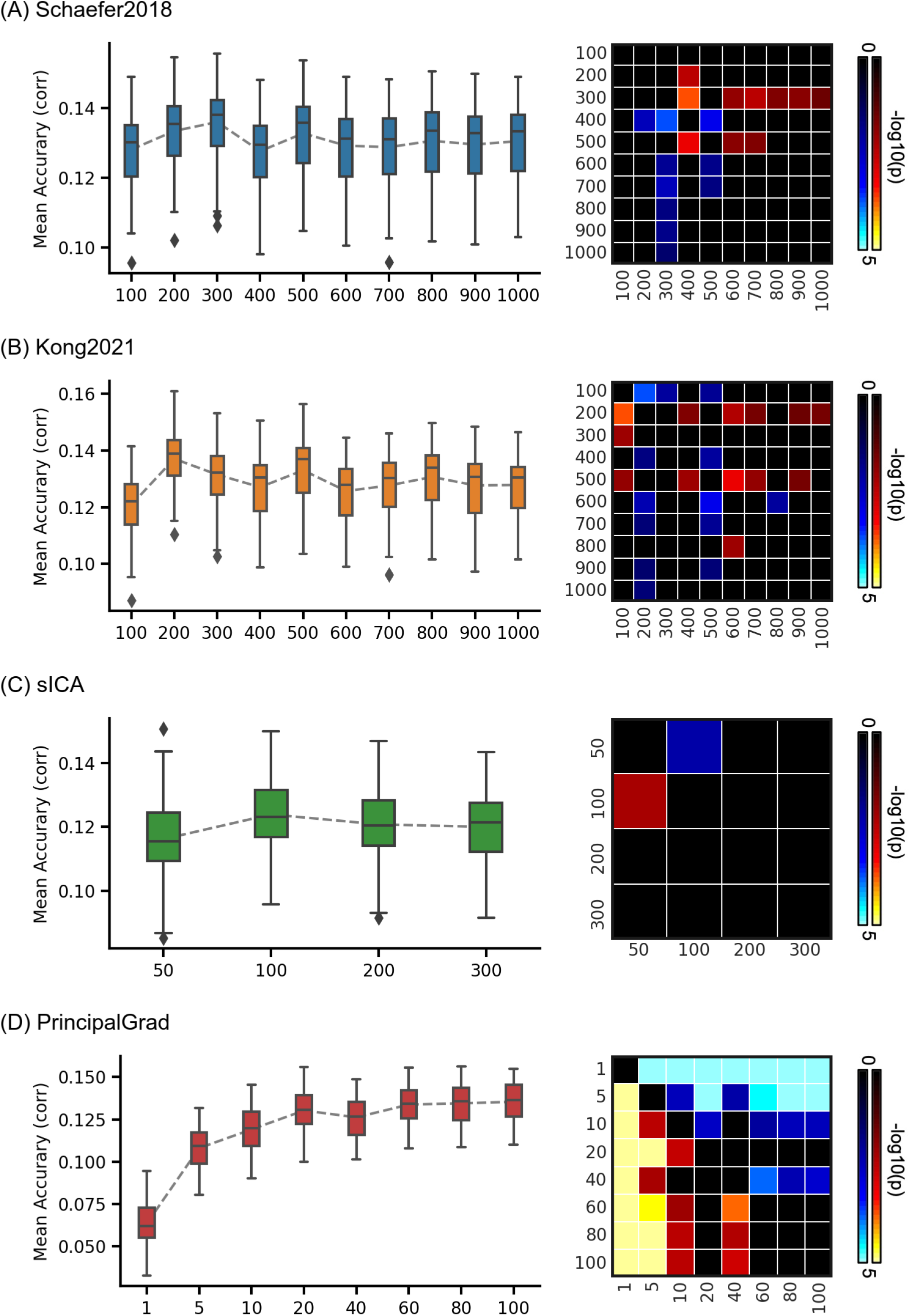
Average prediction accuracies (Pearson’s correlation) of all 36 behavioral measures vary across resolutions for gradient and parcellation approaches using KRR in the ABCD dataset. (A) Prediction accuracies and p values of the hard-parcellation Schaefer2018 with 100 to 1000 ROIs. (B) Prediction accuracies and p values of the hard-parcellation Kong2021 with 100 to 1000 ROIs. (C) Prediction accuracies and p values of the soft-parcellation sICA with 50 to 300 components. (D) Prediction accuracies and p values of the principal gradient PrincipalGrad with 1 to 100 gradients. Boxplots utilized default Python seaborn parameters, that is, box shows median and interquartile range (IQR). Whiskers indicate 1.5 IQR. P values (-log10(p)) were computed between prediction accuracies of each pair of resolutions. Non-black colors denote significantly different prediction performances after correcting for multiple comparisons with FDR q < 0.05. Bright colors indicate small p values, dark colors indicate large p values. For each pair of comparisons, warm colors represent higher prediction accuracies of the “row” resolution than the “column” resolution.

In the HCP dataset, we found the average prediction accuracies of task performance measures were r = 0.156 ± 0.004 (mean ± std), r = 0.160 ± 0.004, r = 0.135 ± 0.004, r = 0.149 ± 0.004, and r = 0.058 ± 0.003 for Schaefer2018, Kong2021, sICA, PrincipalGrad, and LocalGrad, respectively (Figure 2A), while the prediction accuracies of self-reported measures were r = 0.099 ± 0.005, r = 0.115 ± 0.005, r = 0.093 ± 0.005, r = 0.089 ± 0.006, and r = 0.023 ± 0.005 for Schaefer2018, Kong2021, sICA, PrincipalGrad, and LocalGrad, respectively (Figure 2A).

In the ABCD dataset, we found the average prediction accuracies of task performance measures were r = 0.220 ± 0.016 (mean ± std), r = 0.216 ± 0.015, r = 0.209 ± 0.016, r = 0.229 ± 0.014, and r = 0.056 ± 0.013 for Schaefer2018, Kong2021, sICA, PrincipalGrad, and LocalGrad, respectively (Figure 4A), while the prediction accuracies of self-reported measures were r = 0.059 ± 0.015, r = 0.058 ± 0.015, r = 0.049 ± 0.016, r = 0.061 ± 0.0140, and r = 0.014 ± 0.010 for Schaefer2018, Kong2021, sICA, PrincipalGrad, and LocalGrad, respectively (Figure 4A).

These results suggested that on average, task performance measures were more predictable than self-reported measures across all gradient and parcellation approaches (p = 4.0e-4, p = 2.6e-3, p = 7.2e-3, p = 2.6e-4, and p = 3.5e-2 for Schaefer2018, Kong2021, sICA, PrincipalGrad, and LocalGrad, respectively). Similar results were obtained with LRR (Figures S15 and S16). P values remained significant after correcting for multiple comparisons with FDR q < 0.05.

### Prediction performances vary across resolutions for both gradient and parcellation approaches

To explore the impact of the number of gradients and parcels, we performed behavioral prediction for each approach using different resolutions in the HCP and ABCD datasets. The left column of Figure 8 and 9 show the KRR prediction accuracies (Pearson’s correlation) of the average prediction accuracies of all behavioral measures for Schaefer2018, Kong2021, sICA, and PrincipalGrad in the HCP and ABCD datasets with different resolutions. The local gradient approach LocalGrad was not included here because this approach did not have different resolutions.

To compare the prediction accuracies across different resolutions for each approach, p values were computed between each pair of resolutions. The right column of Figure 8 and 9 show the p values of comparing prediction accuracies between each pair of resolutions in the HCP and ABCD datasets. If there were K different resolutions for an approach, the p values of this approach were shown as K×K matrices. The i-th row and j-th column of each matrix represents the p value of comparing prediction accuracies between i-th resolution and j-th resolution. P values remained significant after correcting for multiple comparisons with FDR q < 0.05 were colored based on -log10(p). The bright colors indicate small p values, while the dark colors indicate large p values. The warm colors represent higher prediction accuracies of the “row” resolution than the “column” column resolution. For example, the average prediction accuracy across 58 behavioral measures of Schaefer2018 100 ROIs (1st resolution) was significantly worse than Schaefer2018 400 ROIs (4th resolution). Therefore, the 1st row and 4th column of the Figure 8A p value matrix is blue color, while the 4th row and 1st column of that p value matrix is yellow.

In the HCP dataset, the hard-parcellation approaches Schaefer2018 and Kong2021with low resolutions generally predicted behavioral measures worse than high resolutions, especially in the case of 100 ROIs (Figures 8A and 8B). The prediction accuracies plateaued around 300 ROIs for Schaefer2018 and 200 ROIs for Kong2021. Compared with other resolutions, Schaefer2018 and Kong2021 with 100 ROIs yielded significantly worse prediction accuracies of all 58 behavioral measures with p < 2.6e-7 and p < 8.3e-5, respectively. Similarly, the principal gradient approach PrincipalGrad with low resolutions also predicted behavioral measures worse than high resolutions (Figure 8D). The prediction accuracies kept increasing and plateaued around 40 gradients. Compared with using more than 40 gradients, PrincipalGrad with less than 40 gradients yielded significantly worse prediction accuracies of all 58 behavioral measures with p < 5.2e-5. By contrast, the soft-parcellation approach sICA with high resolutions predicted behavioral measures worse than low resolutions, especially in the case of 300 components (Figure 8C). Compared with other resolutions, sICA with 300 components yielded significantly worse prediction accuracies with p < 3.8e-3.

Intriguingly, in the ABCD dataset, the prediction accuracies of all parcellation approaches Schaefer2018, Kong2021, and sICA exhibited no obvious difference across resolutions (Figure 9A to 9C). The principal gradient approach PrincipalGrad with low resolutions generally predicted behavioral measures worse than high resolutions. The prediction accuracies plateaued around 60 gradients (Figure 9D). Compared with less than 40 gradients, PrincipalGrad with more than 60 gradients yielded significantly higher prediction accuracies with p < 9.8e-3.

Similar results were obtained with LRR (Figures S17 and S18). Prediction results of task performance measures, self-reported measures, and three behavioral components are shown in Figures S9 to S28, yielding similar conclusions. Figure S39 to S42 show the prediction results across different resolutions for all approaches in the same plot.

## Discussion

### Overview

In this manuscript, we compared different parcellation and gradient approaches for RSFC prediction of behavioral measures in two different datasets. Individual-specific hard-parcellation Kong2021 compared favorably with other approaches in the HCP dataset, while principal gradient and parcellation approaches performed similarly in the ABCD dataset. We found that for all parcellation and gradient approaches, task performance measures were more predictable than self-reported measures. We showed that prediction performances varied across resolutions for all gradient and parcellation approaches. Furthermore, RSFC principal gradients at sufficiently high resolution (e.g., more than 40 or 60 gradients) exhibited similar behavioral prediction performance as parcellation-based RSFC. These findings were replicated in both HCP and ABCD datasets using two prediction models KRR and LRR.

### Functional connectivity behavioral prediction using parcellation versus gradients

There has been great interest in functional connectivity prediction of behavioral measures. While most previous studies utilized RSFC from hard- or soft-parcellations, few studies have focused on gradient techniques for behavioral prediction. One recent study showed that principal gradients were behaviorally meaningful (Hong et al., 2020). Hong and colleagues further compared the prediction performance between principal gradients and RSFC from the Schaefer2018 group-level hard-parcellation with 1000 ROIs. They found that 100 principal gradients outperformed Schaefer2018 1000-ROI hard-parcellation in predicting cognitive factor score in HCP dataset.

In our study, instead of only focusing one specific resolution, we considered multiple resolutions for each approach and optimized the resolution as a hyperparameter in the prediction models. We compared the prediction performance between parcellation and gradient approaches using the same prediction framework with two different prediction models. We performed prediction for a wide range of behavioral measures across different domains and three behavioral components derived from a factor analysis. The prediction analyses were done in the HCP healthy young adult dataset, and the ABCD healthy children dataset.

Similarly, we also found that principal gradients could predict behavioral measures as well as RSFC from parcellations in HCP dataset using both KRR and LRR. However, unlike Hong and colleagues (Hong et al., 2020), we found that principal gradients achieved similar level of prediction accuracy as the Schaefer2018 group-level hard-parcellation in HCP dataset. Furthermore, the individual-specific hard-parcellation Kong2021 achieved the best prediction results.

More specifically, Hong and colleagues showed that combining 100 principal gradients was able to predict cognitive factor score with a r = 0.405, while RSFC from Schaefer2018 1000- ROI group-level hard-parcellation only achieved an accuracy of r = 0.181. In our study, we found that principal gradients could predict cognition component score with a r = 0.520 ± 0.011 using KRR and r = 0.487 ± 0.011 using LRR, while RSFC from Schaefer2018 group-level hard-parcellation could achieve an accuracy of r = 0.513 ± 0.013 using KRR and r = 0.514 ± 0.008 using LRR. Therefore, our principal gradient prediction performance was similar to our parcellation-based RSFC prediction performance. We replicated similar results in a children dataset (ABCD).

The main reason for this discrepancy might be due to different prediction models being utilized for principal gradient and parcellation approaches in Hong and colleagues (2020). In their behavioral prediction framework, they utilized the canonical correlation analysis (CCA) for principal gradients, but utilized the connectome-based predictive modeling approach (Shen et al., 2017) for Schaefer2018. Since the choice of prediction model in the behavioral prediction framework affects the prediction performance (Cui and Gong, 2018; Dadi et al., 2019; Pervaiz et al., 2020), the prediction results generated by two different prediction models were not comparable between approaches. In Hong and colleagues (2020), it is unclear whether the superior prediction performance of principal gradients compared to Schaefer2018 is due to different representations (i.e., the principal gradients representing the brain better than the Schaefer2018 parcellation) or different prediction models (i.e., CCA outperforming the connectome-based predictive modeling approach).

Another possible source of this discrepancy might be differences between the cognitive factor score from Hong and colleagues (Hong et al., 2020) and the cognitive component score from the current study. Hong and colleagues used an exploratory factor analysis (EFA) to extract a cognitive factor score from 65 HCP raw behavioral measures, while we used a principal component analysis to derive a cognitive component score from 58 HCP raw behavioral measures. However, the cognitive factor score from Hong and colleagues and the cognition component score from the current study both exhibited strong loading on similar cognitive performance scores such as fluid intelligence (PMAT), visual episodic memory, reading, reading (pronunciation), vocabulary (picture matching), and spatial orientation. Therefore, this might not be the main reason.

Interestingly, the local gradient approach performed much worse than parcellation approaches and principal gradients. The local gradient approach has been widely used as a tool to derive hard-parcellation (Laumann et al., 2015; Gordon et al., 2016). Specifically, an edge detection approach (i.e. watershed algorithm) was applied on the gradient map to generate the binarized boundary map, which could be used to define non-overlapping ROIs (Laumann et al., 2015; Gordon et al., 2016). Recent studies (Bijsterbosch et al., 2018; Kong et al., 2019) have suggested that network topography is behaviorally meaningful, so we hypothesize that local gradient maps can also be used to predict behavior. Therefore, instead of using the hard-parcellation from the local gradient approach to predict behavioral measures, we used the individual-specific gradient maps. However, a single gradient map might lose too much information compared with the RSFC from the local gradient hard-parcellation, yielding poor prediction performance. The local gradient map was especially suited for delineating brain regions because detecting abrupt changes in RSFC is somewhat similar to delineate histological boundaries of cortical areas (Cohen et al., 2008; Buckner and Yeo, 2014; Wig et al., 2014). In fact, the local gradient map was partially used in Schaefer2018 and Kong2021 to derive better hard-parcellations. Overall, this suggests that the local gradients might be helpful for deriving cortical parcellations but might not contain much behaviorally-relevant information.

### Prediction of task performance measures is better than self-reported measures

Our results suggested that the task performance measures were more predictable than self-reported measures by RSFC for all parcellation and gradient approaches in both HCP and ABCD datasets (Figures 6,7, S15 and S16). This distinction between task performance and self-reported measures echoed well with previous investigations of RSFC–behavior relationships. It has been shown that RSFC could predict cognition and task performance measures better than self-reported measures (Dubois et al., 2018a; Li et al., 2019a; Kong et al., 2021a). Dynamic functional connectivity is also more strongly associated with task performance measures than self-reported measures (Vidaurre et al., 2017; Liégeois et al., 2019; Ikeda et al., 2022). Furthermore, utilizing functional connectivity from task fMRI rather resting-state fMRI has been shown to improve the prediction of cognition more than personality and mental health (Chen et al., 2022).

One possible reason for better prediction accuracies in task performance measures might be a result of the subjective nature of self-reported measures. For example, the self-reported personality measures NEO-FFI could be influenced by an individual’s insight, impression management, and reference group effects (Dubois et al., 2018b), leading to unreliable estimate of personality. We might be able to predict self-reported measures better with more accurate estimates of personality, emotion and mental health.

### Impact of resolutions in connectivity prediction of behavioral measures

Previous studies have established that the optimal resolution for behavioral prediction varied across behavioral measures using RSFC from different soft- and hard-parcellation approaches (Dadi et al., 2019, 2020). Within a reasonable range, the impact of resolutions in prediction accuracy was small (Dadi et al., 2019). Pervaiz and colleagues also explored the impact of sICA resolution for predicting fluid intelligence(Pervaiz et al., 2020). They found that increasing dimensionality of sICA could lead to an increase in prediction accuracy. Furthermore, sICA outperformed the group-level hard-parcellation approach Schaefer2018 in predicting fluid intelligence (Pervaiz et al., 2020). One recent study compared cognition prediction accuracies of different resolutions of a new soft-parcellation approach (Farahibozorg et al., 2021). They focused on high-resolution soft-parcellations (i.e. 100, 150, 200 components) and found the prediction performance was generally similar across resolutions (Farahibozorg et al., 2021).

Consistent with previous studies, we found the optimal resolution in predicting behavioral measures varied across behavioral phenotypes for each parcellation approach. Within a range of high resolutions, the impact of resolutions in prediction performance were relatively small for hard-parcellation approaches. We also found that increasing resolutions of different parcellation approaches might not yield better prediction performance. Specifically, the soft-parcellation approach sICA tended to have relatively lower prediction accuracy with very high resolution in the HCP dataset. For example, the average prediction accuracy of all 58 behavioral measures of HCP using sICA 200 components were significantly worse than lower resolutions 50, 100, and 150 (Figure 8C).

Intriguingly, we found that the prediction accuracies of soft- and hard-parcellation approaches had no obvious difference across resolutions in the ABCD children dataset. One possible reason for this might be due to discrepancy in brain organization between healthy young adults and young children. Specifically, the Schaefer2018 group-level hard-parcellations were derived by healthy young adults, which might not be optimal for representing RSFC of young children.

While there have been several studies focusing on the impact of different resolutions for soft- and hard-parcellation approaches in RSFC prediction of behavioral measures (Dadi et al., 2019; Pervaiz et al., 2020; Farahibozorg et al., 2021), few studies have looked into the gradient approach. A recent study (Hong et al., 2020) compared the prediction performances between using a single principal gradient versus combining 100 gradients. In our study, we considered a wide range of resolutions for principal gradients. With increased number of principal gradients, we found that the prediction performance increased in both HCP and ABCD dataset. Most studies have focused mainly on the first or first several principal gradients (Margulies et al., 2016; Paquola et al., 2019; Wang et al., 2019; Tian et al., 2020; Kong et al., 2021b), since these gradients captured the most variance in RSFC (Huntenburg et al., 2018). However, the first principal gradient alone predicted behavioral measures very poorly in both HCP and ABCD datasets. In fact, the prediction accuracy of principal gradients plateaued only after more than 40 gradients in the HCP dataset and 60 gradients in the ABCD dataset.

We note that the optimal number of principal gradients might depend on the goal of a particular study. The first few gradients are relatively stable features of brain organization that might not vary significantly across individuals. As such, most literature interested in organization of the brain at the group-level mainly focused on the first few principal gradients. However, the less widely studied higher order gradients captured more idiosyncrasies across participants. Principal gradients studies focusing on individual differences in human behaviors would need at least 40 to 60 gradients to not lose significant behaviorally relevant information. On the other hand, for other studies, a smaller number of gradients might suffice.

### Limitations

There are a wide range of options that exist for parcellation approaches (Glasser2016, Shen2013, Gordon2016, Farahibozorg2021). The gradients could also be derived using other manifold learning algorithms such as principal component analysis (PCA; Hong et al. 2020) and Laplacian eigenmaps (LE) (Haak et al., 2018; Tian et al., 2020). While our behavioral prediction framework is applicable to other parcellation and gradient approaches, we focused on three representative parcellation approaches (Schaefer2018, Kong2021, and sICA) and two gradient approaches (PrincipalGrad, LocalGrad) in this paper. Previous papers have demonstrated that the performance of RSFC-based behavioral prediction model could vary a lot with different parcellation approaches (Dadi et al., 2019; Pervaiz et al., 2020; Farahibozorg et al., 2021). Future work with more choices of parcellation and gradient approaches could potentially bring new insights into the comparison between parcellations versus gradients for RSFC-based behavioral prediction.

Similar to previous studies (Dadi et al., 2019; Pervaiz et al., 2020; Farahibozorg et al., 2021), we also found RSFC computed using full correlation (Pearson’s correlation) performed worse than partial correlation for both hard- and soft-parcellation approaches (Figures S3 and S10). This suggests that partial correlation RSFC could provide more behaviorally relevant information than full correlation RSFC. However, it is unclear what is the equivalence of partial correlation for gradient approaches. It might be possible to improve the prediction performance for gradient approaches by using a different representation. We leave this for future work.

### Conclusions

We compared 3 different parcellation approaches (Schaefer2018, Kong2021 and sICA) and 2 different gradient techniques (PrincipalGrad and LocalGrad) for RSFC prediction of behavioral measures from HCP and ABCD datasets using KRR and LRR. We showed that functional connectivity principal gradients could predict behavioral measures similar to parcellation approaches with optimized resolutions. Comparing different approaches, individual-specific hard-parcellation approach performed the best in the HCP dataset, while principal gradient and parcellation approaches performed similarly in the ABCD dataset. In both datasets, we found that the task performance measures could be predicted better than self-reported measures for all parcellation and gradient approaches. Furthermore, hard-parcellations and principal gradients with very low resolutions performed worse than high resolutions, but this is not necessarily true for soft-parcellation approach sICA. Overall, our results suggested that principal gradients with relatively high resolution (> 40 or > 60 gradients) could predict behavioral measures no worse than parcellation approaches.

## Acknowledgements

Our research is supported by the Singapore National Research Foundation (NRF) Fellowship (Class of 2017), the NUS Yong Loo Lin School of Medicine (NUHSRO/2020/124/TMR/LOA), the Singapore National Medical Research Council (NMRC) LCG (OFLCG19May-0035), NMRC STaR (STaR20nov-0003), Singapore Ministry of Health (MOH) Centre Grant (CG21APR1009) and the United States National Institutes of Health (R01MH120080). Our computational work was partially performed on resources of the National Supercomputing Centre, Singapore (https://www.nscc.sg). Any opinions, findings and conclusions or recommendations expressed in this material are those of the authors and do not reflect the views of the Singapore NRF, NMRC or MOH.

The Wellcome Centre for Integrative Neuroimaging is supported by core funding from the Wellcome Trust (203139/Z/16/Z). For the purpose of open access, the author has applied a CC BY public copyright license to any Author Accepted Manuscript version arising from this submission.

Data were provided [in part] by the Human Connectome Project, WU-Minn Consortium (Principal Investigators: David Van Essen and Kamil Ugurbil; 1U54MH091657) funded by the 16 NIH Institutes and Centers that support the NIH Blueprint for Neuroscience Research; and by the McDonnell Center for Systems Neuroscience at Washington University.

Data used in the preparation of this article were obtained from the Adolescent Brain Cognitive DevelopmentSM (ABCD) Study (https://abcdstudy.org), held in the NIMH Data Archive (NDA). This is a multisite, longitudinal study designed to recruit more than 10,000 children age 9-10 and follow them over 10 years into early adulthood. The ABCD Study® is supported by the National Institutes of Health and additional federal partners under award numbers U01DA041048, U01DA050989, U01DA051016, U01DA041022, U01DA051018, U01DA051037, U01DA050987, U01DA041174, U01DA041106, U01DA041117, U01DA041028, U01DA041134, U01DA050988, U01DA051039, U01DA041156, U01DA041025, U01DA041120, U01DA051038, U01DA041148, U01DA041093, U01DA041089, U24DA041123, U24DA041147. A full list of supporters is available at https://abcdstudy.org/federal-partners.html. A listing of participating sites and a complete listing of the study investigators can be found at https://abcdstudy.org/consortium_members/. ABCD consortium investigators designed and implemented the study and/or provided data but did not necessarily participate in the analysis or writing of this report. This manuscript reflects the views of the authors and may not reflect the opinions or views of the NIH or ABCD consortium investigators. The ABCD data repository grows and changes over time. The ABCD data used in this report came from http://dx.doi.org/10.15154/1504041.

## Supplementary Figures

**Table S1.** Site clusters for ABCD.

| **ABCD Site** | **Make** | **Model** | **N** | **Site-cluster** |
| --- | --- | --- | --- | --- |
| 16 | Siemens | Prisma | 187 | A |
| 13 | GE | Discovery MR750 | 164 | B |
| 4 | GE | Discovery MR750 | 150 | C |
| 22 | GE | Discovery MR750 | 11 | C |
| 14 | Siemens | Prisma/Prisma fit | 121 | D |
| 15 | Siemens | Prisma fit | 23 | D |
| 10 | GE | Discovery MR750 | 120 | E |
| 11 | Siemens | Prisma | 44 | E |
| 3 | Siemens | Prisma | 111 | F |
| 5 | Siemens | Prisma fit | 61 | F |
| 2 | Siemens | Prisma fit | 68 | G |
| 7 | Siemens | Prisma fit | 56 | G |
| 6 | Siemens | Prisma fit | 47 | H |
| 8 | GE | Discovery MR750 | 55 | H |
| 9 | Siemens | Prisma fit | 20 | H |
| 20 | Siemens | Prisma/Prisma fit | 61 | H |
| 12 | Siemens | Prisma fit | 50 | I |
| 18 | GE | Discovery MR750 | 72 | I |
| 21 | Siemens | Prisma fit/Prisma | 55 | I |

**Figure S1.**
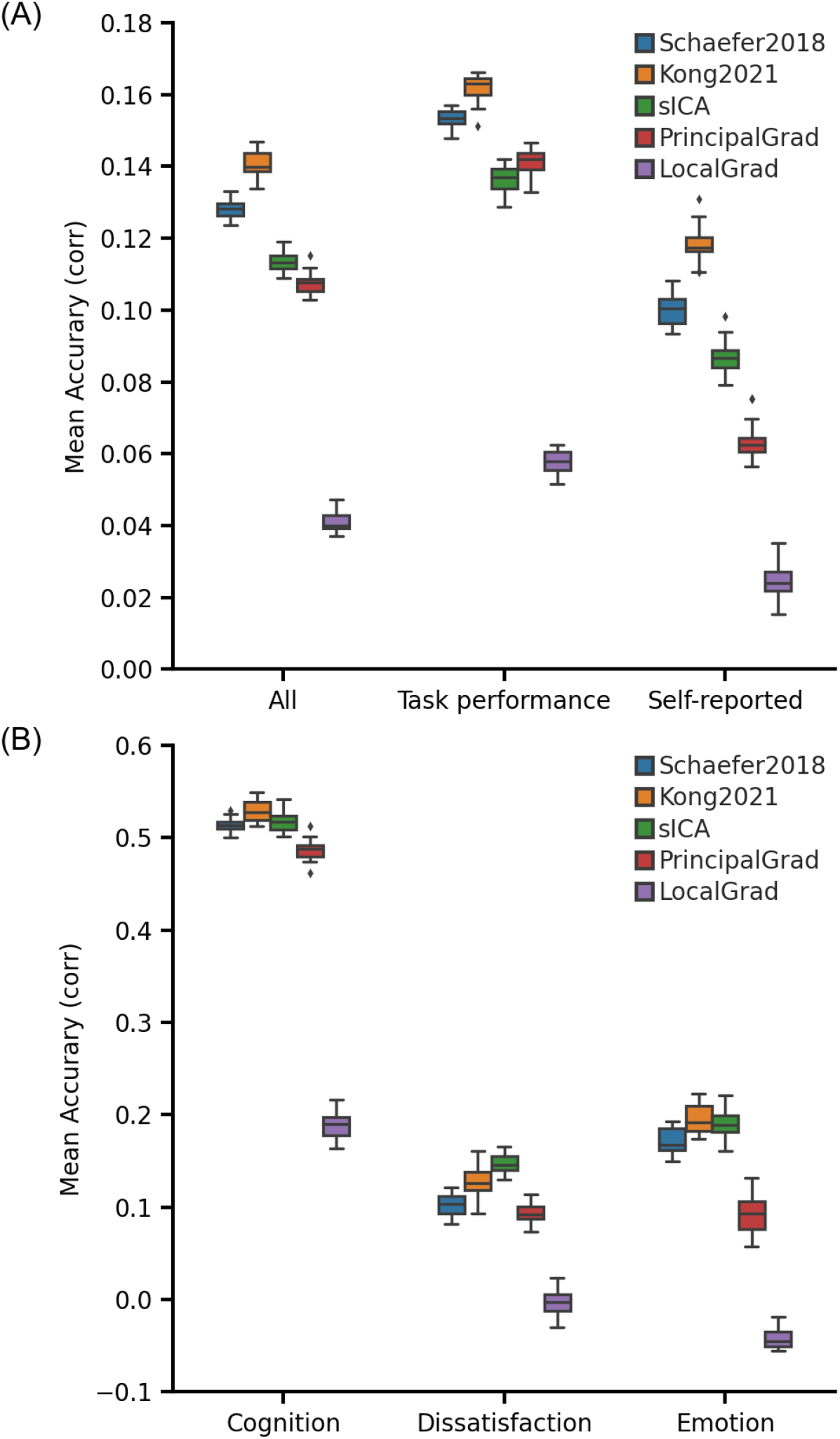
Individual-specific hard-parcellation approach Kong2021 compared favorably with other approaches for linear ridge regression (LRR) in the HCP dataset. (A) Average prediction accuracies (Pearson’s correlation) of all 58 behavioral measures, task performance measures, and self-reported measures. (B) Prediction accuracies (Pearson’s correlation) of three behavioral components: cognition, dissatisfaction, and emotion. Boxplots utilized default Python seaborn parameters, that is, box shows median and interquartile range (IQR). Whiskers indicate 1.5 IQR.

**Figure S2.**
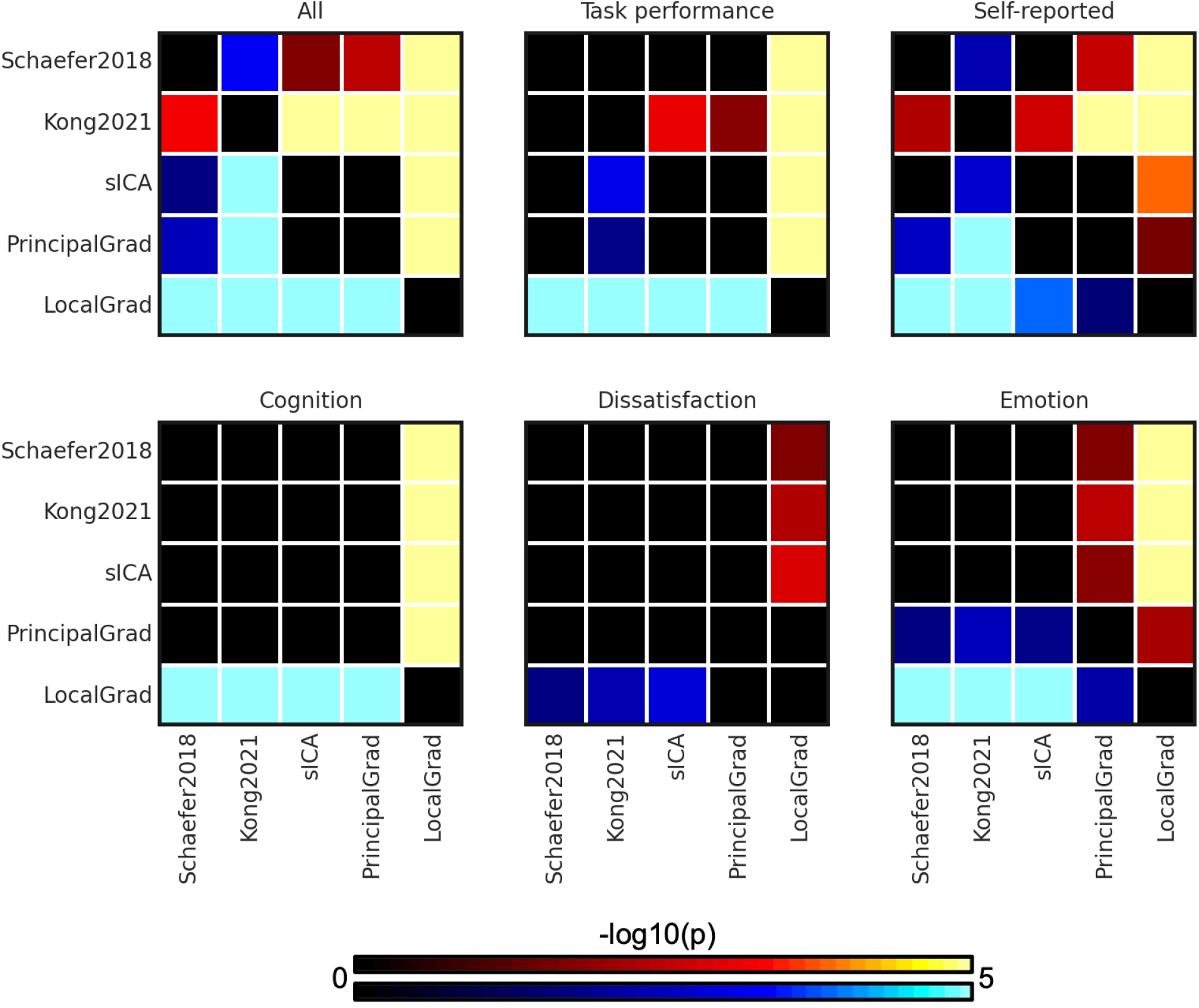
P values (-log10(p)) of comparing prediction accuracies between each pair of approaches for linear ridge regression (LRR) in the HCP dataset. Non-black colors denote significantly different prediction performances after correcting for multiple comparisons with FDR q < 0.05. Bright colors indicate small p values, dark colors indicate large p values. For each pair of comparisons, warm colors represent higher prediction accuracies of the “row” approach than the “column” approach. Individual-specific hard-parcellation approach Kong2021 compared favorably with the other approaches, as can be seen from warm colors along the rows corresponding to Kong2021.

**Figure S3.**
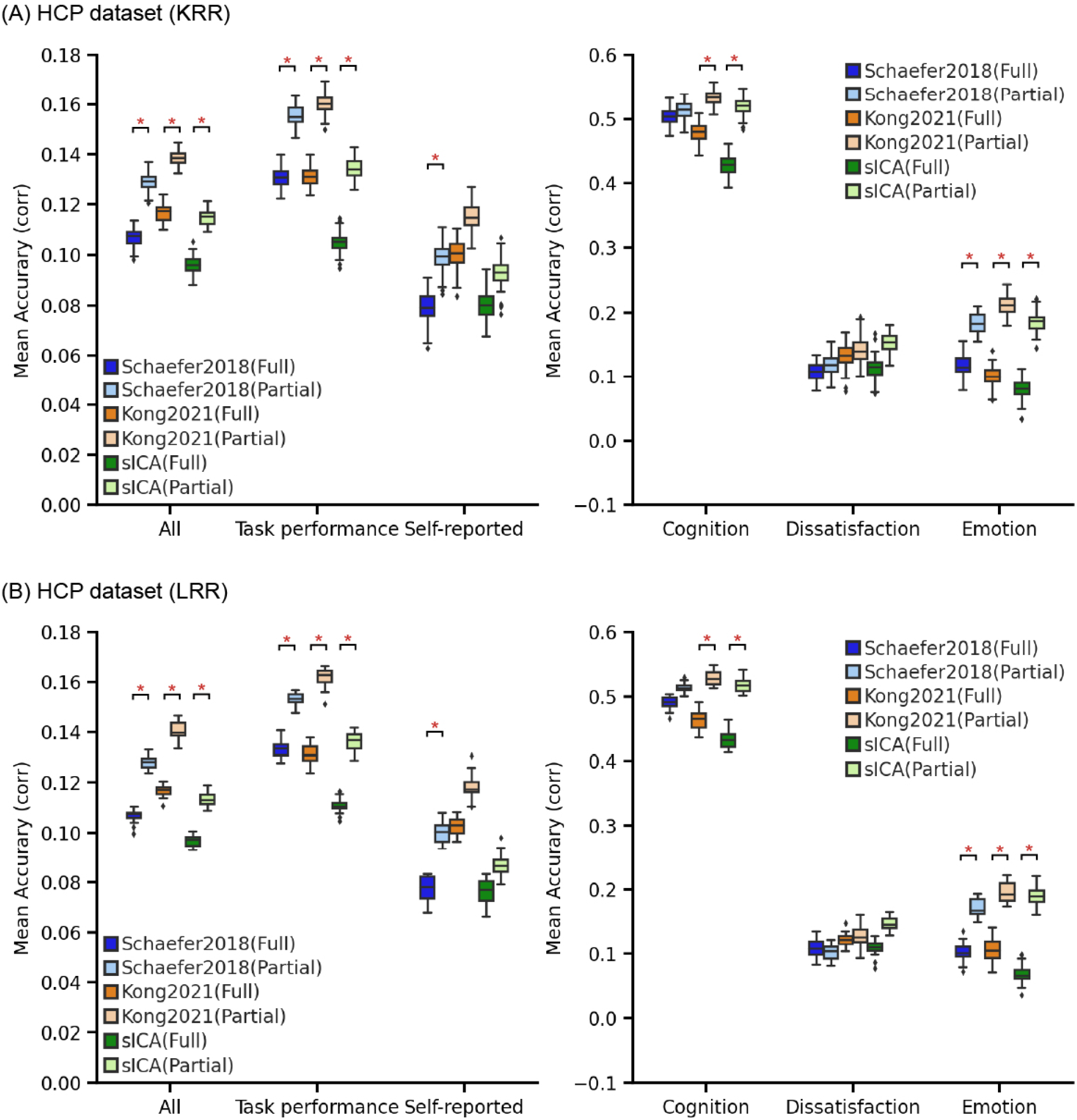
Schaefer2018, Kong201, and sICA using full correlation RSFC performed worse than using partial correlation RSFC in the HCP dataset for (A) kernel ridge regression (KRR) and (B) linear ridge regression (LRR). Boxplots utilized default Python seaborn parameters, that is, box shows median and interquartile range (IQR). Whiskers indicate 1.5 IQR. Red * represents significant p-value after FDR (q < 0.05) correction.

**Figure S4.**
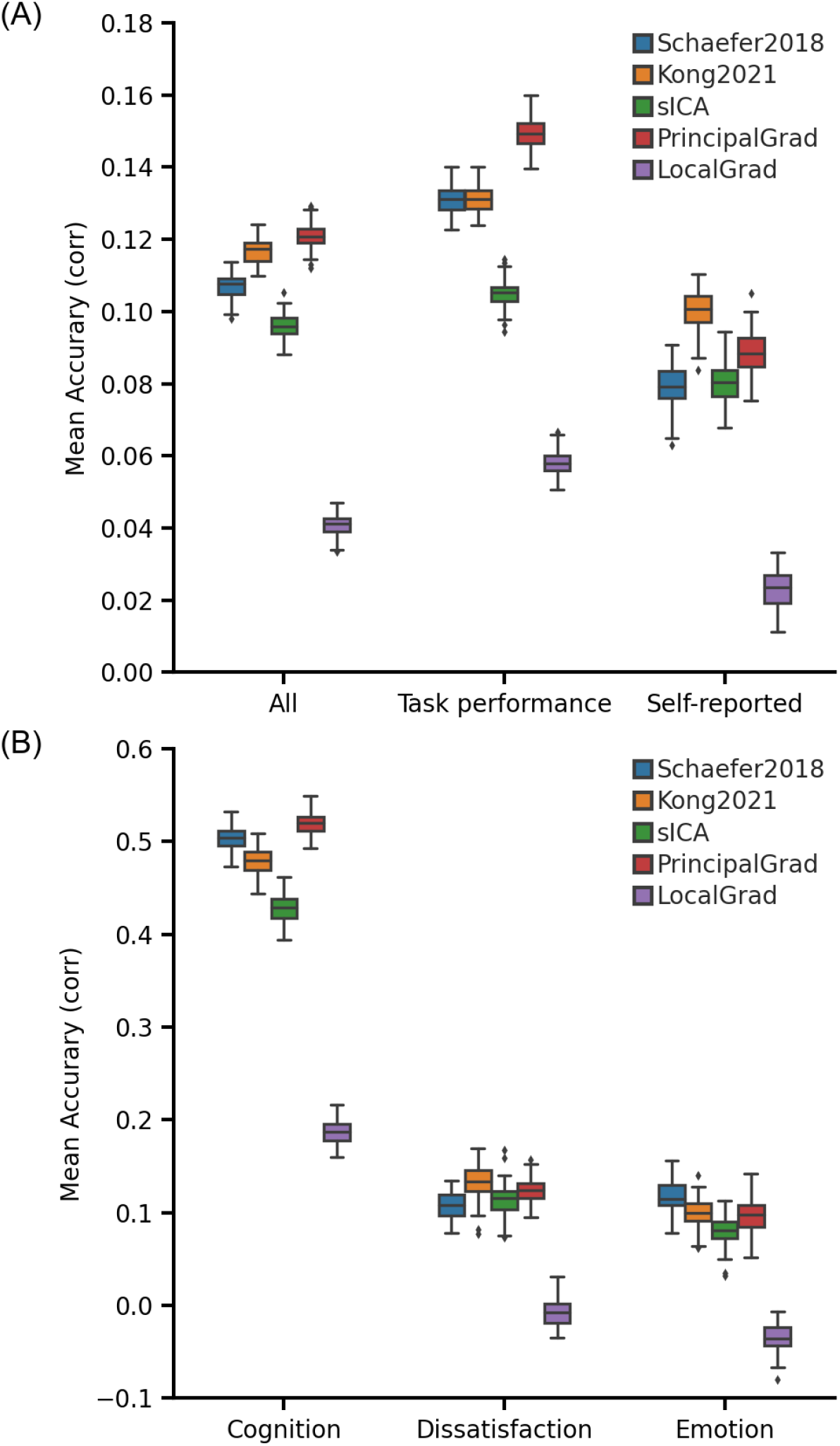
Prediction performance across different approaches using full correlation RSFC for Schaefer2018, Kong2021, and sICA with kernel ridge regression (KRR) in the HCP dataset. (A) Average prediction accuracies (Pearson’s correlation) of all 58 behavioral measures, task performance measures, and self-reported measures. (B) Prediction accuracies (Pearson’s correlation) of three behavioral components: cognition, dissatisfaction, and emotion. Boxplots utilized default Python seaborn parameters, that is, box shows median and interquartile range (IQR). Whiskers indicate 1.5 IQR.

**Figure S5.**
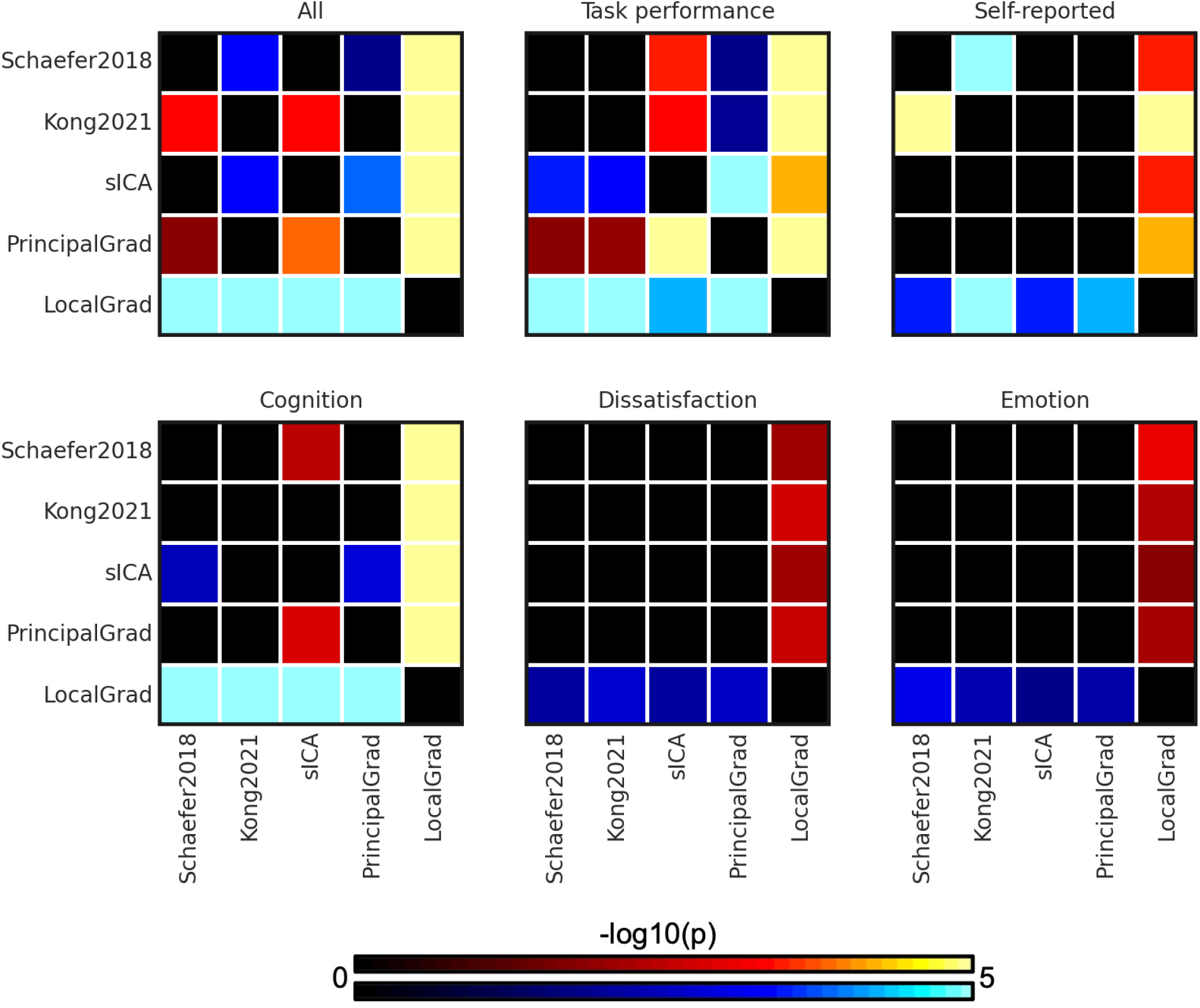
P values (-log10(p)) of comparing prediction accuracies between each pair of approaches for kernel ridge regression (KRR) in the HCP dataset. RSFC of Schaefer2018, Kong2021, and sICA were generated by full correlation. Non-black colors denote significantly different prediction performances after correcting for multiple comparisons with FDR q < 0.05. Bright colors indicate small p values, dark colors indicate large p values. For each pair of comparisons, warm colors represent higher prediction accuracies of the “row” approach than the “column” approach.

**Figure S6.**
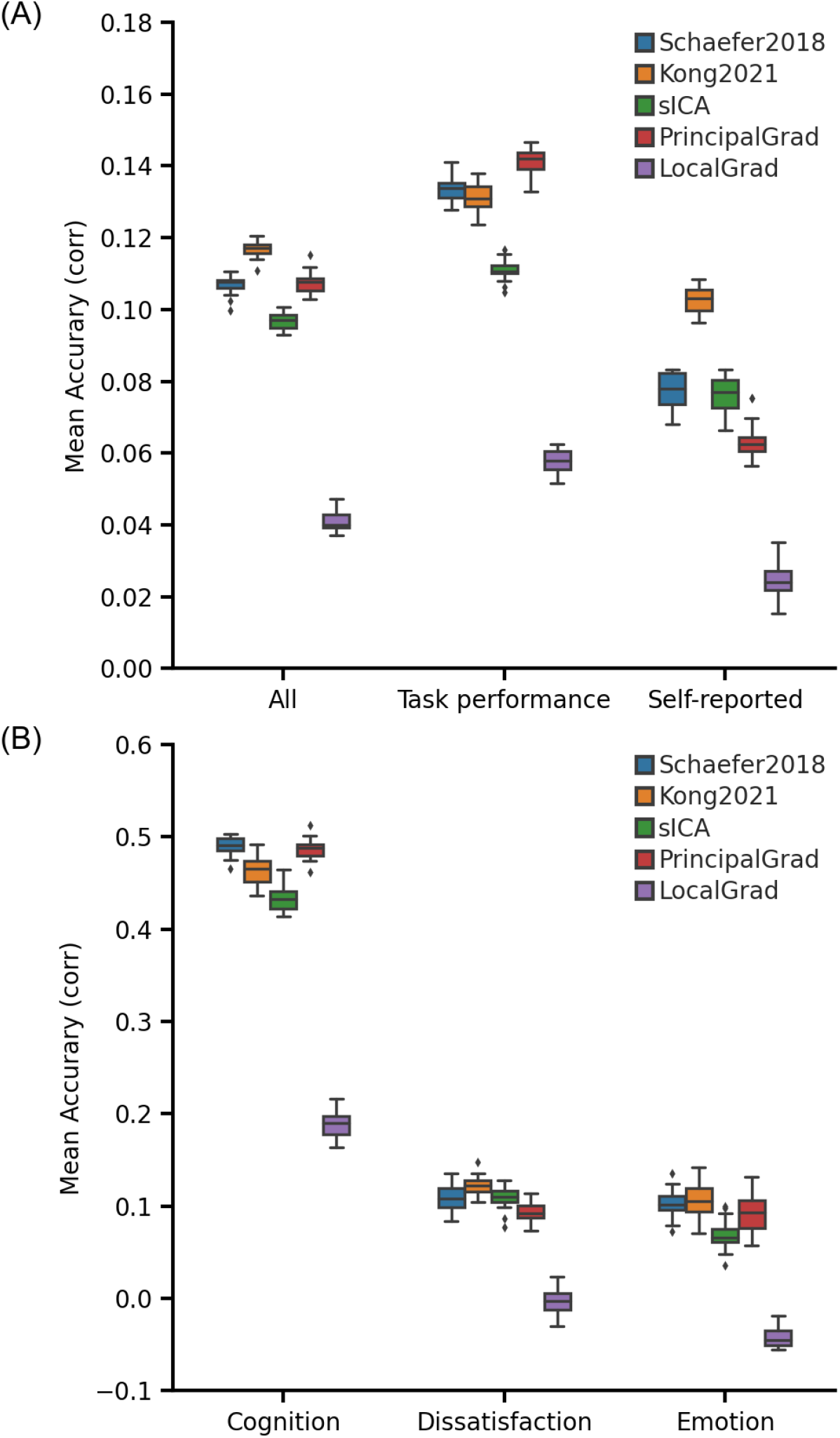
Prediction performance across different approaches using full correlation RSFC for Schaefer2018, Kong2021, and sICA with linear ridge regression (LRR) in the HCP dataset. (A) Average prediction accuracies (Pearson’s correlation) of all 58 behavioral measures, task performance measures, and self-reported measures. (B) Prediction accuracies (Pearson’s correlation) of three behavioral components: cognition, dissatisfaction, and emotion. Boxplots utilized default Python seaborn parameters, that is, box shows median and interquartile range (IQR). Whiskers indicate 1.5 IQR.

**Figure S7.**
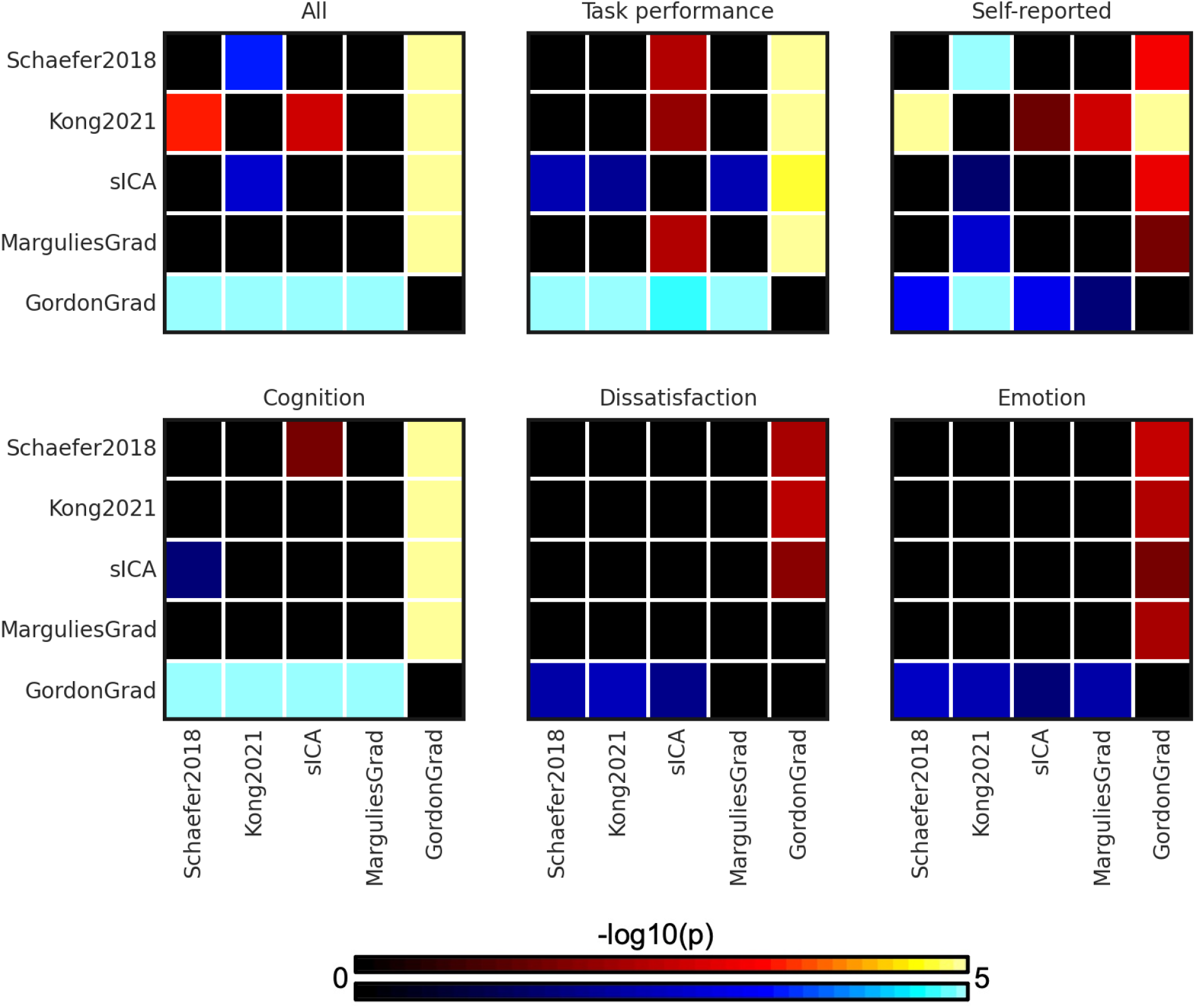
P values (-log10(p)) of comparing prediction accuracies between each pair of approaches for linear ridge regression (LRR) in the HCP dataset. RSFC of Schaefer2018, Kong2021, and sICA were generated by full correlation. Non-black colors denote significantly different prediction performances after correcting for multiple comparisons with FDR q < 0.05. Bright colors indicate small p values, dark colors indicate large p values. For each pair of comparisons, warm colors represent higher prediction accuracies of the “row” approach than the “column” approach.

**Figure S8.**
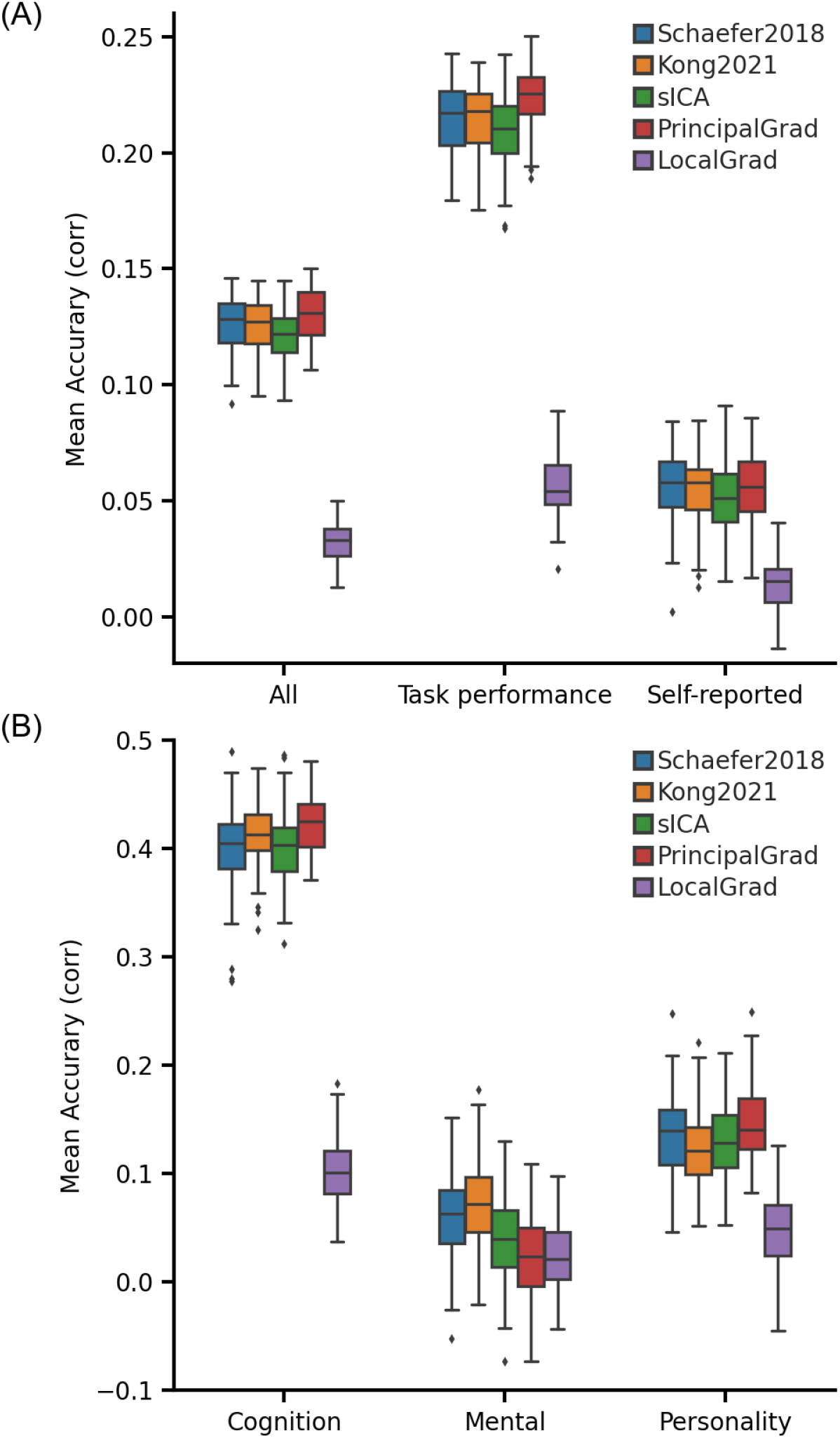
Principal gradient approach achieves comparable behavioral prediction performance as parcellation approaches for linear ridge regression (LRR) in the ABCD dataset. (A) Average prediction accuracies (Pearson’s correlation) of all 36 behavioral measures, task performance measures, and self-reported measures. (B) Prediction accuracies (Pearson’s correlation) of three behavioral components: cognition, mental health, and personality. Boxplots utilized default Python seaborn parameters, that is, box shows median and interquartile range (IQR). Whiskers indicate 1.5 IQR. The principal gradient approach PrincipalGrad was numerically the best for most cases, but there was largely no statistical difference among the approaches.

**Figure S9.**
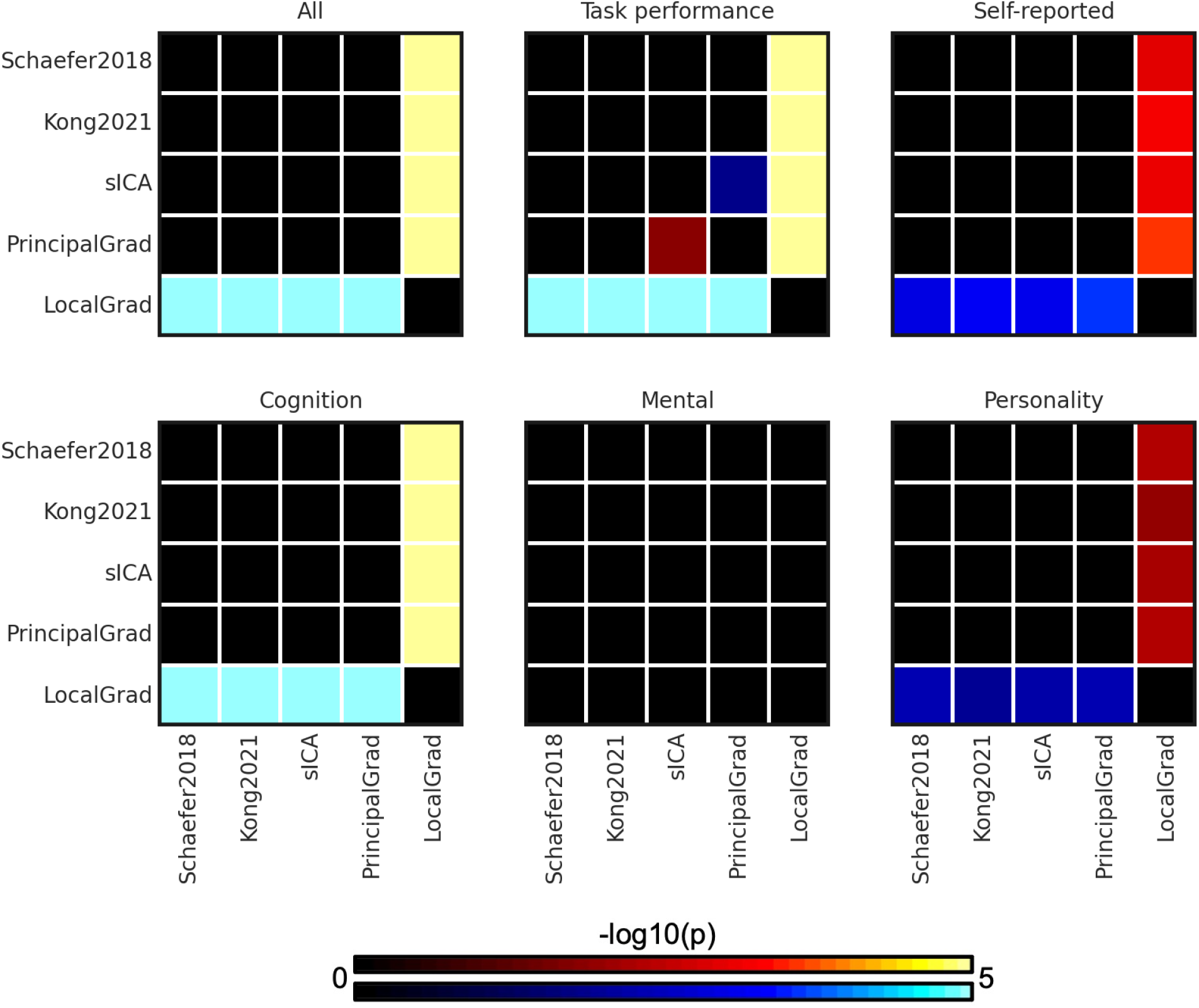
P values (-log10(p)) of comparing prediction accuracies between each pair of approaches for linear ridge regression (LRR) in the ABCD dataset. Non-black colors denote significantly different prediction performances after correcting for multiple comparisons with FDR q < 0.05. Bright colors indicate small p values, dark colors indicate large p values. For each pair of comparisons, warm colors represent higher prediction accuracies of the “row” approach than the “column” approach. There was no statistical difference among most approaches.

**Figure S10.**
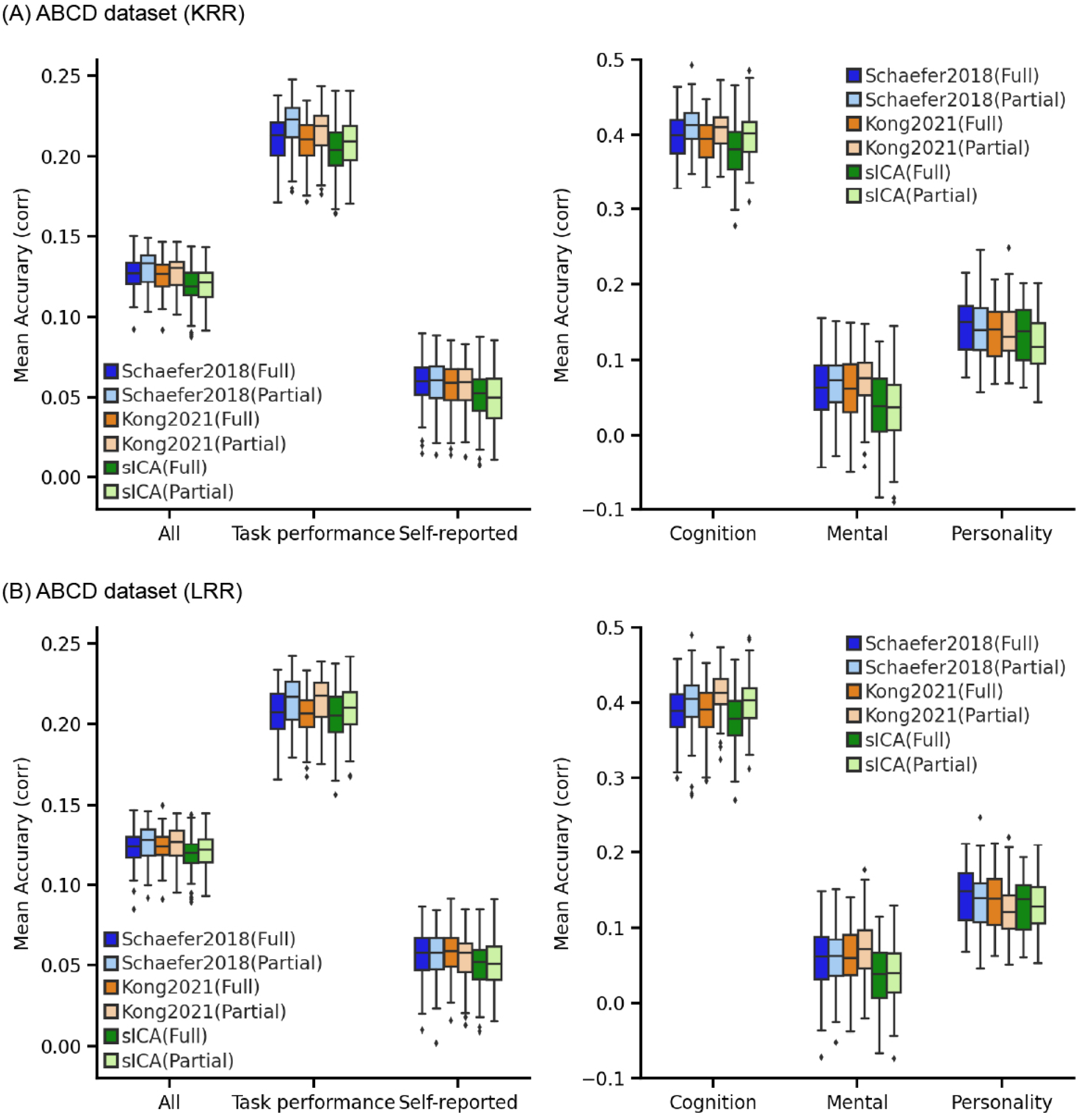
Schaefer2018, Kong201, and sICA using full correlation RSFC performed numerically worse than using partial correlation RSFC in the ABCD dataset for (A) kernel ridge regression (KRR) and (B) linear ridge regression (LRR), but differences were not significant. Boxplots utilized default Python seaborn parameters, that is, box shows median and interquartile range (IQR). Whiskers indicate 1.5 IQR.

**Figure S11.**
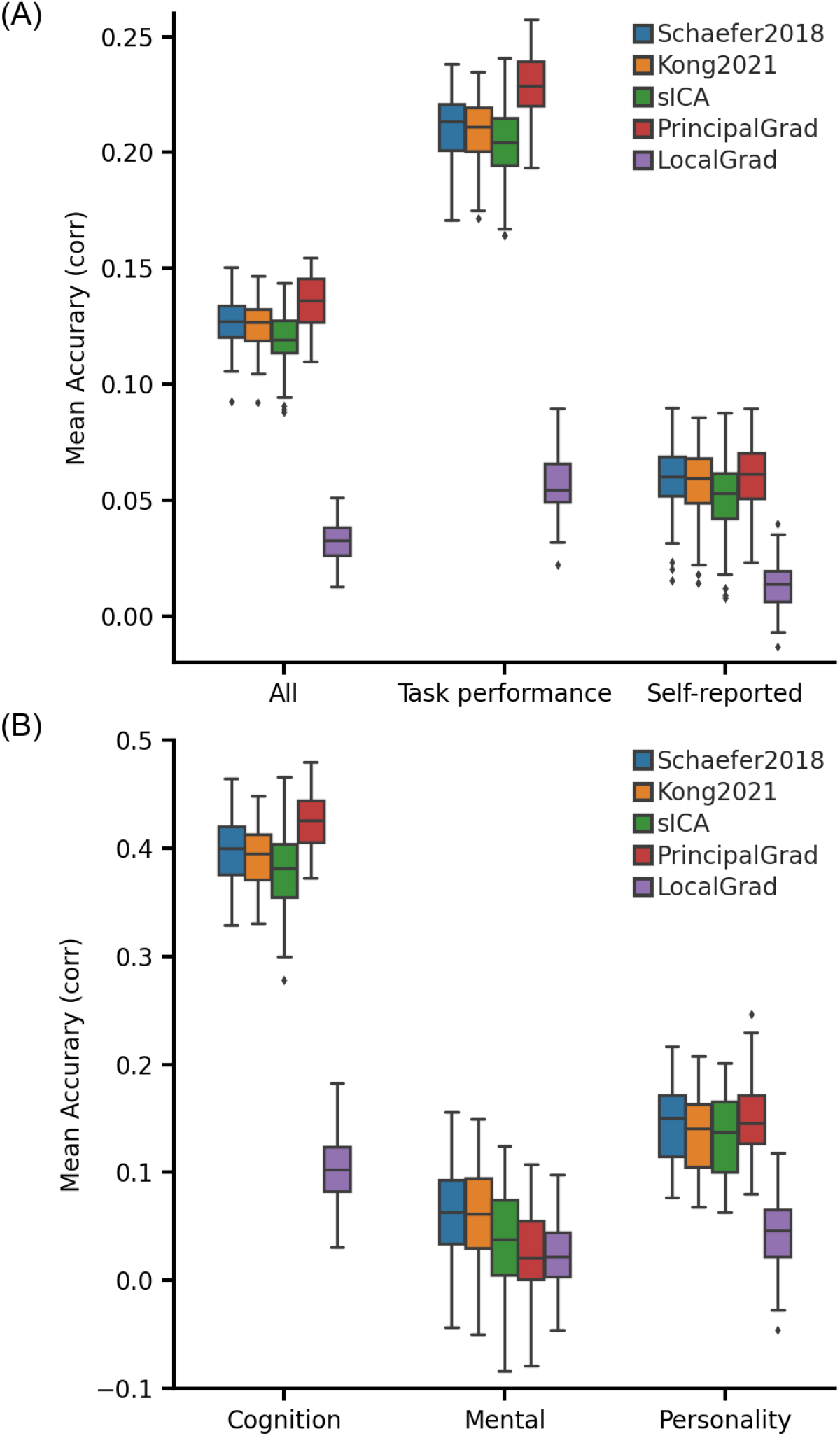
Prediction performance across different approaches using full correlation RSFC for Schaefer2018, Kong2021, and sICA with kernel ridge regression (KRR) in the ABCD dataset. (A) Average prediction accuracies (Pearson’s correlation) of all 36 behavioral measures, task performance measures, and self-reported measures. (B) Prediction accuracies (Pearson’s correlation) of three behavioral components: cognition, mental health, and personality. Boxplots utilized default Python seaborn parameters, that is, box shows median and interquartile range (IQR). Whiskers indicate 1.5 IQR. The principal gradient approach PrincipalGrad was numerically the best for most cases, but there was largely no statistical difference among the approaches.

**Figure S12.**
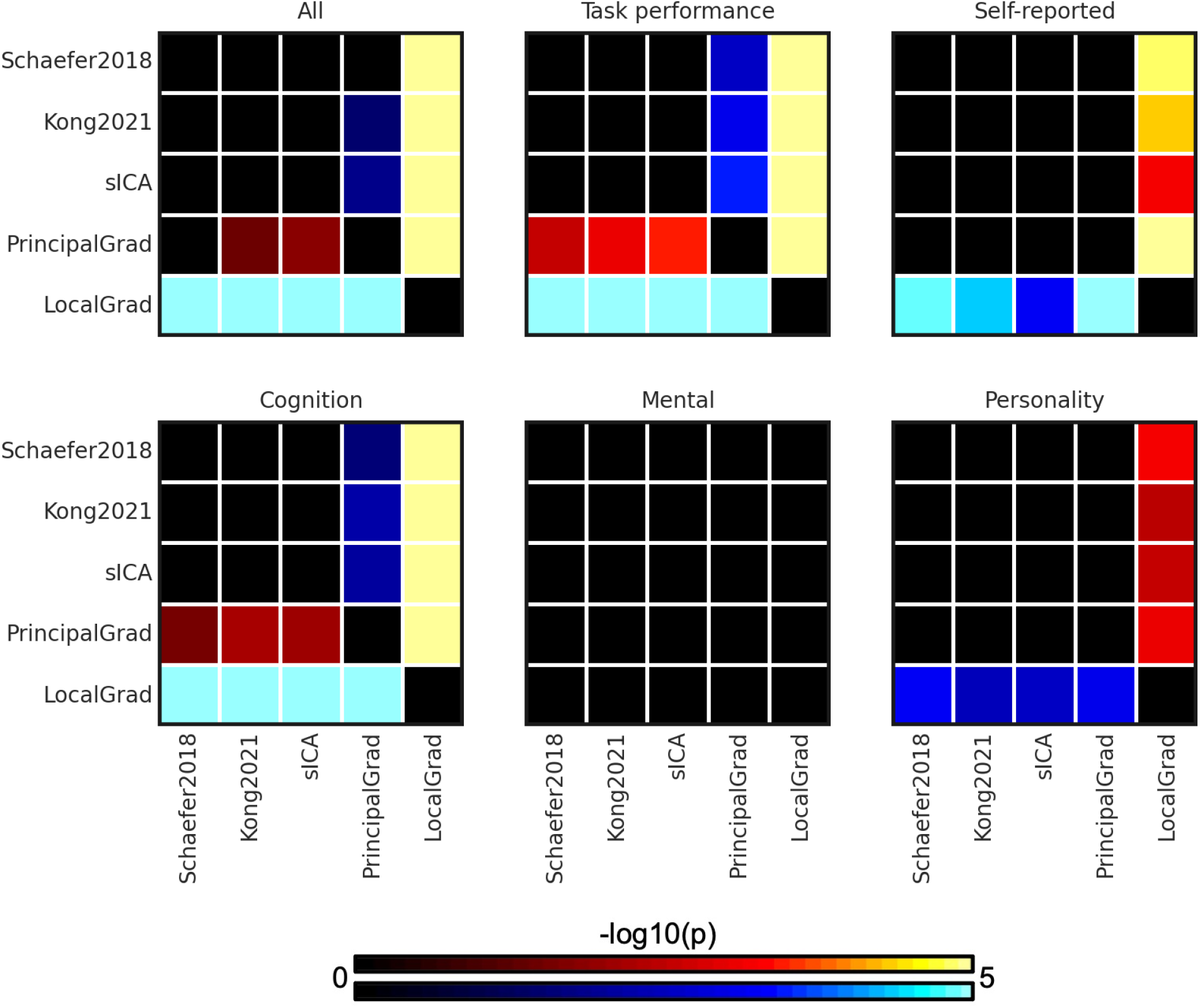
P values (-log10(p)) of comparing prediction accuracies between each pair of approaches for kernel ridge regression (KRR) in the ABCD dataset. RSFC of Schaefer2018, Kong2021, and sICA were generated by full correlation. Non-black colors denote significantly different prediction performances after correcting for multiple comparisons with FDR q < 0.05. Bright colors indicate small p values, dark colors indicate large p values. For each pair of comparisons, warm colors represent higher prediction accuracies of the “row” approach than the “column” approach.

**Figure S13.**
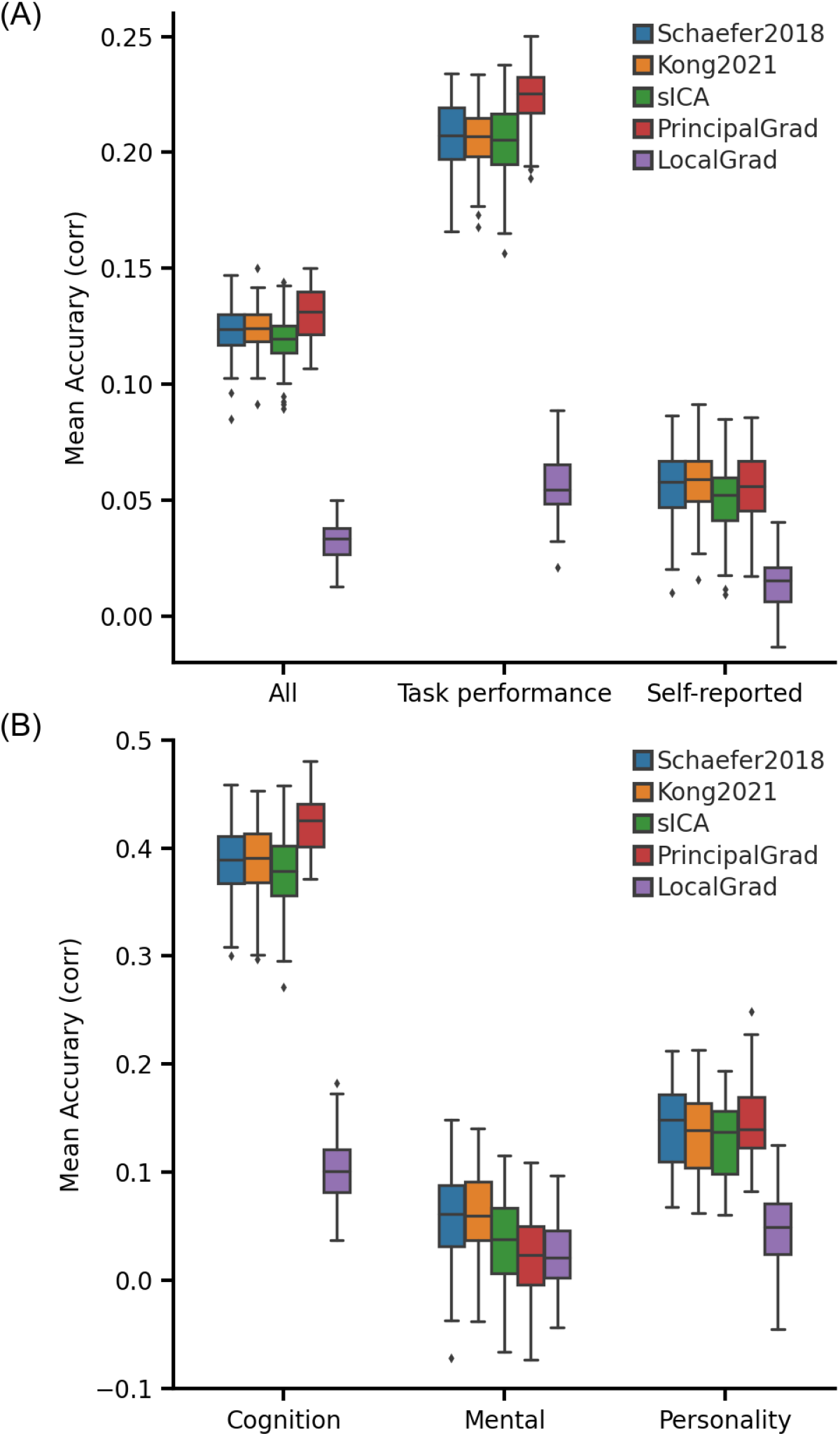
Prediction performance across different approaches using full correlation RSFC for Schaefer2018, Kong2021, and sICA with linear ridge regression (LRR) in the ABCD dataset. (A) Average prediction accuracies (Pearson’s correlation) of all 36 behavioral measures, task performance measures, and self-reported measures. (B) Prediction accuracies (Pearson’s correlation) of three behavioral components: cognition, mental health, and personality. Boxplots utilized default Python seaborn parameters, that is, box shows median and interquartile range (IQR). Whiskers indicate 1.5 IQR.

**Figure S14.**
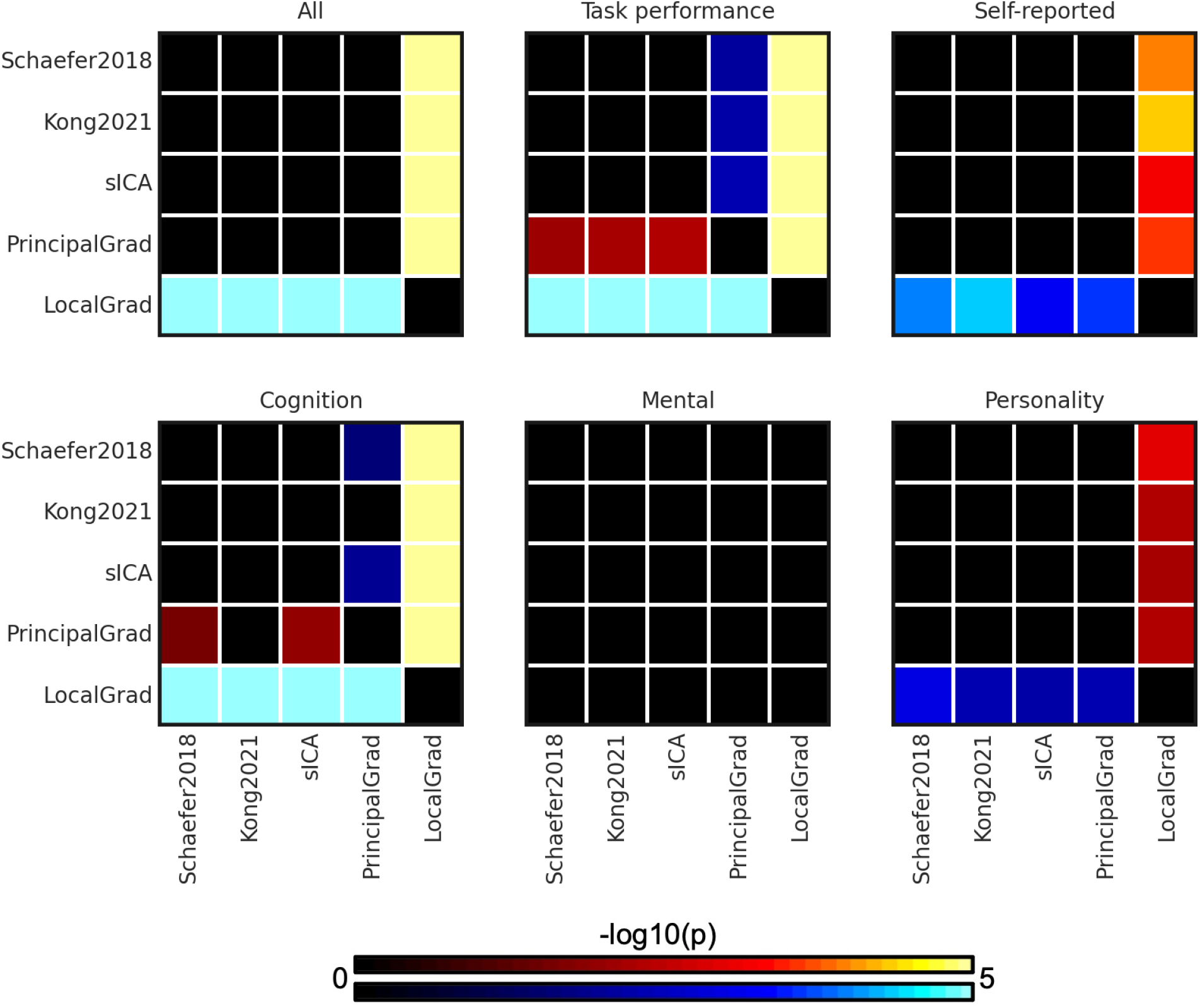
P values (-log10(p)) of comparing prediction accuracies between each pair of approaches for linear ridge regression (LRR) in the ABCD dataset. RSFC of Schaefer2018, Kong2021, and sICA were generated by full correlation. Non-black colors denote significantly different prediction performances after correcting for multiple comparisons with FDR q < 0.05. Bright colors indicate small p values, dark colors indicate large p values. For each pair of comparisons, warm colors represent higher prediction accuracies of the “row” approach than the “column” approach.

**Figure S15.**
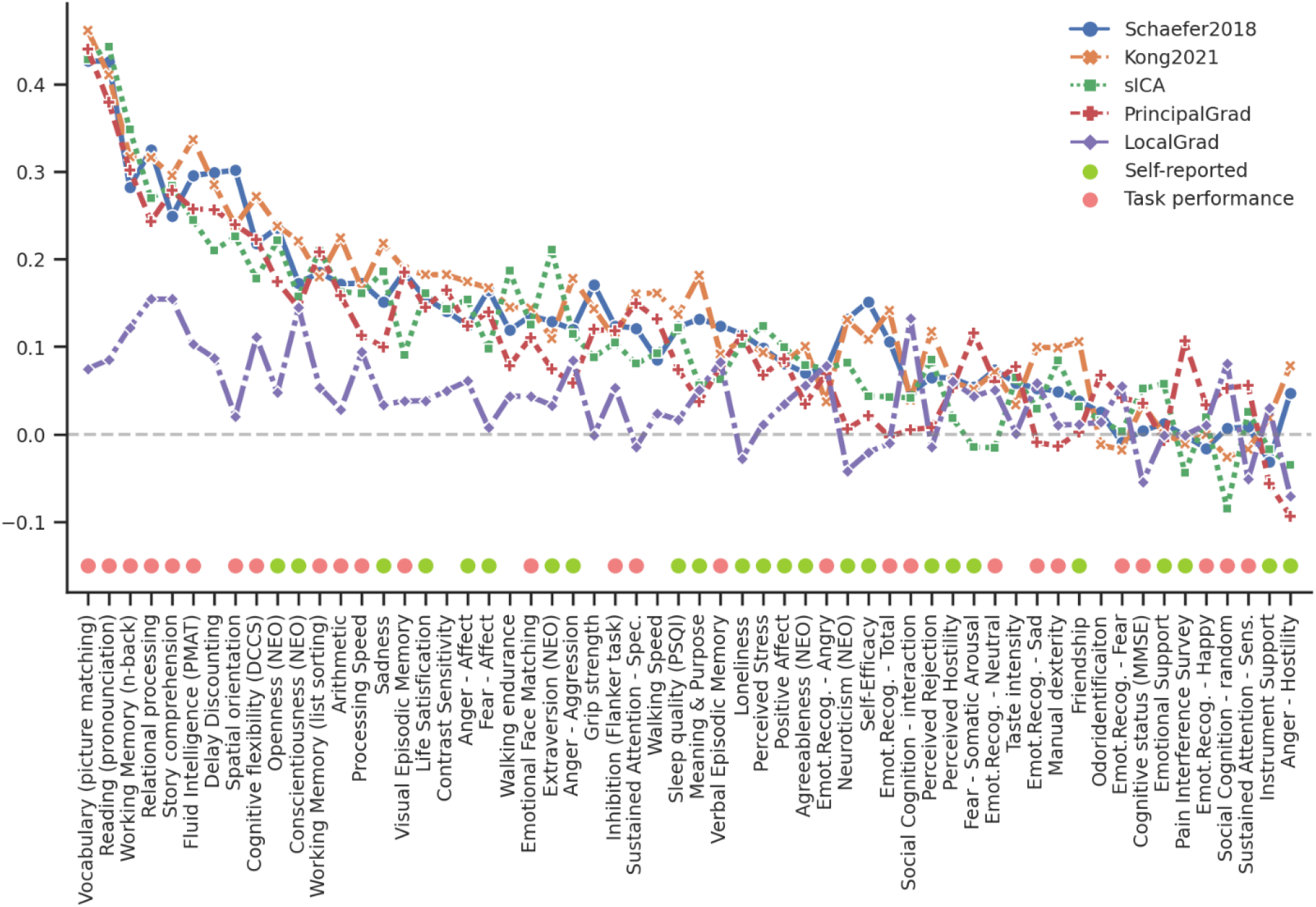
Task performance measures were predicted better than self-reported measures across different gradient and parcellation approaches with optimized resolutions for LRR in the HCP dataset. 58 behavioral measures were ordered based on average prediction accuracies across Schaefer2018, Kong2021, sICA, PrincipalGrad, and LocalGrad. Pink circles indicate task performance measures. Green circles indicate self-reported measures. Boxplots utilized default Python seaborn parameters, that is, box shows median and interquartile range (IQR). Whiskers indicate 1.5 IQR. Designation of behavioral measures into “self-reported” and “task-performance” measures followed previous studies (Li et al., 2019a; Liégeois et al., 2019; Kong et al., 2021a).

**Figure S16.**
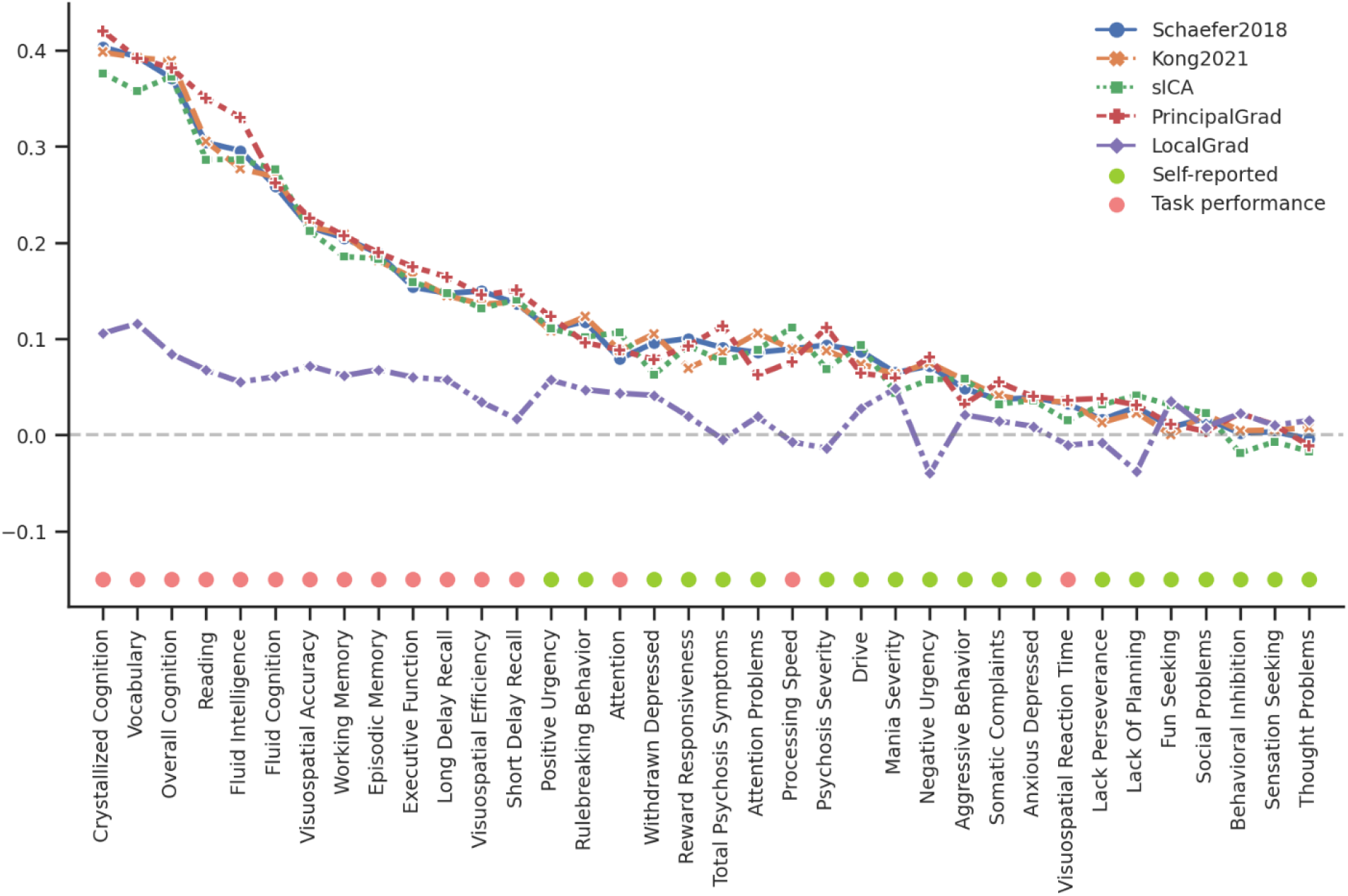
Task performance measures were predicted better than self-reported measures across different gradient and parcellation approaches with optimized resolutions for LRR in the ABCD dataset. 36 behavioral measures were ordered based on average prediction accuracies across Schaefer2018, Kong2021, sICA, PrincipalGrad, and LocalGrad. Pink circles indicate task performance measures. Green circles indicate self-reported measures. Boxplots utilized default Python seaborn parameters, that is, box shows median and interquartile range (IQR). Whiskers indicate 1.5 IQR. Designation of behavioral measures into “self-reported” and “task-performance” measures based on ABCD behavioral measures description (Li et al., 2019a; Liégeois et al., 2019; Kong et al., 2021a).

**Figure S17.**
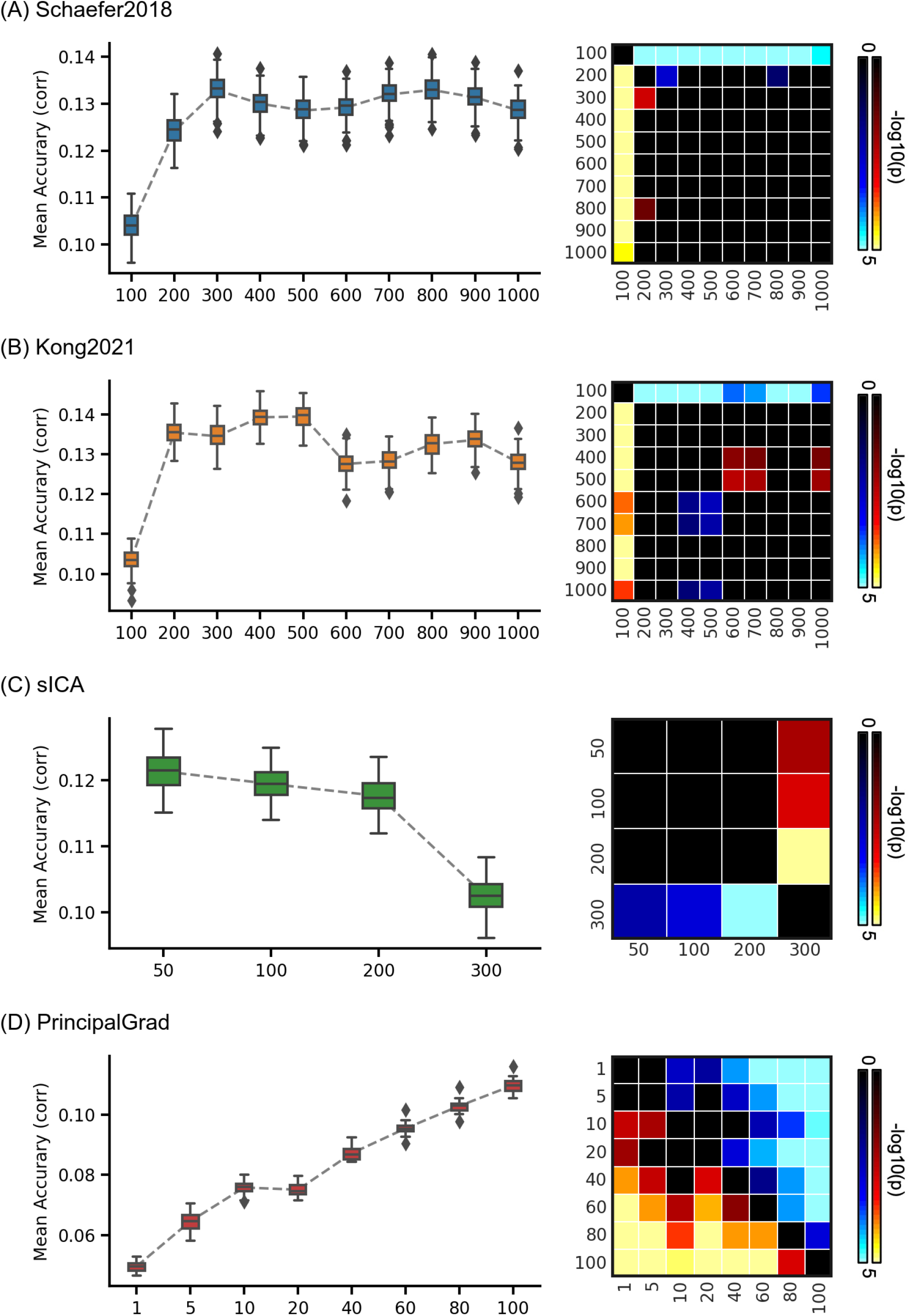
Average prediction accuracies (Pearson’s correlation) of all 58 behavioral measures vary across resolutions for gradient and parcellation approaches using LRR in the HCP dataset. (A) Prediction accuracies and p values of the hard-parcellation Schaefer2018 with 100 to 1000 ROIs. (B) Prediction accuracies and p values of the hard-parcellation Kong2021 with 100 to 1000 ROIs. (C) Prediction accuracies and p values of the soft-parcellation sICA with 50 to 300 components. (D) Prediction accuracies and p values of the principal gradient PrincipalGrad with 1 to 100 gradients. Boxplots utilized default Python seaborn parameters, that is, box shows median and interquartile range (IQR). Whiskers indicate 1.5 IQR. P values (-log10(p)) were computed between prediction accuracies of each pair of resolutions. Non-black colors denote significantly different prediction performances after correcting for multiple comparisons with FDR q < 0.05. Bright colors indicate small p values, dark colors indicate large p values. For each pair of comparisons, warm colors represent higher prediction accuracies of the “row” resolution than the “column” resolution.

**Figure S18.**
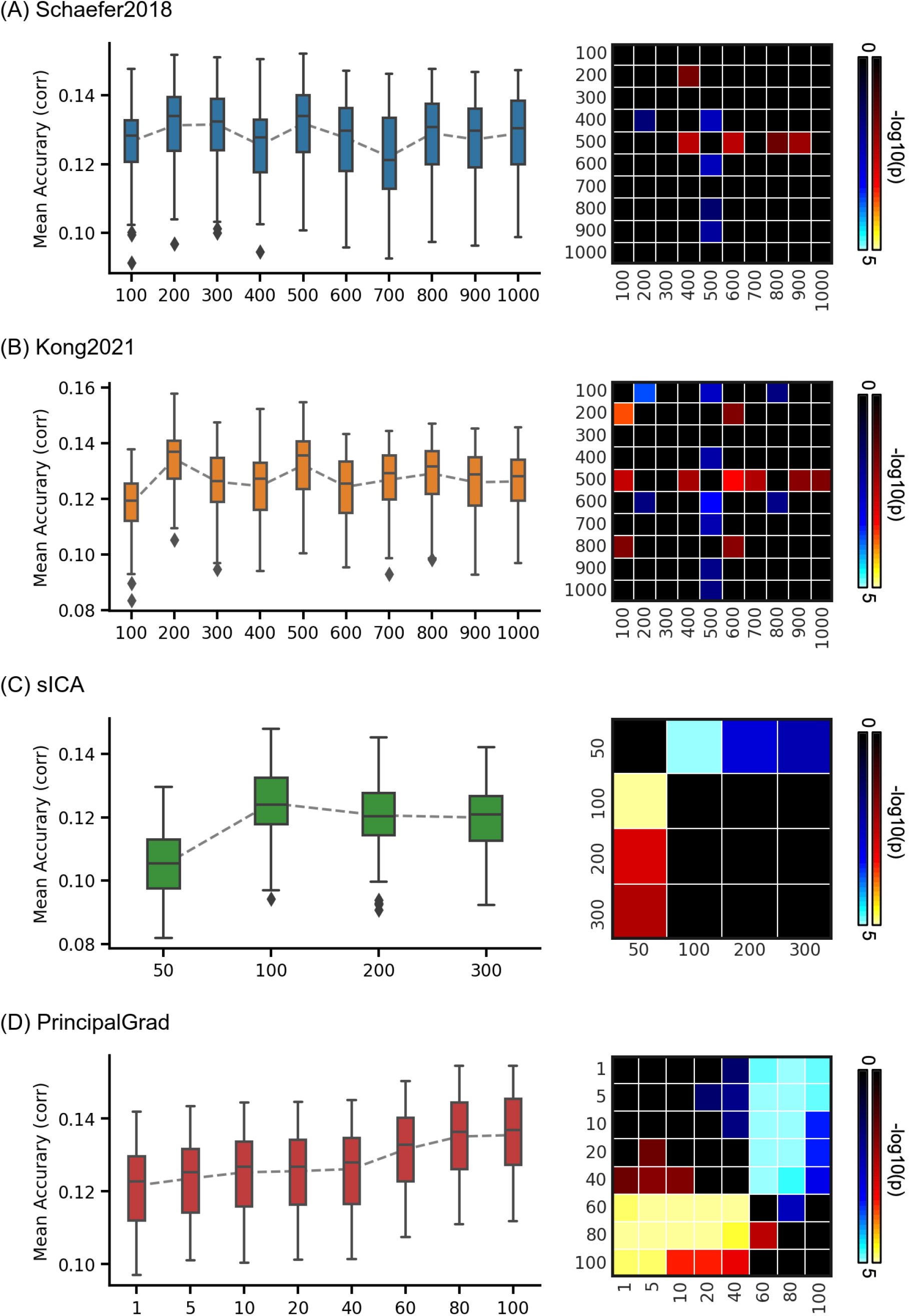
Average prediction accuracies (Pearson’s correlation) of all 36 behavioral measures vary across resolutions for gradient and parcellation approaches using LRR in the ABCD dataset. (A) Prediction accuracies and p values of the hard-parcellation Schaefer2018 with 100 to 1000 ROIs. (B) Prediction accuracies and p values of the hard-parcellation Kong2021 with 100 to 1000 ROIs. (C) Prediction accuracies and p values of the soft-parcellation sICA with 50 to 300 components. (D) Prediction accuracies and p values of the principal gradient PrincipalGrad with 1 to 100 gradients. Boxplots utilized default Python seaborn parameters, that is, box shows median and interquartile range (IQR). Whiskers indicate 1.5 IQR. P values (-log10(p)) were computed between prediction accuracies of each pair of resolutions. Non-black colors denote significantly different prediction performances after correcting for multiple comparisons with FDR q < 0.05. Bright colors indicate small p values, dark colors indicate large p values. For each pair of comparisons, warm colors represent higher prediction accuracies of the “row” resolution than the “column” resolution.

**Figure S19.**
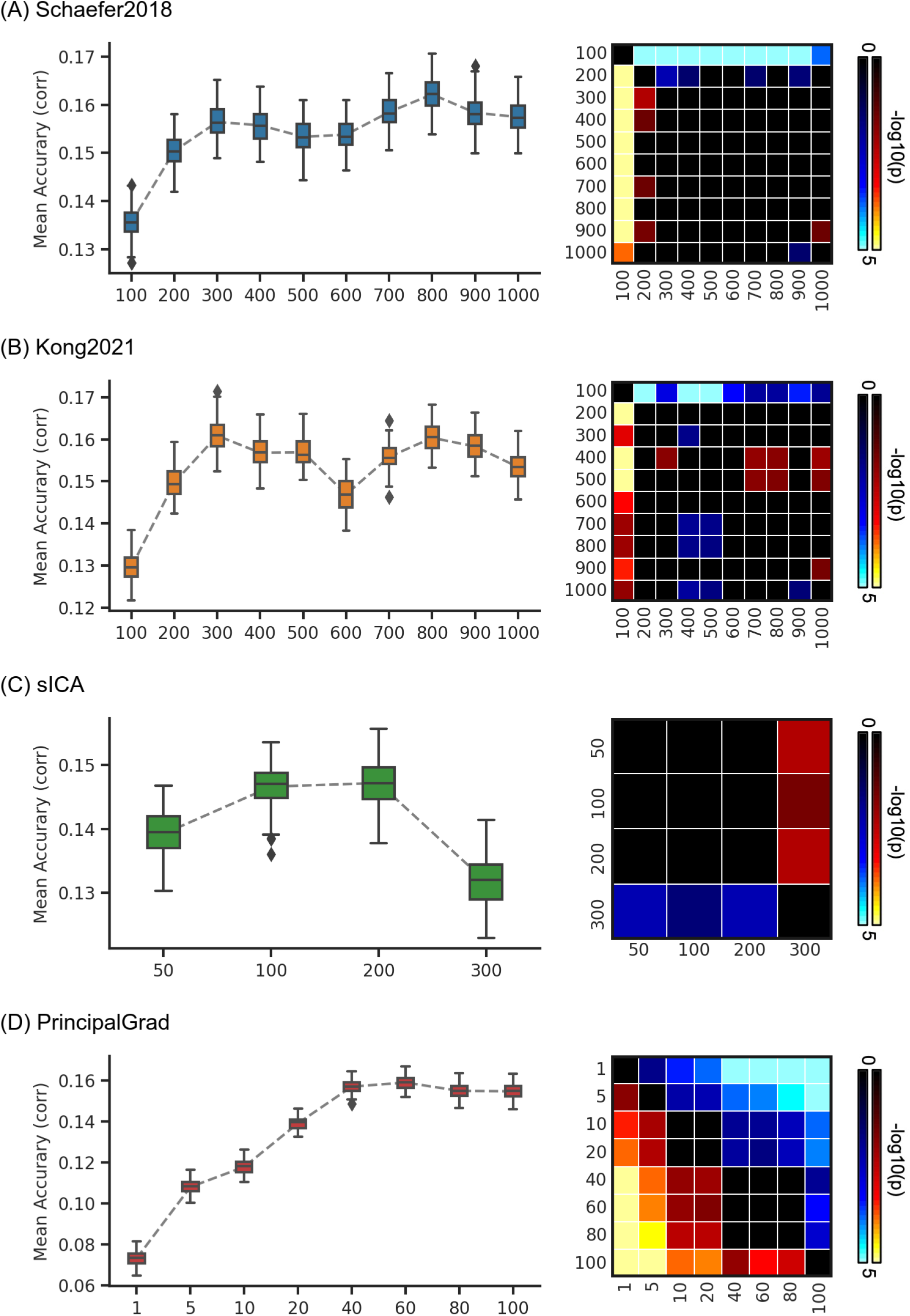
Average prediction accuracies (Pearson’s correlation) of task performance measures vary across resolutions for gradient and parcellation approaches using KRR in the HCP dataset. (A) Prediction accuracies and p values of the hard-parcellation Schaefer2018 with 100 to 1000 ROIs. (B) Prediction accuracies and p values of the hard-parcellation Kong2021 with 100 to 1000 ROIs. (C) Prediction accuracies and p values of the soft-parcellation sICA with 50 to 300 components. (D) Prediction accuracies and p values of the principal gradient PrincipalGrad with 1 to 100 gradients. Boxplots utilized default Python seaborn parameters, that is, box shows median and interquartile range (IQR). Whiskers indicate 1.5 IQR. P values (-log10(p)) were computed between prediction accuracies of each pair of resolutions. Non-black colors denote significantly different prediction performances after correcting for multiple comparisons with FDR q < 0.05. Bright colors indicate small p values, dark colors indicate large p values. For each pair of comparisons, warm colors represent higher prediction accuracies of the “row” resolution than the “column” resolution.

**Figure S20.**
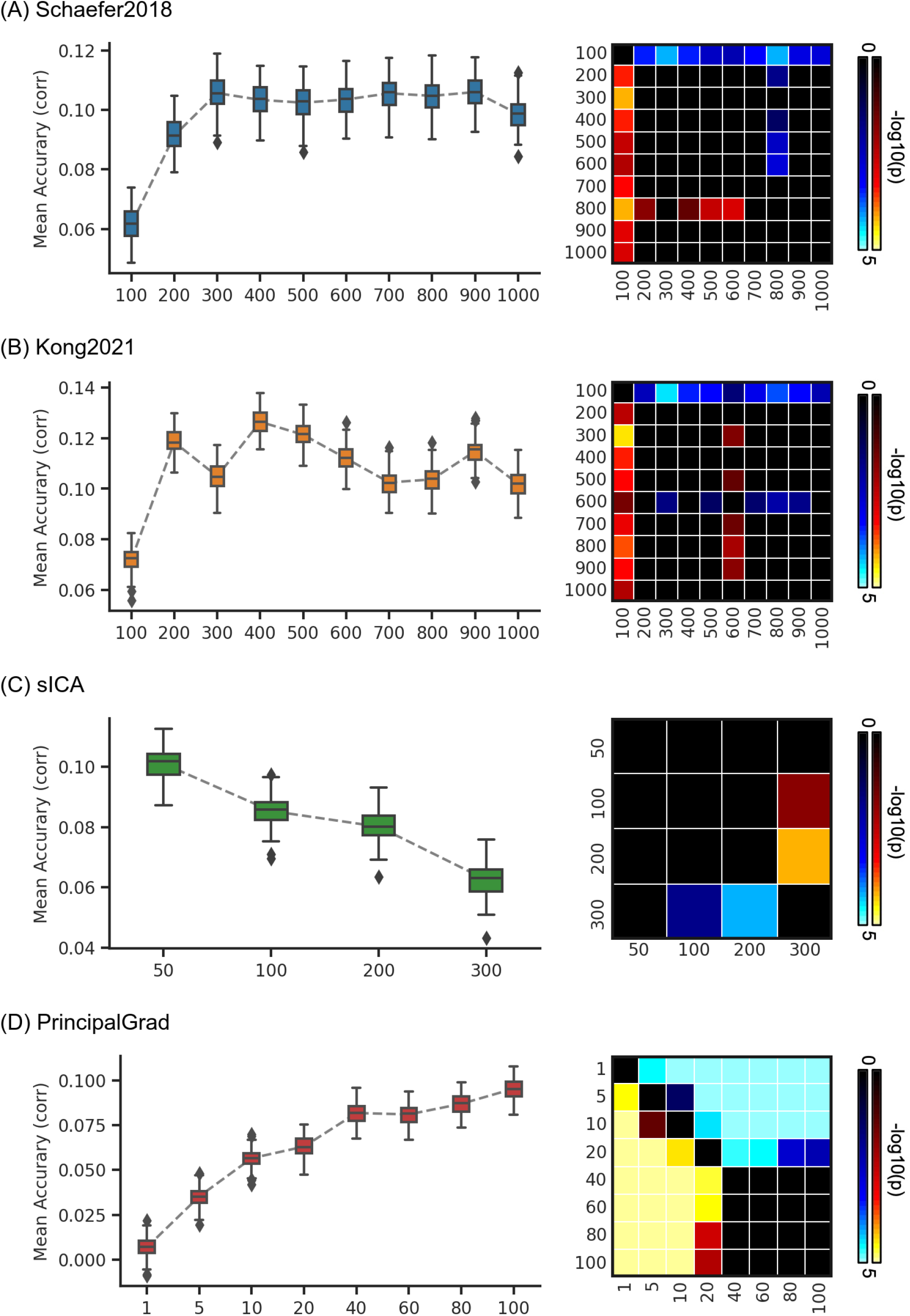
Average prediction accuracies (Pearson’s correlation) of self-reported measures vary across resolutions for gradient and parcellation approaches using KRR in the HCP dataset. (A) Prediction accuracies and p values of the hard-parcellation Schaefer2018 with 100 to 1000 ROIs. (B) Prediction accuracies and p values of the hard-parcellation Kong2021 with 100 to 1000 ROIs. (C) Prediction accuracies and p values of the soft-parcellation sICA with 50 to 300 components. (D) Prediction accuracies and p values of the principal gradient PrincipalGrad with 1 to 100 gradients. Boxplots utilized default Python seaborn parameters, that is, box shows median and interquartile range (IQR). Whiskers indicate 1.5 IQR. P values (-log10(p)) were computed between prediction accuracies of each pair of resolutions. Non-black colors denote significantly different prediction performances after correcting for multiple comparisons with FDR q < 0.05. Bright colors indicate small p values, dark colors indicate large p values. For each pair of comparisons, warm colors represent higher prediction accuracies of the “row” resolution than the “column” resolution.

**Figure S21.**
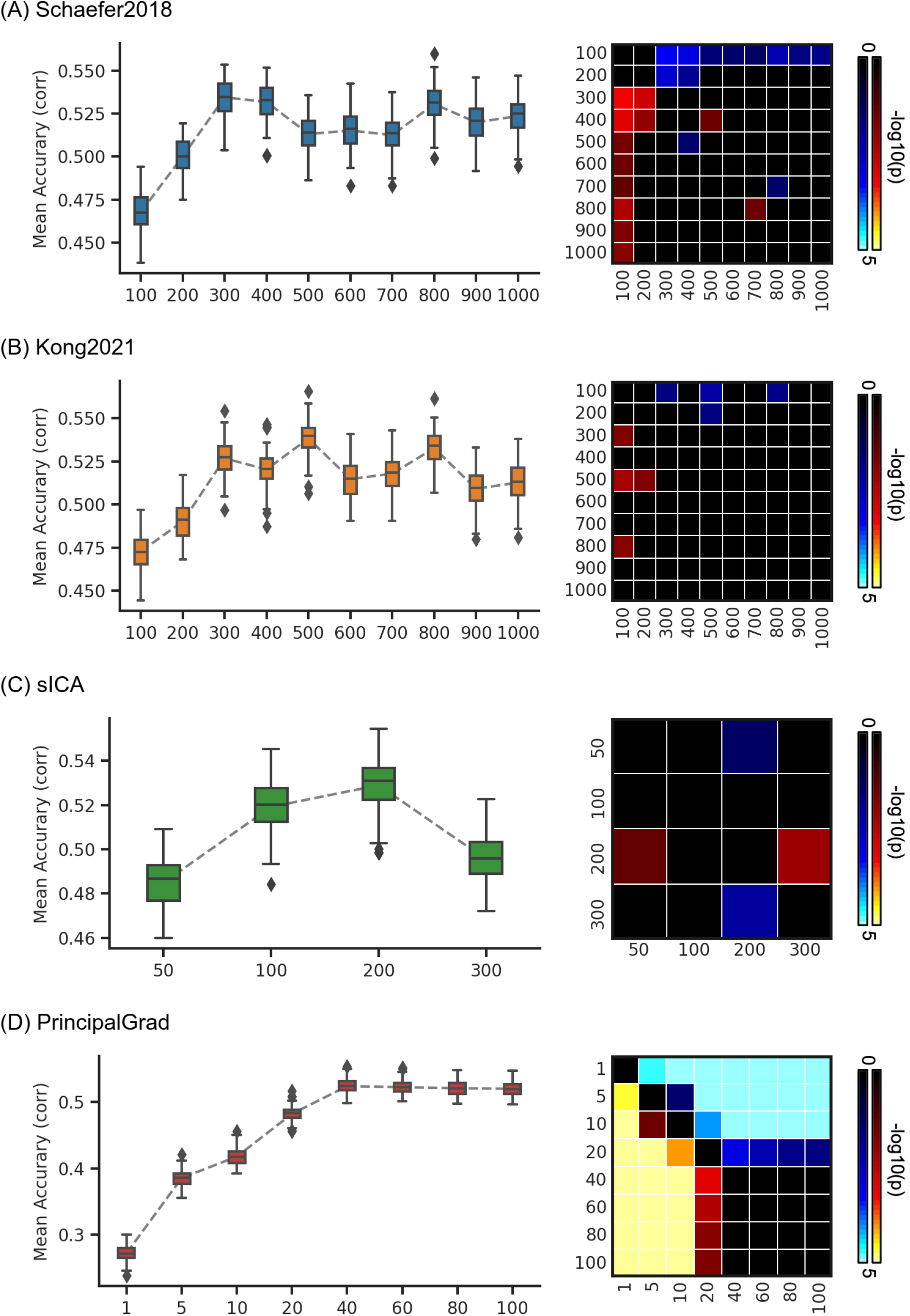
Prediction accuracies (Pearson’s correlation) of cognition vary across resolutions for gradient and parcellation approaches using KRR in the HCP dataset. (A) Prediction accuracies and p values of the hard-parcellation Schaefer2018 with 100 to 1000 ROIs. (B) Prediction accuracies and p values of the hard-parcellation Kong2021 with 100 to 1000 ROIs. (C) Prediction accuracies and p values of the soft-parcellation sICA with 50 to 300 components. (D) Prediction accuracies and p values of the principal gradient PrincipalGrad with 1 to 100 gradients. Boxplots utilized default Python seaborn parameters, that is, box shows median and interquartile range (IQR). Whiskers indicate 1.5 IQR. P values (-log10(p)) were computed between prediction accuracies of each pair of resolutions. Non-black colors denote significantly different prediction performances after correcting for multiple comparisons with FDR q < 0.05. Bright colors indicate small p values, dark colors indicate large p values. For each pair of comparisons, warm colors represent higher prediction accuracies of the “row” resolution than the “column” resolution.

**Figure S22.**
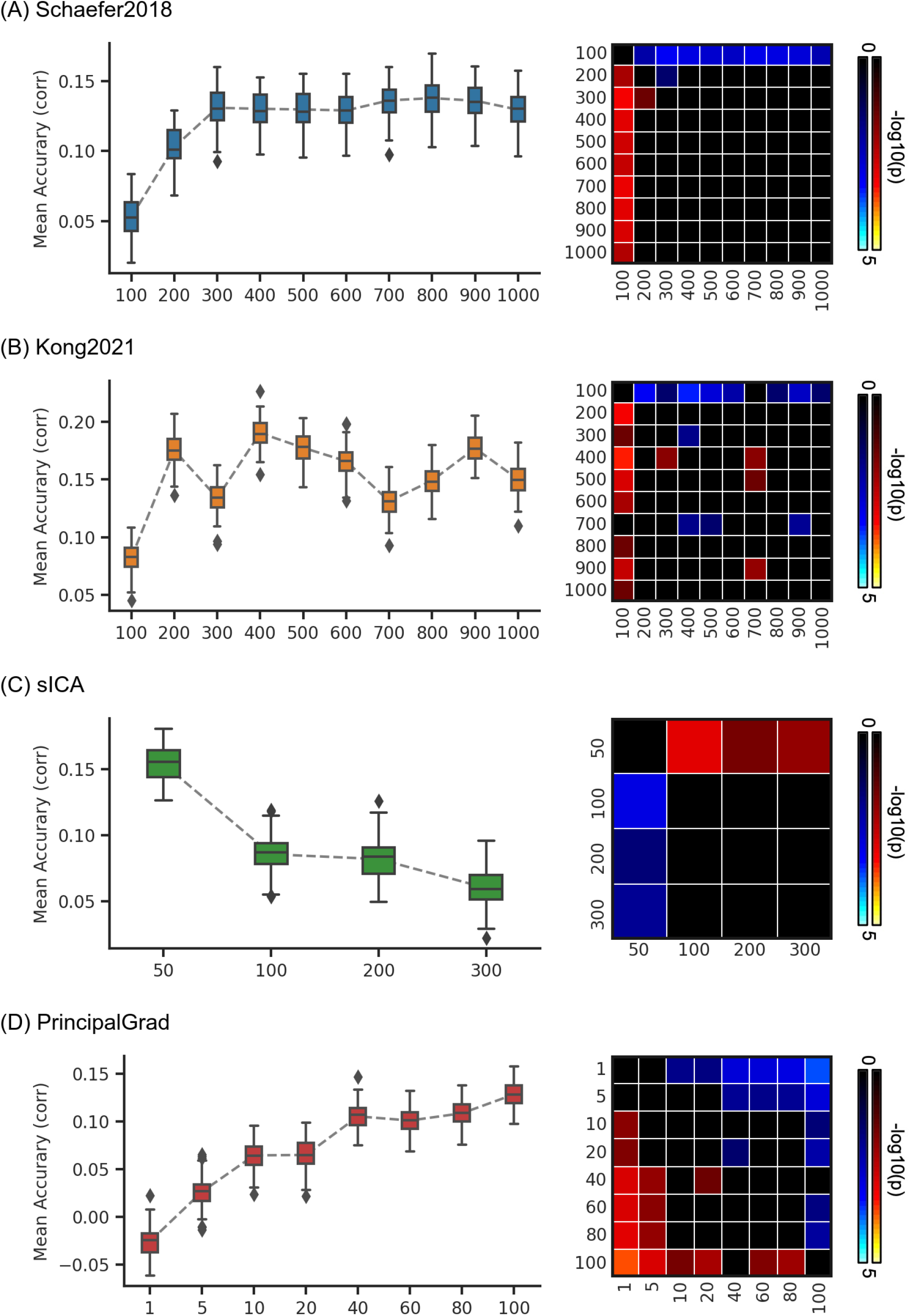
Prediction accuracies (Pearson’s correlation) of dissatisfaction vary across resolutions for gradient and parcellation approaches using KRR in the HCP dataset. (A) Prediction accuracies and p values of the hard-parcellation Schaefer2018 with 100 to 1000 ROIs. (B) Prediction accuracies and p values of the hard-parcellation Kong2021 with 100 to 1000 ROIs. (C) Prediction accuracies and p values of the soft-parcellation sICA with 50 to 300 components. (D) Prediction accuracies and p values of the principal gradient PrincipalGrad with 1 to 100 gradients. Boxplots utilized default Python seaborn parameters, that is, box shows median and interquartile range (IQR). Whiskers indicate 1.5 IQR. P values (-log10(p)) were computed between prediction accuracies of each pair of resolutions. Non-black colors denote significantly different prediction performances after correcting for multiple comparisons with FDR q < 0.05. Bright colors indicate small p values, dark colors indicate large p values. For each pair of comparisons, warm colors represent higher prediction accuracies of the “row” resolution than the “column” resolution.

**Figure S23.**
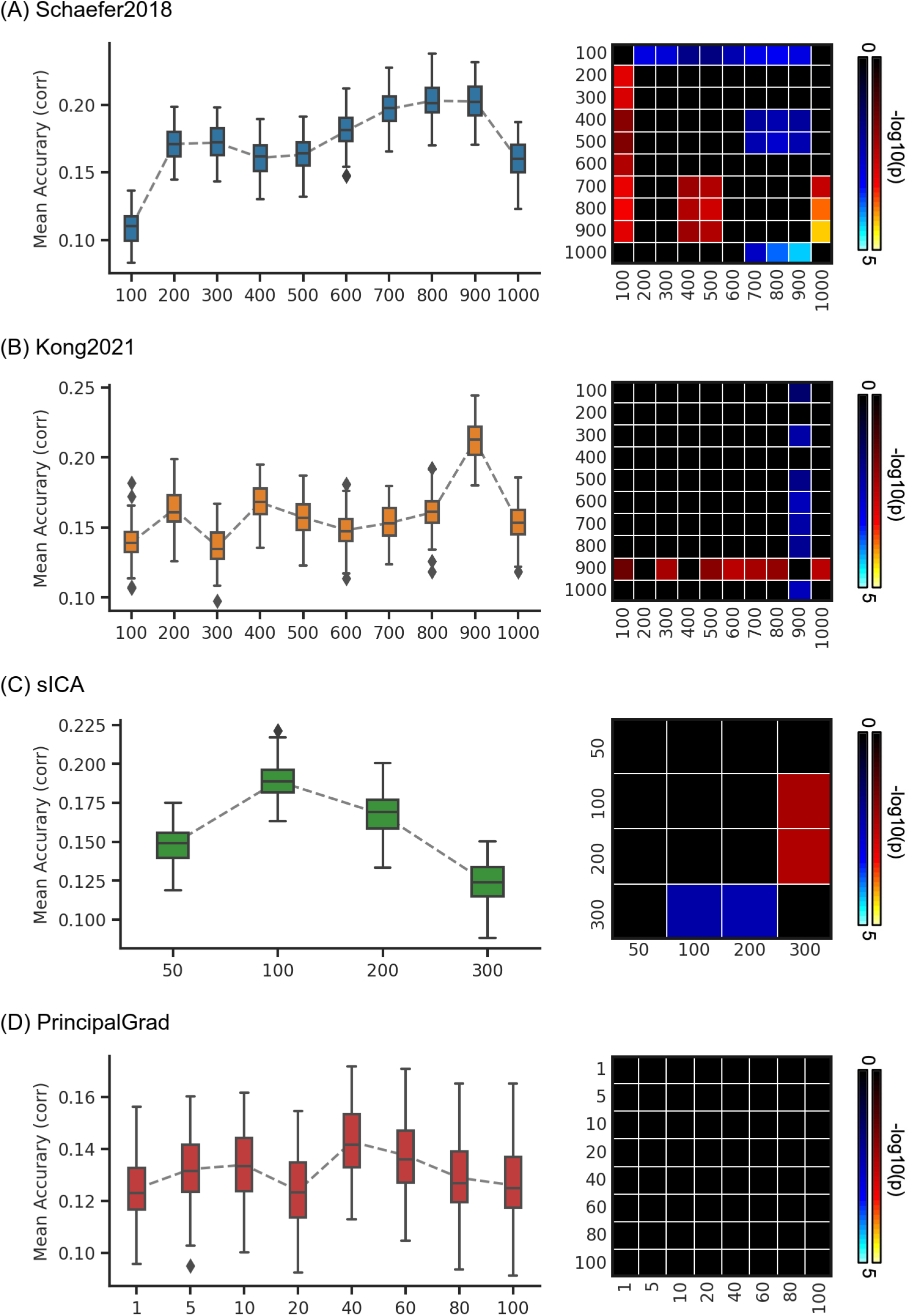
Prediction accuracies (Pearson’s correlation) of emotion vary across resolutions for gradient and parcellation approaches using KRR in the HCP dataset. (A) Prediction accuracies and p values of the hard-parcellation Schaefer2018 with 100 to 1000 ROIs. (B) Prediction accuracies and p values of the hard-parcellation Kong2021 with 100 to 1000 ROIs. (C) Prediction accuracies and p values of the soft-parcellation sICA with 50 to 300 components. (D) Prediction accuracies and p values of the principal gradient PrincipalGrad with 1 to 100 gradients. Boxplots utilized default Python seaborn parameters, that is, box shows median and interquartile range (IQR). Whiskers indicate 1.5 IQR. P values (-log10(p)) were computed between prediction accuracies of each pair of resolutions. Non-black colors denote significantly different prediction performances after correcting for multiple comparisons with FDR q < 0.05. Bright colors indicate small p values, dark colors indicate large p values. For each pair of comparisons, warm colors represent higher prediction accuracies of the “row” resolution than the “column” resolution.

**Figure S24.**
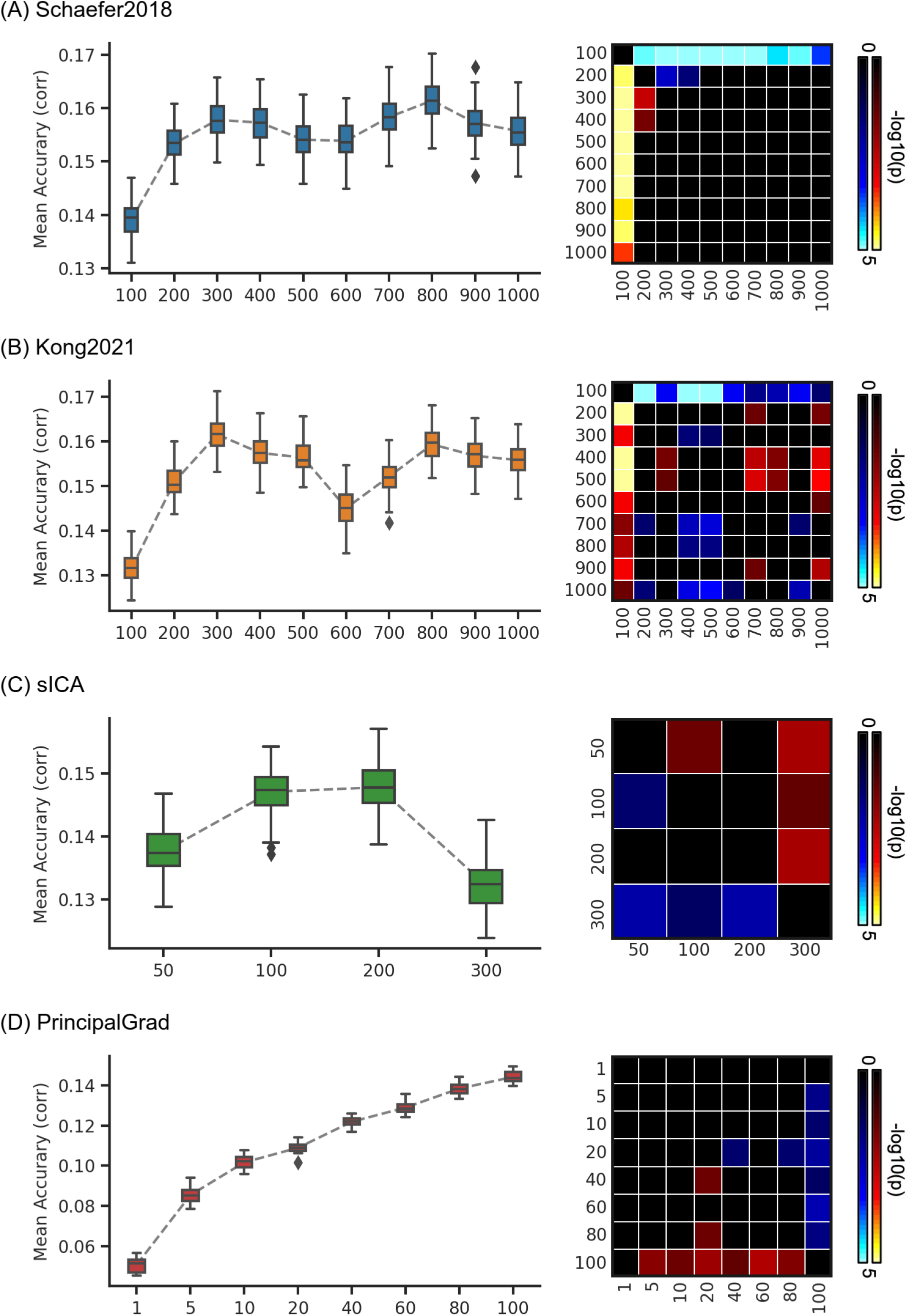
Average prediction accuracies (Pearson’s correlation) of task performance measures vary across resolutions for gradient and parcellation approaches using LRR in the HCP dataset. (A) Prediction accuracies and p values of the hard-parcellation Schaefer2018 with 100 to 1000 ROIs. (B) Prediction accuracies and p values of the hard-parcellation Kong2021 with 100 to 1000 ROIs. (C) Prediction accuracies and p values of the soft-parcellation sICA with 50 to 300 components. (D) Prediction accuracies and p values of the principal gradient PrincipalGrad with 1 to 100 gradients. Boxplots utilized default Python seaborn parameters, that is, box shows median and interquartile range (IQR). Whiskers indicate 1.5 IQR. P values (-log10(p)) were computed between prediction accuracies of each pair of resolutions. Non-black colors denote significantly different prediction performances after correcting for multiple comparisons with FDR q < 0.05. Bright colors indicate small p values, dark colors indicate large p values. For each pair of comparisons, warm colors represent higher prediction accuracies of the “row” resolution than the “column” resolution.

**Figure S25.**
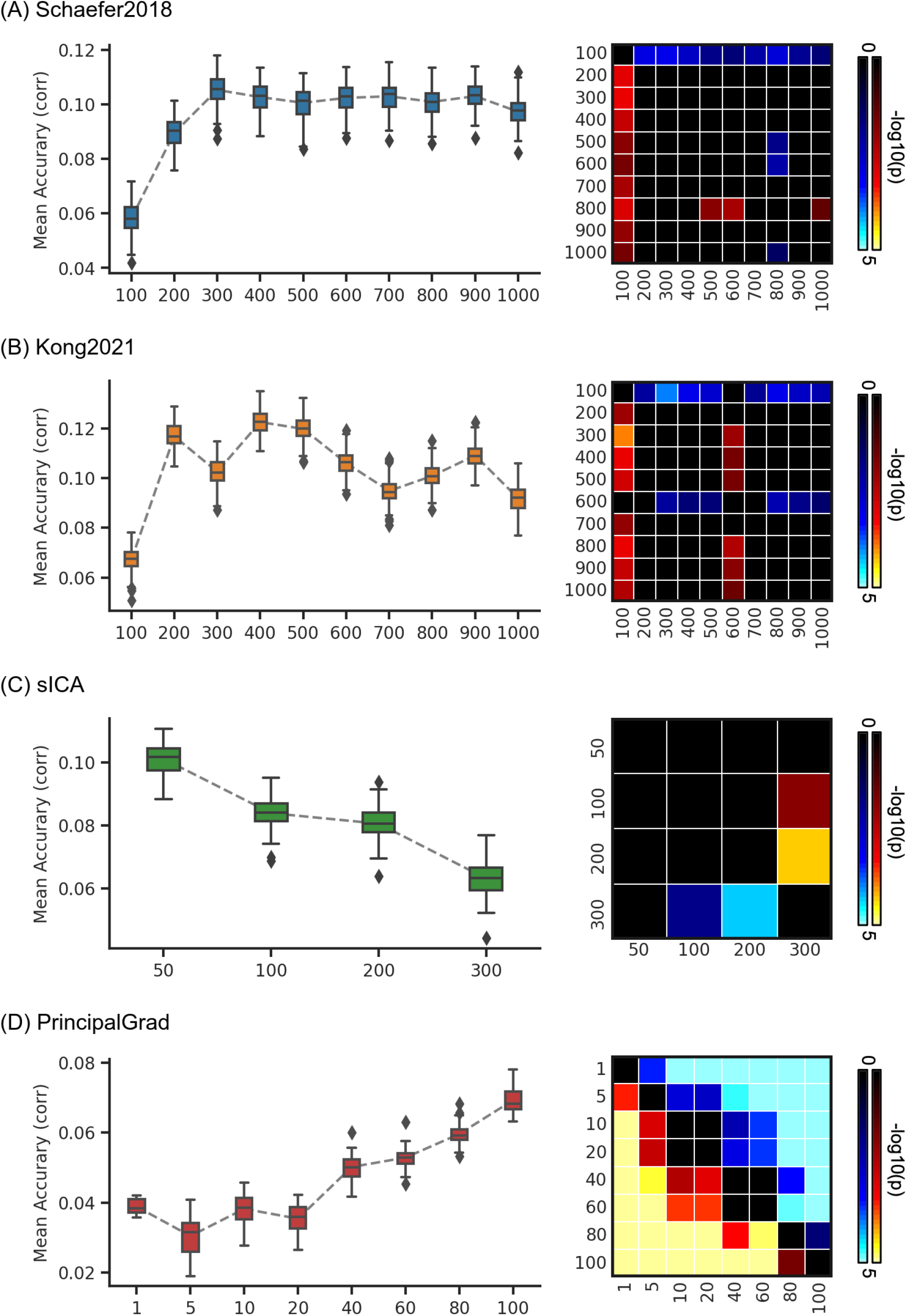
Average prediction accuracies (Pearson’s correlation) of self-reported measures vary across resolutions for gradient and parcellation approaches using LRR in the HCP dataset. (A) Prediction accuracies and p values of the hard-parcellation Schaefer2018 with 100 to 1000 ROIs. (B) Prediction accuracies and p values of the hard-parcellation Kong2021 with 100 to 1000 ROIs. (C) Prediction accuracies and p values of the soft-parcellation sICA with 50 to 300 components. (D) Prediction accuracies and p values of the principal gradient PrincipalGrad with 1 to 100 gradients. Boxplots utilized default Python seaborn parameters, that is, box shows median and interquartile range (IQR). Whiskers indicate 1.5 IQR. P values (-log10(p)) were computed between prediction accuracies of each pair of resolutions. Non-black colors denote significantly different prediction performances after correcting for multiple comparisons with FDR q < 0.05. Bright colors indicate small p values, dark colors indicate large p values. For each pair of comparisons, warm colors represent higher prediction accuracies of the “row” resolution than the “column” resolution.

**Figure S26.**
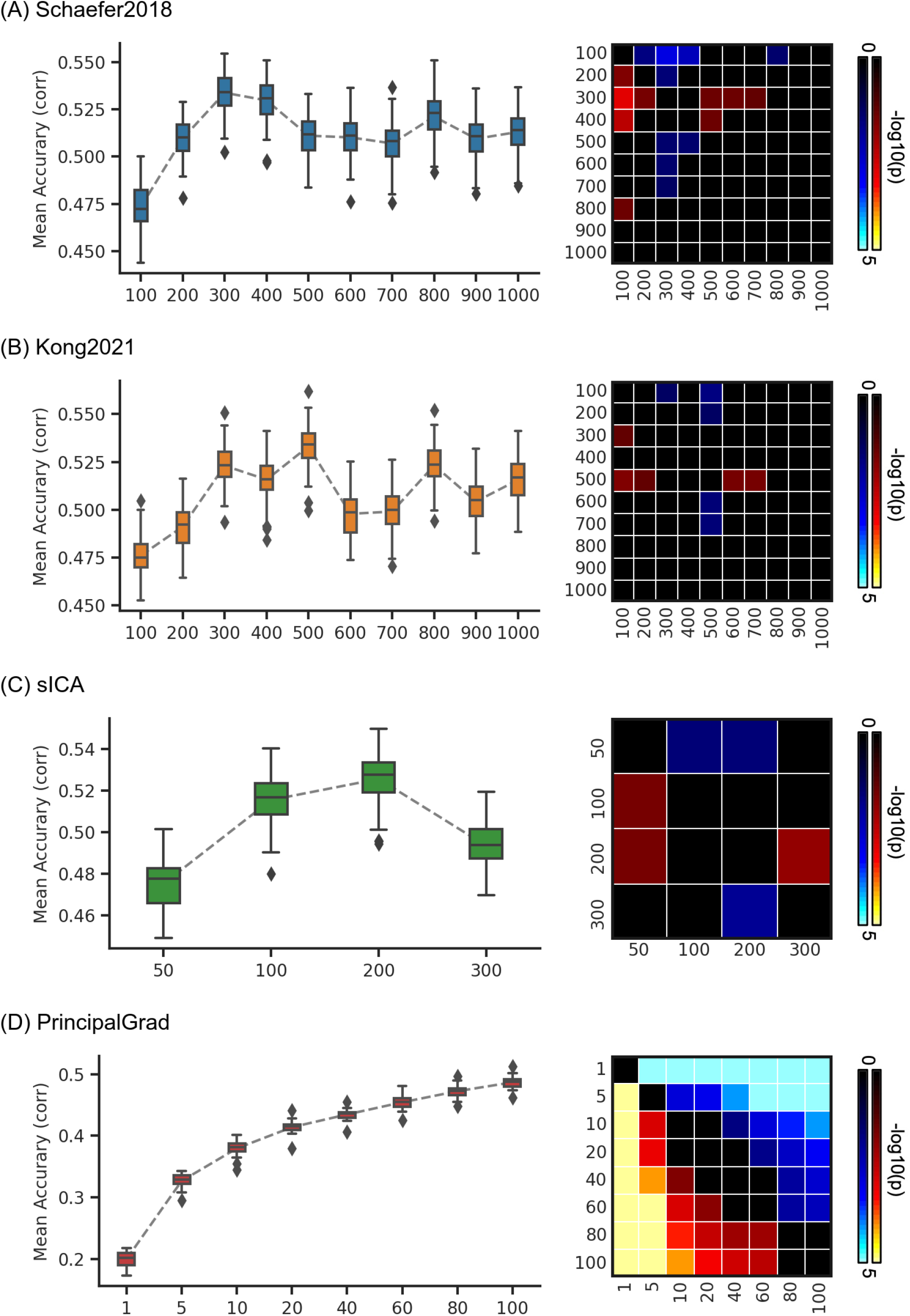
Prediction accuracies (Pearson’s correlation) of cognition vary across resolutions for gradient and parcellation approaches using LRR in the HCP dataset. (A) Prediction accuracies and p values of the hard-parcellation Schaefer2018 with 100 to 1000 ROIs. (B) Prediction accuracies and p values of the hard-parcellation Kong2021 with 100 to 1000 ROIs. (C) Prediction accuracies and p values of the soft-parcellation sICA with 50 to 300 components. (D) Prediction accuracies and p values of the principal gradient PrincipalGrad with 1 to 100 gradients. Boxplots utilized default Python seaborn parameters, that is, box shows median and interquartile range (IQR). Whiskers indicate 1.5 IQR. P values (-log10(p)) were computed between prediction accuracies of each pair of resolutions. Non-black colors denote significantly different prediction performances after correcting for multiple comparisons with FDR q < 0.05. Bright colors indicate small p values, dark colors indicate large p values. For each pair of comparisons, warm colors represent higher prediction accuracies of the “row” resolution than the “column” resolution.

**Figure S27.**
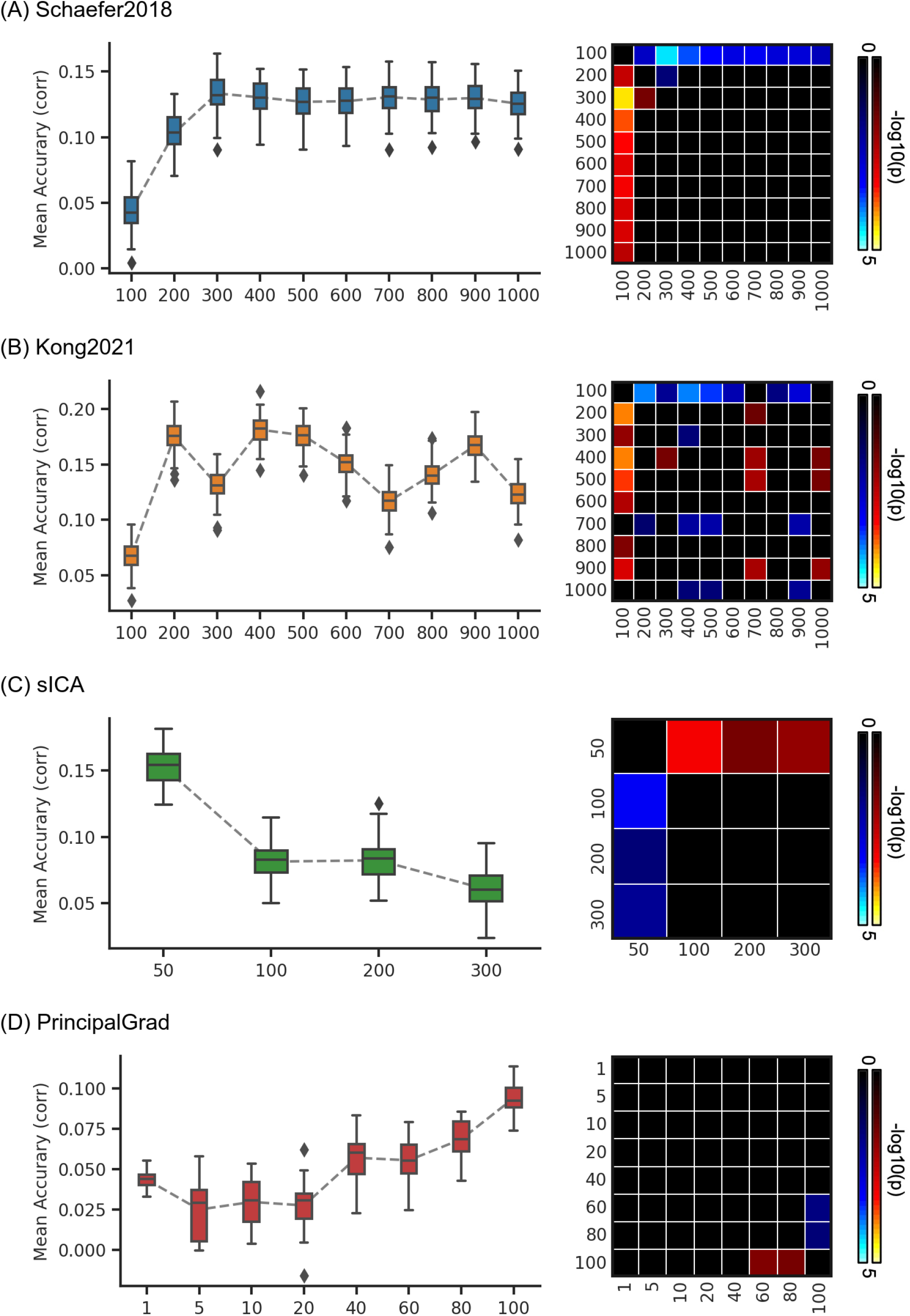
Prediction accuracies (Pearson’s correlation) of dissatisfaction vary across resolutions for gradient and parcellation approaches using LRR in the HCP dataset. (A) Prediction accuracies and p values of the hard-parcellation Schaefer2018 with 100 to 1000 ROIs. (B) Prediction accuracies and p values of the hard-parcellation Kong2021 with 100 to 1000 ROIs. (C) Prediction accuracies and p values of the soft-parcellation sICA with 50 to 300 components. (D) Prediction accuracies and p values of the principal gradient PrincipalGrad with 1 to 100 gradients. Boxplots utilized default Python seaborn parameters, that is, box shows median and interquartile range (IQR). Whiskers indicate 1.5 IQR. P values (-log10(p)) were computed between prediction accuracies of each pair of resolutions. Non-black colors denote significantly different prediction performances after correcting for multiple comparisons with FDR q < 0.05. Bright colors indicate small p values, dark colors indicate large p values. For each pair of comparisons, warm colors represent higher prediction accuracies of the “row” resolution than the “column” resolution.

**Figure S28.**
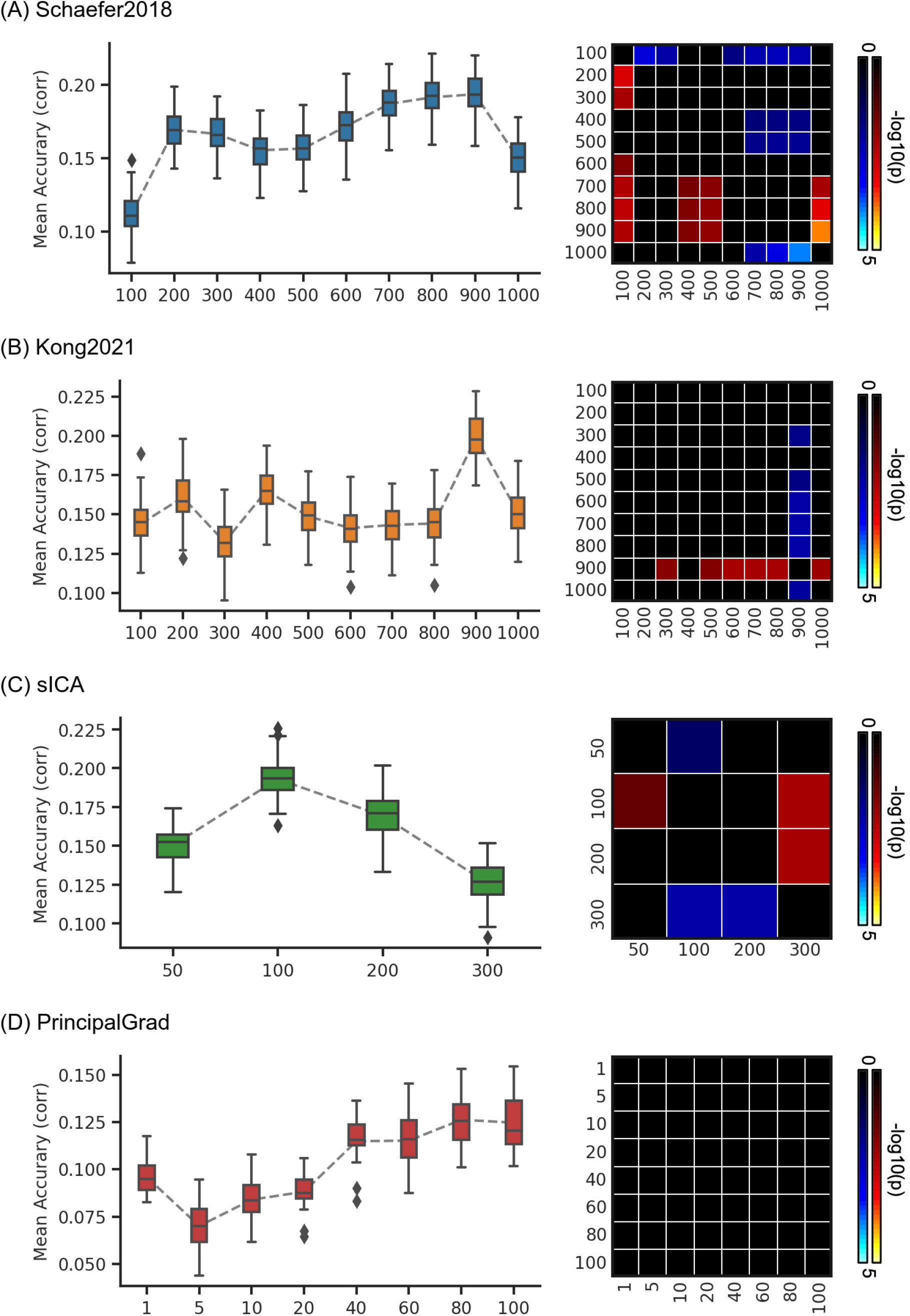
Prediction accuracies (Pearson’s correlation) of emotion vary across resolutions for gradient and parcellation approaches using LRR in the HCP dataset. (A) Prediction accuracies and p values of the hard-parcellation Schaefer2018 with 100 to 1000 ROIs. (B) Prediction accuracies and p values of the hard-parcellation Kong2021 with 100 to 1000 ROIs. (C) Prediction accuracies and p values of the soft-parcellation sICA with 50 to 300 components. (D) Prediction accuracies and p values of the principal gradient PrincipalGrad with 1 to 100 gradients. Boxplots utilized default Python seaborn parameters, that is, box shows median and interquartile range (IQR). Whiskers indicate 1.5 IQR. P values (-log10(p)) were computed between prediction accuracies of each pair of resolutions. Non-black colors denote significantly different prediction performances after correcting for multiple comparisons with FDR q < 0.05. Bright colors indicate small p values, dark colors indicate large p values. For each pair of comparisons, warm colors represent higher prediction accuracies of the “row” resolution than the “column” resolution.

**Figure S29.**
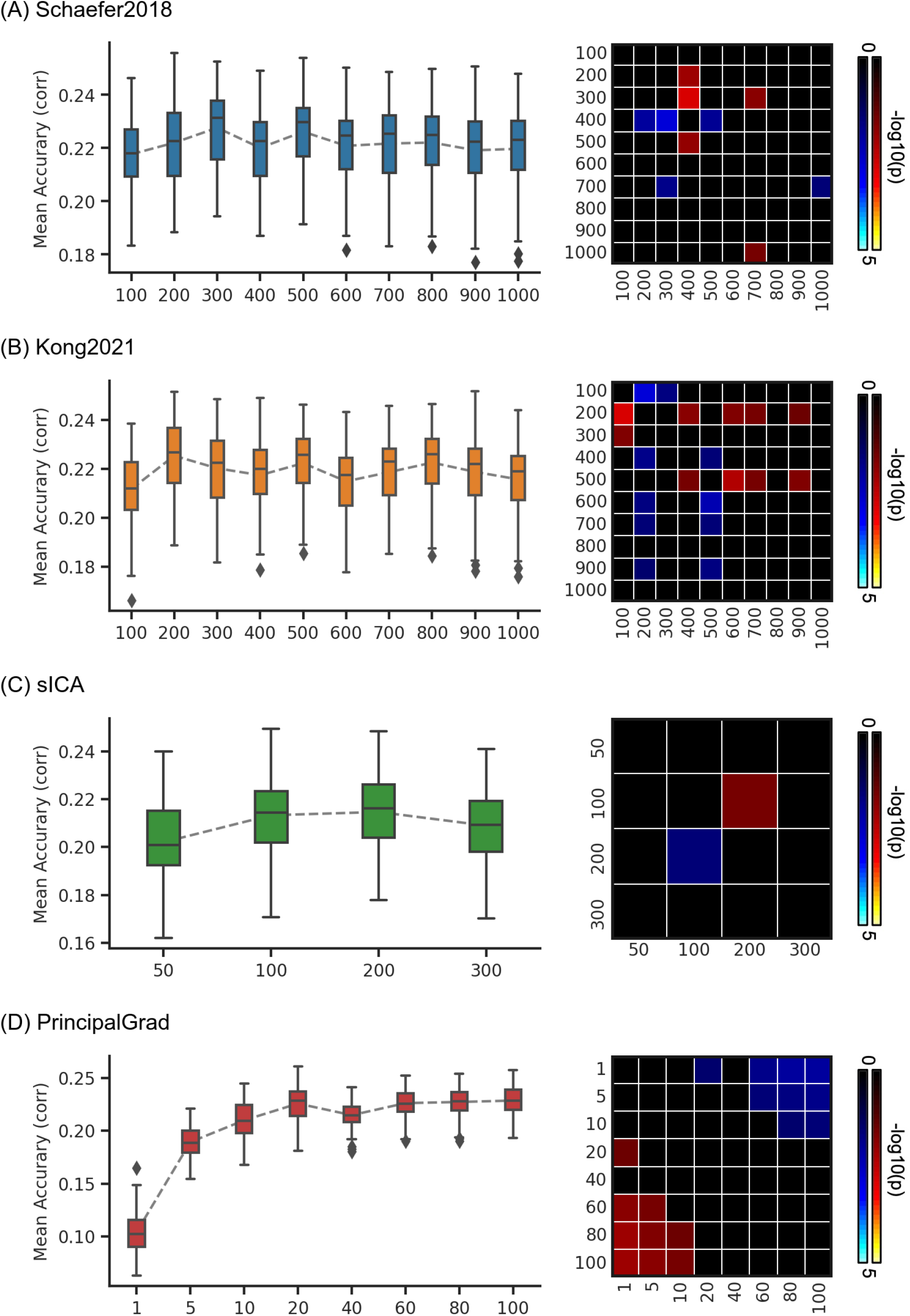
Average prediction accuracies (Pearson’s correlation) of task performance measures vary across resolutions for gradient and parcellation approaches using KRR in the ABCD dataset. (A) Prediction accuracies and p values of the hard-parcellation Schaefer2018 with 100 to 1000 ROIs. (B) Prediction accuracies and p values of the hard-parcellation Kong2021 with 100 to 1000 ROIs. (C) Prediction accuracies and p values of the soft-parcellation sICA with 50 to 300 components. (D) Prediction accuracies and p values of the principal gradient PrincipalGrad with 1 to 100 gradients. Boxplots utilized default Python seaborn parameters, that is, box shows median and interquartile range (IQR). Whiskers indicate 1.5 IQR. P values (-log10(p)) were computed between prediction accuracies of each pair of resolutions. Non-black colors denote significantly different prediction performances after correcting for multiple comparisons with FDR q < 0.05. Bright colors indicate small p values, dark colors indicate large p values. For each pair of comparisons, warm colors represent higher prediction accuracies of the “row” resolution than the “column” resolution.

**Figure S30.**
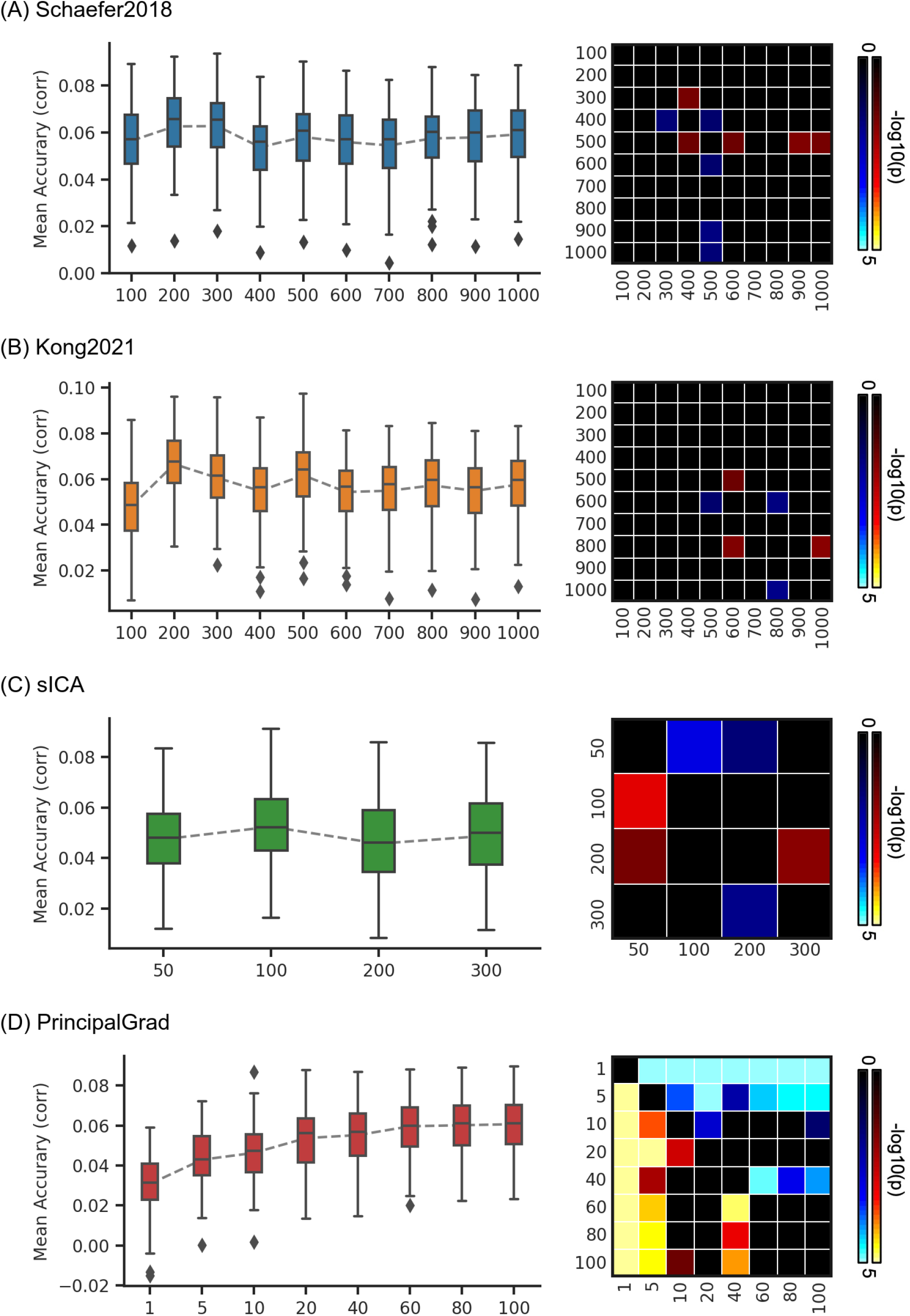
Average prediction accuracies (Pearson’s correlation) of self-reported measures vary across resolutions for gradient and parcellation approaches using KRR in the ABCD dataset. (A) Prediction accuracies and p values of the hard-parcellation Schaefer2018 with 100 to 1000 ROIs. (B) Prediction accuracies and p values of the hard-parcellation Kong2021 with 100 to 1000 ROIs. (C) Prediction accuracies and p values of the soft-parcellation sICA with 50 to 300 components. (D) Prediction accuracies and p values of the principal gradient PrincipalGrad with 1 to 100 gradients. Boxplots utilized default Python seaborn parameters, that is, box shows median and interquartile range (IQR). Whiskers indicate 1.5 IQR. P values (-log10(p)) were computed between prediction accuracies of each pair of resolutions. Non-black colors denote significantly different prediction performances after correcting for multiple comparisons with FDR q < 0.05. Bright colors indicate small p values, dark colors indicate large p values. For each pair of comparisons, warm colors represent higher prediction accuracies of the “row” resolution than the “column” resolution.

**Figure S31.**
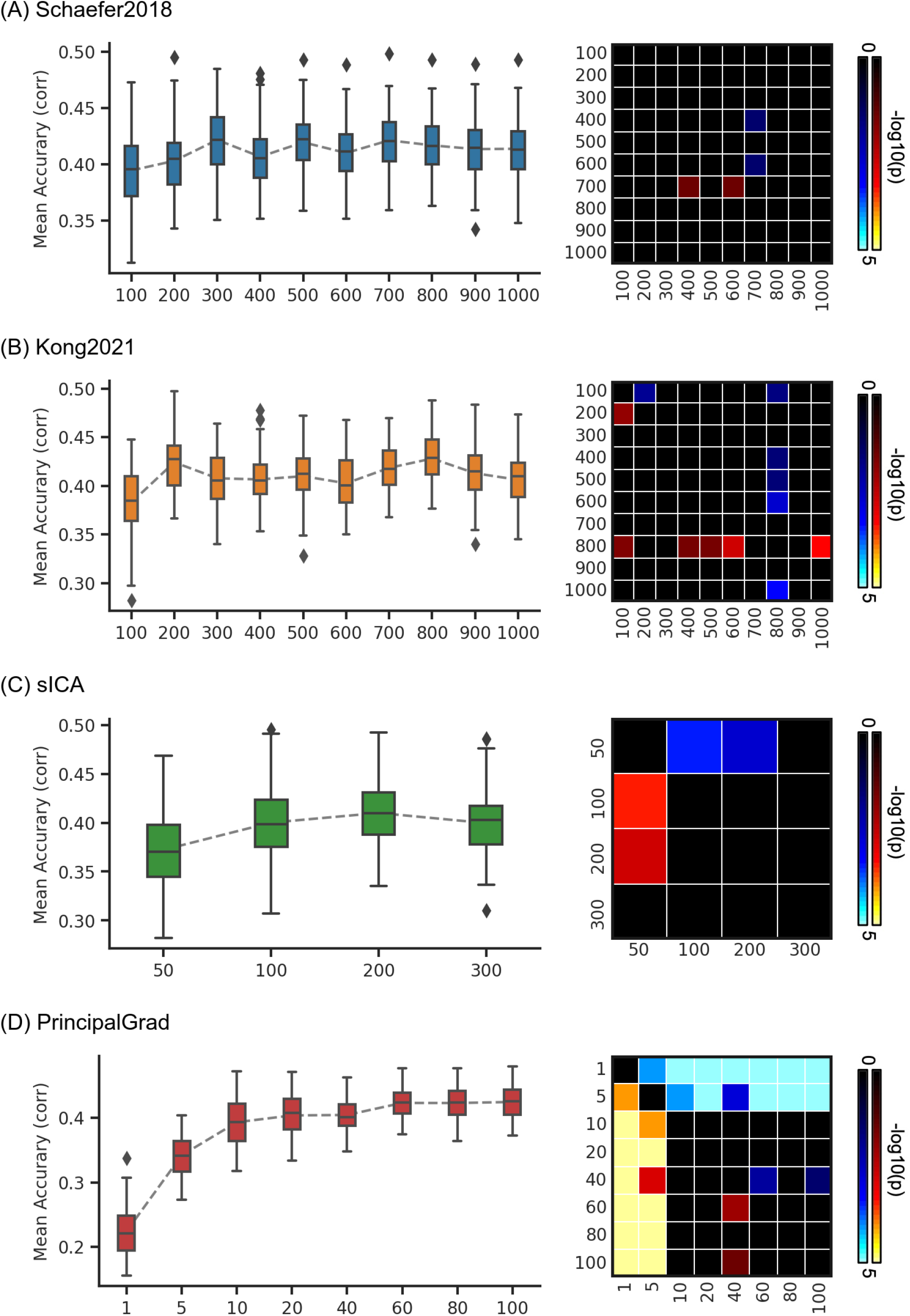
Prediction accuracies (Pearson’s correlation) of cognition vary across resolutions for gradient and parcellation approaches using KRR in the ABCD dataset. (A) Prediction accuracies and p values of the hard-parcellation Schaefer2018 with 100 to 1000 ROIs. (B) Prediction accuracies and p values of the hard-parcellation Kong2021 with 100 to 1000 ROIs. (C) Prediction accuracies and p values of the soft-parcellation sICA with 50 to 300 components. (D) Prediction accuracies and p values of the principal gradient PrincipalGrad with 1 to 100 gradients. Boxplots utilized default Python seaborn parameters, that is, box shows median and interquartile range (IQR). Whiskers indicate 1.5 IQR. P values (-log10(p)) were computed between prediction accuracies of each pair of resolutions. Non-black colors denote significantly different prediction performances after correcting for multiple comparisons with FDR q < 0.05. Bright colors indicate small p values, dark colors indicate large p values. For each pair of comparisons, warm colors represent higher prediction accuracies of the “row” resolution than the “column” resolution.

**Figure S32.**
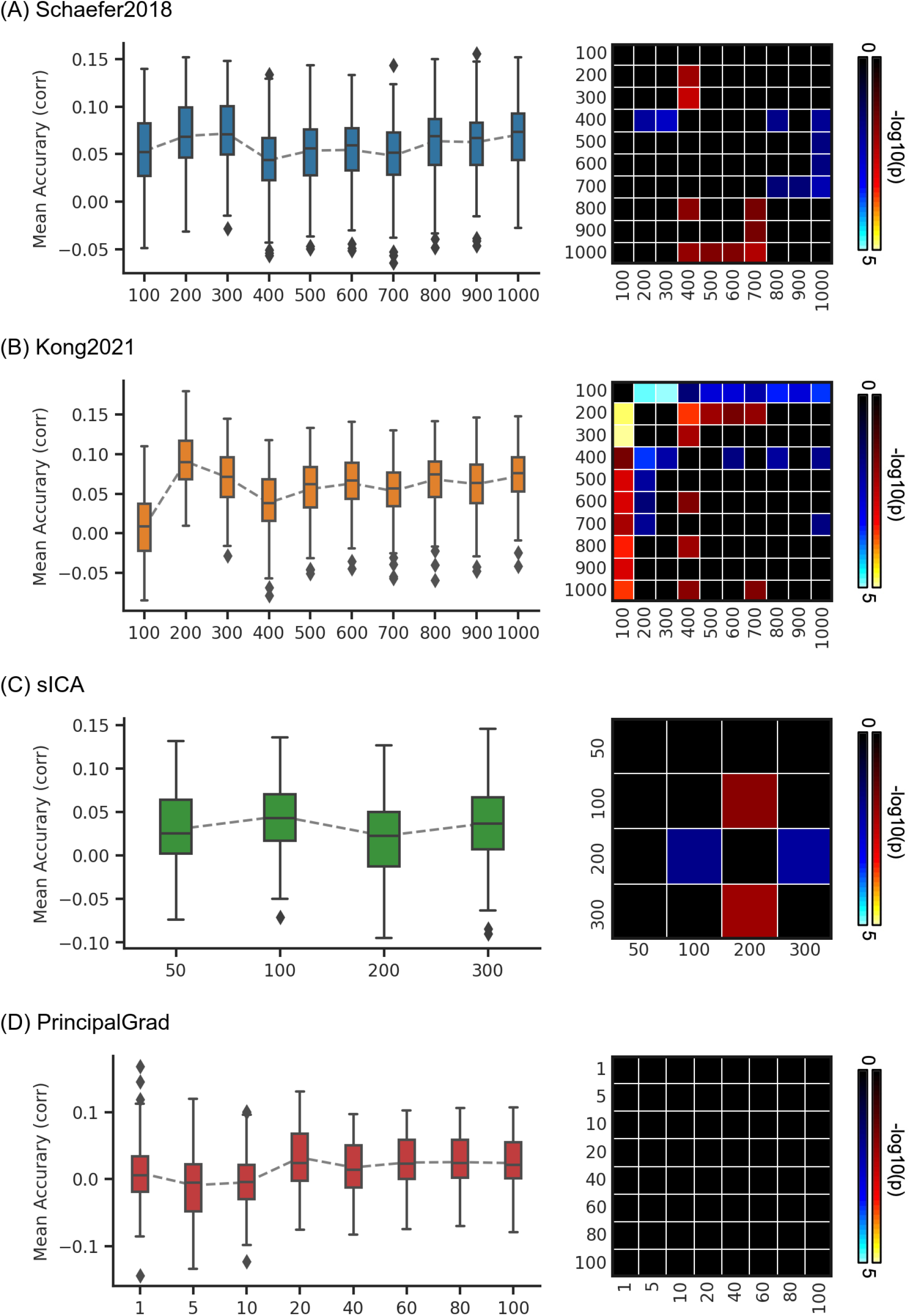
Prediction accuracies (Pearson’s correlation) of mental health vary across resolutions for gradient and parcellation approaches using KRR in the ABCD dataset. (A) Prediction accuracies and p values of the hard-parcellation Schaefer2018 with 100 to 1000 ROIs. (B) Prediction accuracies and p values of the hard-parcellation Kong2021 with 100 to 1000 ROIs. (C) Prediction accuracies and p values of the soft-parcellation sICA with 50 to 300 components. (D) Prediction accuracies and p values of the principal gradient PrincipalGrad with 1 to 100 gradients. Boxplots utilized default Python seaborn parameters, that is, box shows median and interquartile range (IQR). Whiskers indicate 1.5 IQR. P values (-log10(p)) were computed between prediction accuracies of each pair of resolutions. Non-black colors denote significantly different prediction performances after correcting for multiple comparisons with FDR q < 0.05. Bright colors indicate small p values, dark colors indicate large p values. For each pair of comparisons, warm colors represent higher prediction accuracies of the “row” resolution than the “column” resolution.

**Figure S33.**
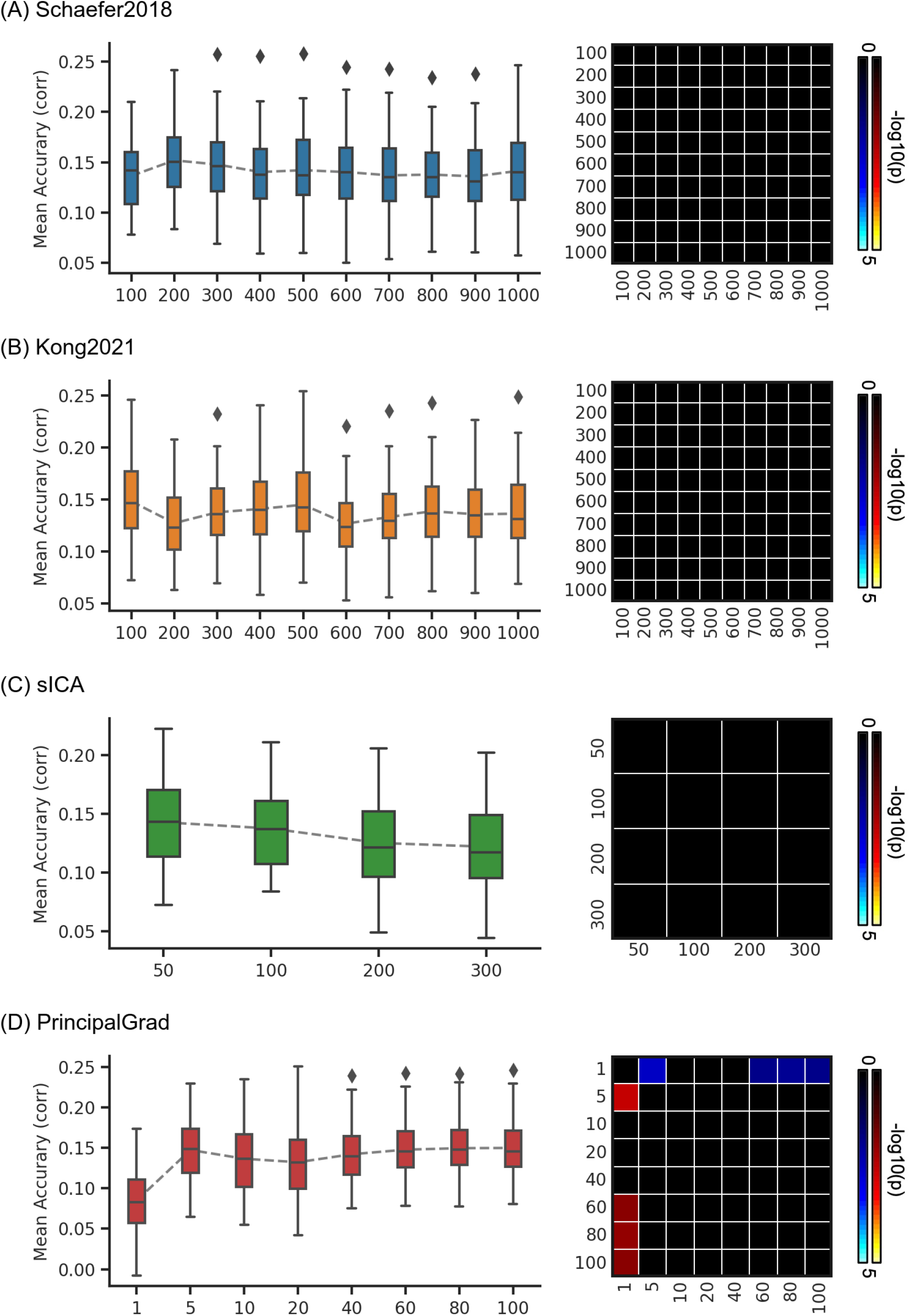
Prediction accuracies (Pearson’s correlation) of personality vary across resolutions for gradient and parcellation approaches using KRR in the ABCD dataset. (A) Prediction accuracies and p values of the hard-parcellation Schaefer2018 with 100 to 1000 ROIs. (B) Prediction accuracies and p values of the hard-parcellation Kong2021 with 100 to 1000 ROIs. Prediction accuracies and p values of the soft-parcellation sICA with 50 to 300 components. Prediction accuracies and p values of the principal gradient PrincipalGrad with 1 to 100 gradients. Boxplots utilized default Python seaborn parameters, that is, box shows median and interquartile range (IQR). Whiskers indicate 1.5 IQR. P values (-log10(p)) were computed between prediction accuracies of each pair of resolutions. Non-black colors denote significantly different prediction performances after correcting for multiple comparisons with FDR q < 0.05. Bright colors indicate small p values, dark colors indicate large p values. For each pair of comparisons, warm colors represent higher prediction accuracies of the “row” resolution than the “column” resolution.

**Figure S34.**
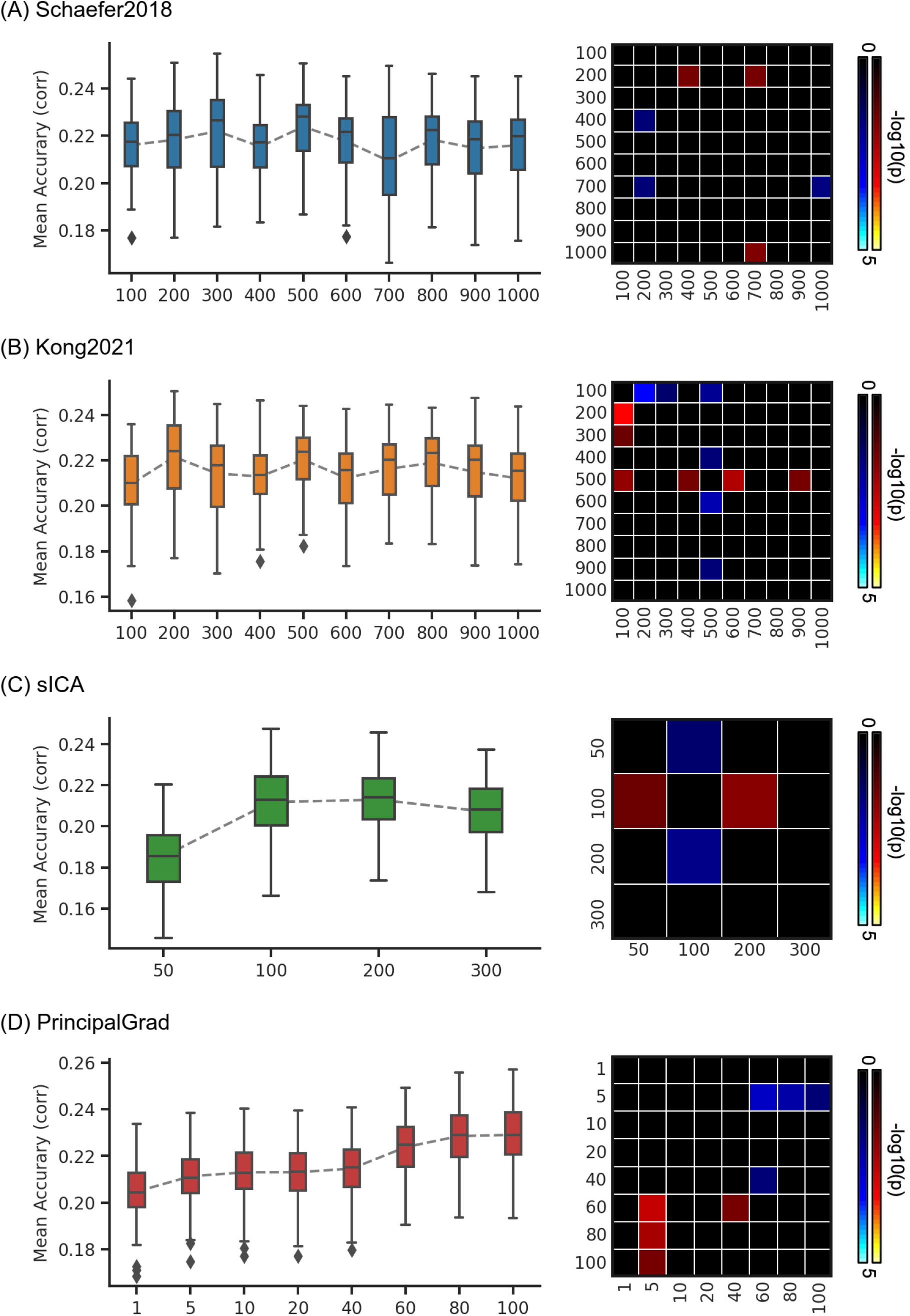
Average prediction accuracies (Pearson’s correlation) of task performance measures vary across resolutions for gradient and parcellation approaches using LRR in the ABCD dataset. (A) Prediction accuracies and p values of the hard-parcellation Schaefer2018 with 100 to 1000 ROIs. (B) Prediction accuracies and p values of the hard-parcellation Kong2021 with 100 to 1000 ROIs. (C) Prediction accuracies and p values of the soft-parcellation sICA with 50 to 300 components. (D) Prediction accuracies and p values of the principal gradient PrincipalGrad with 1 to 100 gradients. Boxplots utilized default Python seaborn parameters, that is, box shows median and interquartile range (IQR). Whiskers indicate 1.5 IQR. P values (-log10(p)) were computed between prediction accuracies of each pair of resolutions. Non-black colors denote significantly different prediction performances after correcting for multiple comparisons with FDR q < 0.05. Bright colors indicate small p values, dark colors indicate large p values. For each pair of comparisons, warm colors represent higher prediction accuracies of the “row” resolution than the “column” resolution.

**Figure S35.**
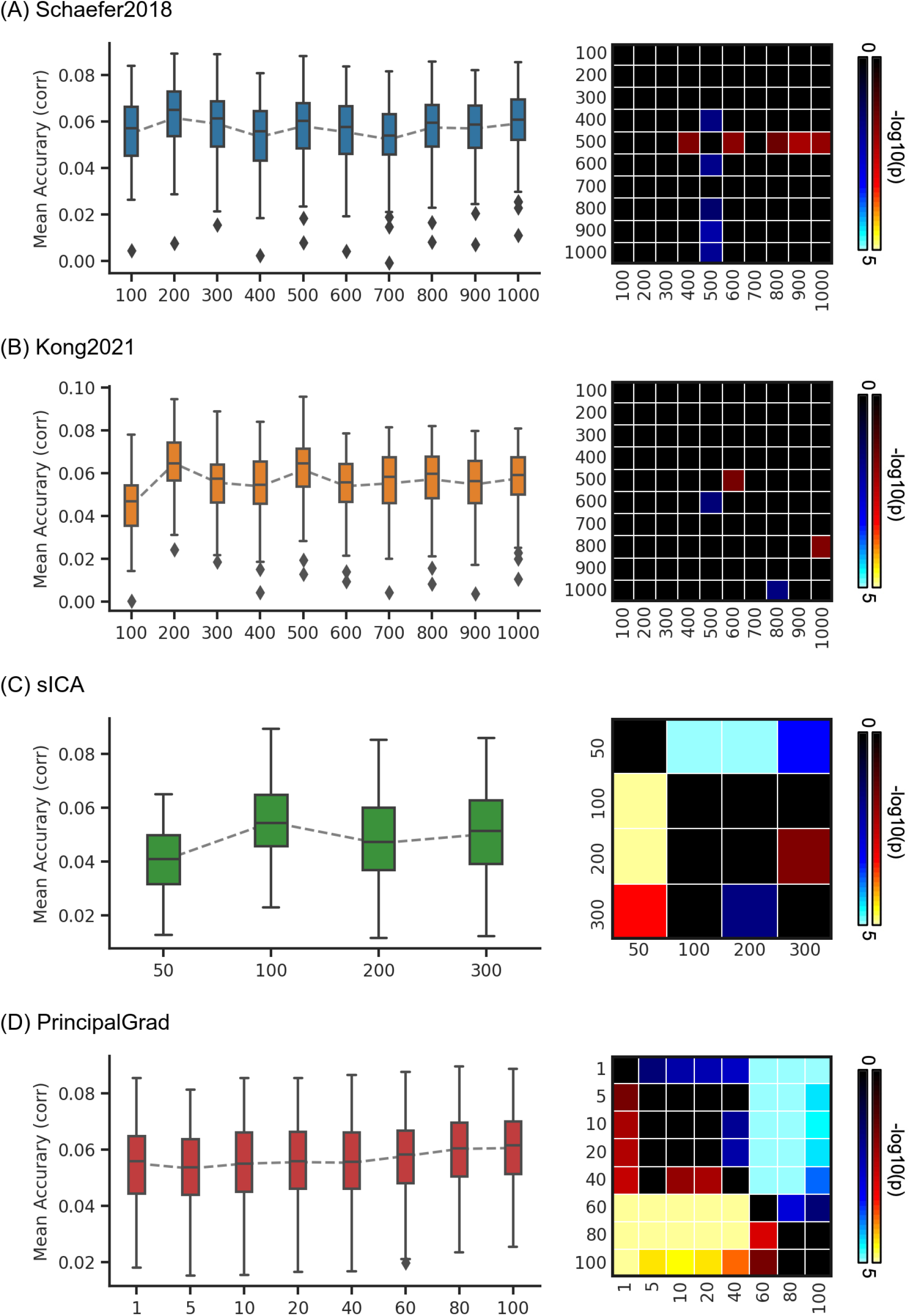
Average prediction accuracies (Pearson’s correlation) of self-reported measures vary across resolutions for gradient and parcellation approaches using LRR in the ABCD dataset. (A) Prediction accuracies and p values of the hard-parcellation Schaefer2018 with 100 to 1000 ROIs. Prediction accuracies and p values of the hard-parcellation Kong2021 with 100 to 1000 ROIs. (C) Prediction accuracies and p values of the soft-parcellation sICA with 50 to 300 components. (D) Prediction accuracies and p values of the principal gradient PrincipalGrad with 1 to 100 gradients. Boxplots utilized default Python seaborn parameters, that is, box shows median and interquartile range (IQR). Whiskers indicate 1.5 IQR. P values (-log10(p)) were computed between prediction accuracies of each pair of resolutions. Non-black colors denote significantly different prediction performances after correcting for multiple comparisons with FDR q < 0.05. Bright colors indicate small p values, dark colors indicate large p values. For each pair of comparisons, warm colors represent higher prediction accuracies of the “row” resolution than the “column” resolution.

**Figure S36.**
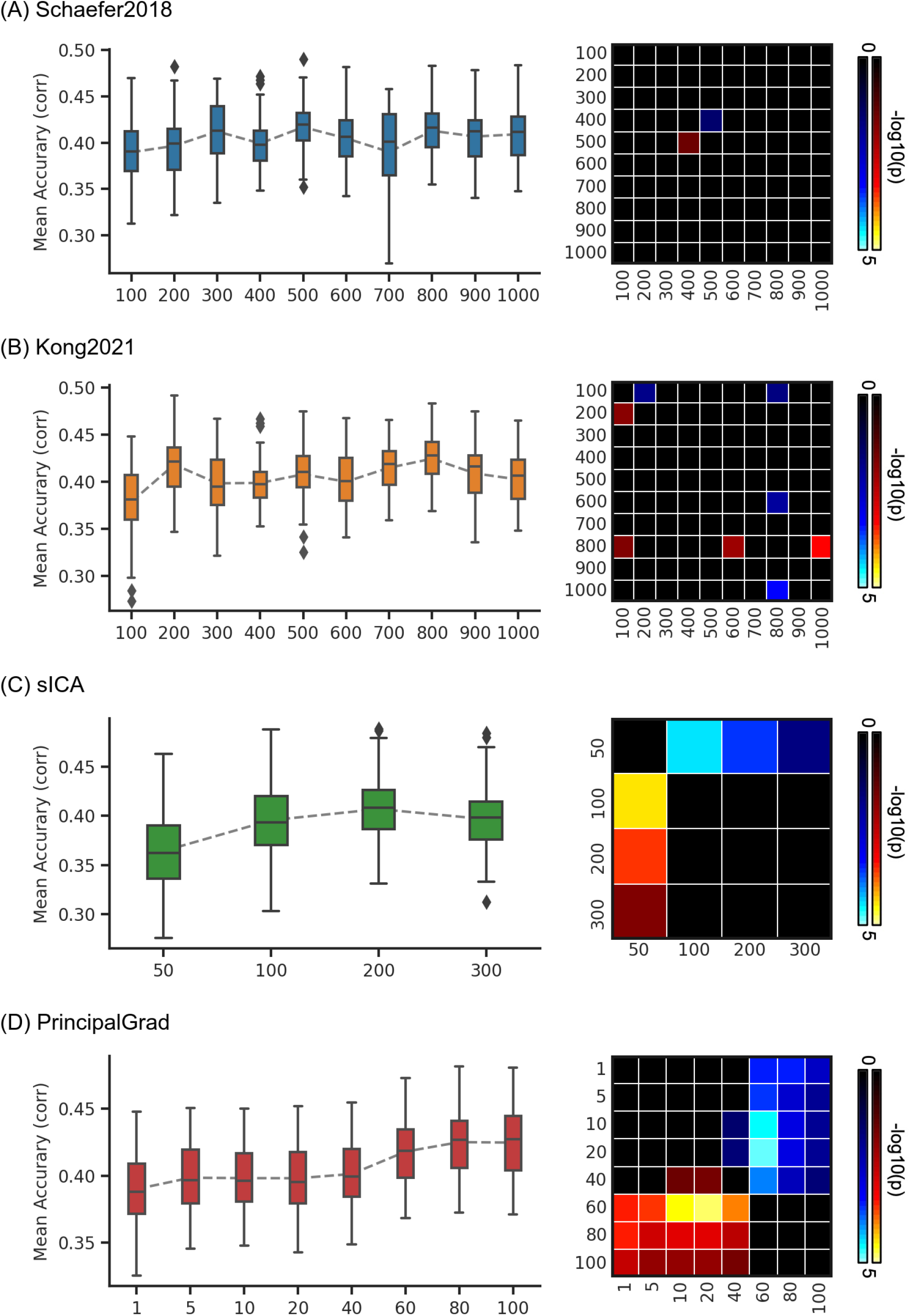
Prediction accuracies (Pearson’s correlation) of cognition vary across resolutions for gradient and parcellation approaches using LRR in the ABCD dataset. (A) Prediction accuracies and p values of the hard-parcellation Schaefer2018 with 100 to 1000 ROIs. (B) Prediction accuracies and p values of the hard-parcellation Kong2021 with 100 to 1000 ROIs. (C) Prediction accuracies and p values of the soft-parcellation sICA with 50 to 300 components. (D) Prediction accuracies and p values of the principal gradient PrincipalGrad with 1 to 100 gradients. Boxplots utilized default Python seaborn parameters, that is, box shows median and interquartile range (IQR). Whiskers indicate 1.5 IQR. P values (-log10(p)) were computed between prediction accuracies of each pair of resolutions. Non-black colors denote significantly different prediction performances after correcting for multiple comparisons with FDR q < 0.05. Bright colors indicate small p values, dark colors indicate large p values. For each pair of comparisons, warm colors represent higher prediction accuracies of the “row” resolution than the “column” resolution.

**Figure S37.**
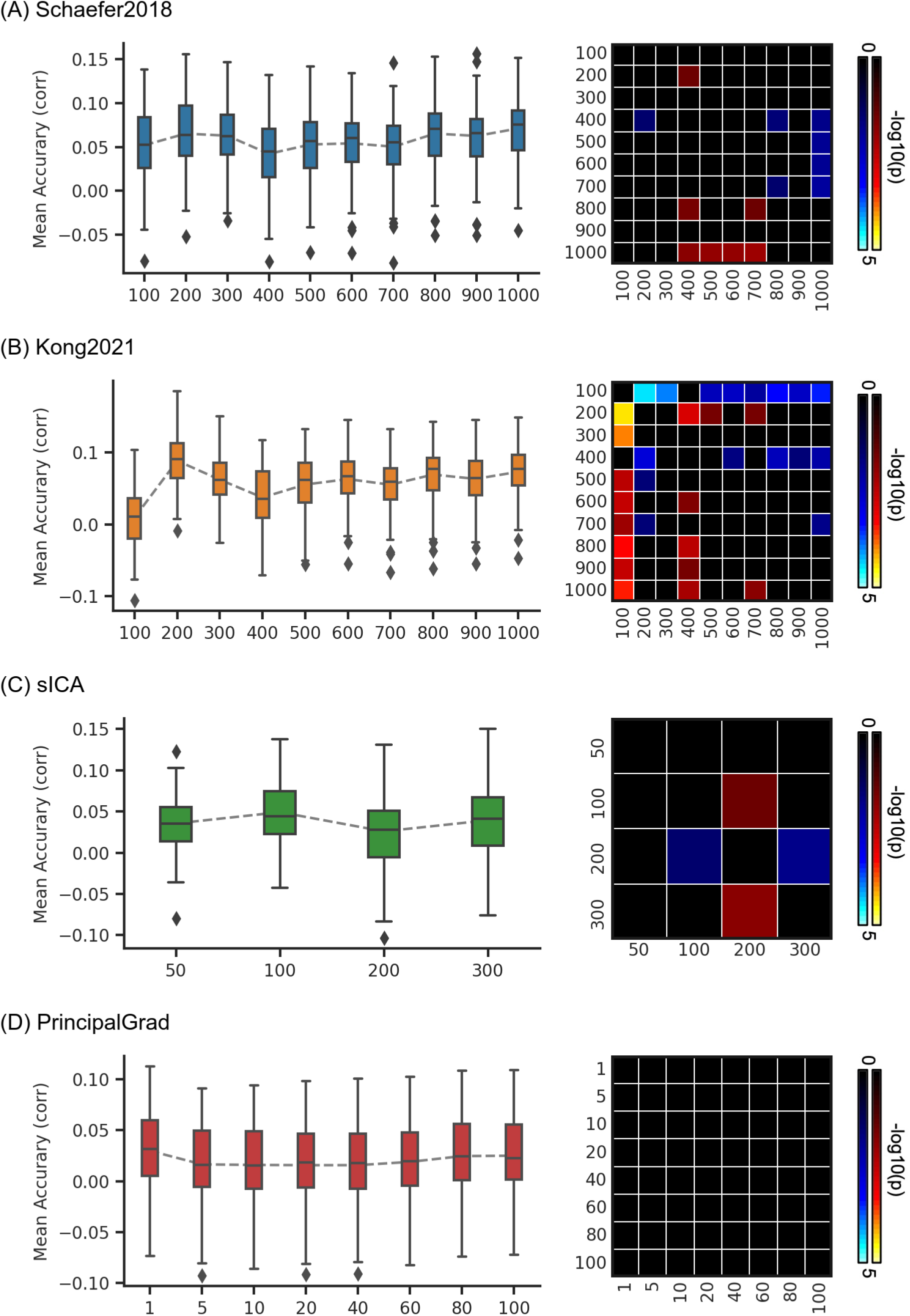
Prediction accuracies (Pearson’s correlation) of mental health vary across resolutions for gradient and parcellation approaches using LRR in the ABCD dataset. (A) Prediction accuracies and p values of the hard-parcellation Schaefer2018 with 100 to 1000 ROIs. (B) Prediction accuracies and p values of the hard-parcellation Kong2021 with 100 to 1000 ROIs. Prediction accuracies and p values of the soft-parcellation sICA with 50 to 300 components. Prediction accuracies and p values of the principal gradient PrincipalGrad with 1 to 100 gradients. Boxplots utilized default Python seaborn parameters, that is, box shows median and interquartile range (IQR). Whiskers indicate 1.5 IQR. P values (-log10(p)) were computed between prediction accuracies of each pair of resolutions. Non-black colors denote significantly different prediction performances after correcting for multiple comparisons with FDR q < 0.05. Bright colors indicate small p values, dark colors indicate large p values. For each pair of comparisons, warm colors represent higher prediction accuracies of the “row” resolution than the “column” resolution.

**Figure S38.**
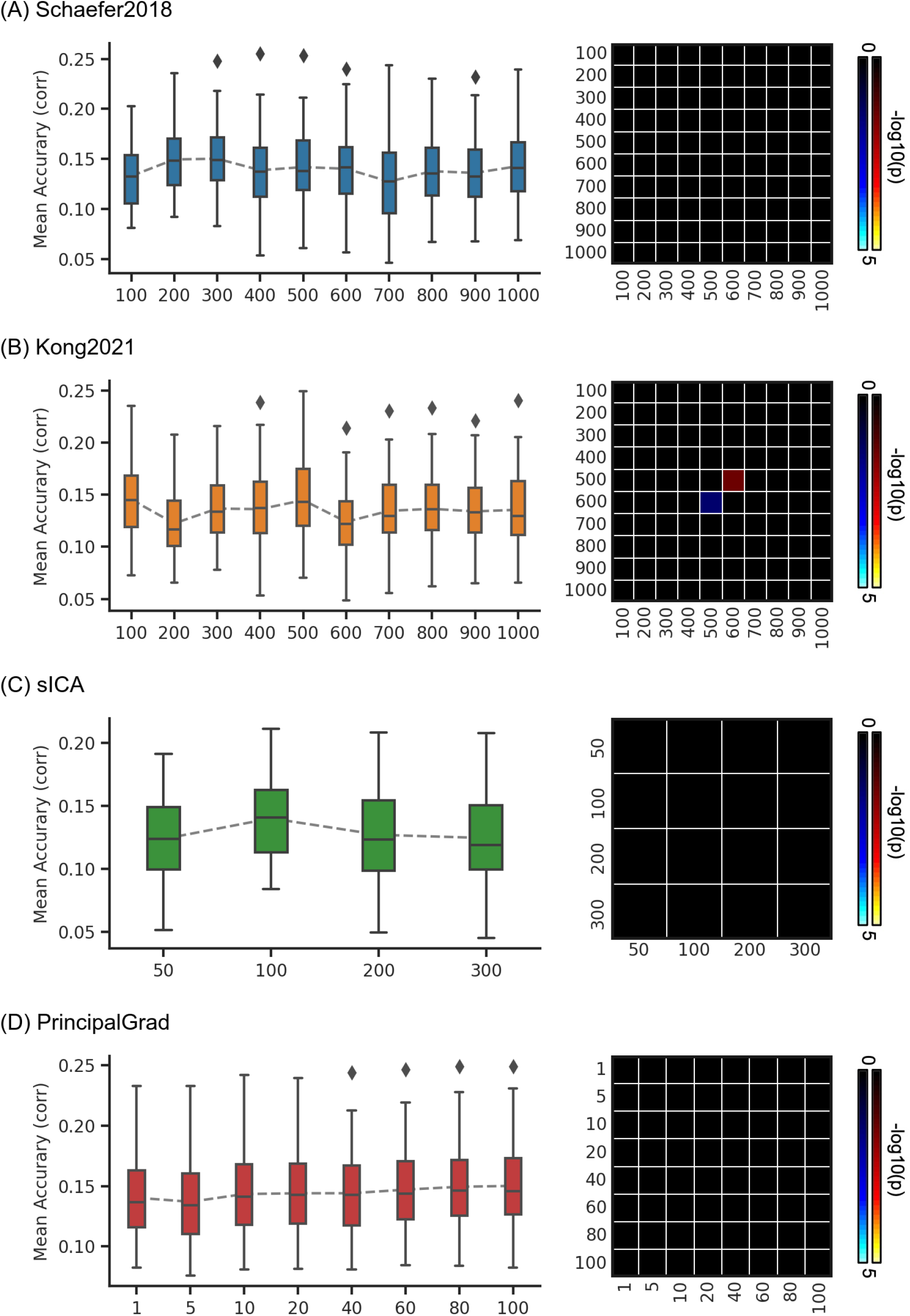
Prediction accuracies (Pearson’s correlation) of emotion vary across resolutions for gradient and parcellation approaches using LRR in the ABCD dataset. (A) Prediction accuracies and p values of the hard-parcellation Schaefer2018 with 100 to 1000 ROIs. (B) Prediction accuracies and p values of the hard-parcellation Kong2021 with 100 to 1000 ROIs. (C) Prediction accuracies and p values of the soft-parcellation sICA with 50 to 300 components. (D) Prediction accuracies and p values of the principal gradient PrincipalGrad with 1 to 100 gradients. Boxplots utilized default Python seaborn parameters, that is, box shows median and interquartile range (IQR). Whiskers indicate 1.5 IQR. P values (-log10(p)) were computed between prediction accuracies of each pair of resolutions. Non-black colors denote significantly different prediction performances after correcting for multiple comparisons with FDR q < 0.05. Bright colors indicate small p values, dark colors indicate large p values. For each pair of comparisons, warm colors represent higher prediction accuracies of the “row” resolution than the “column” resolution.

**Figure S39.**
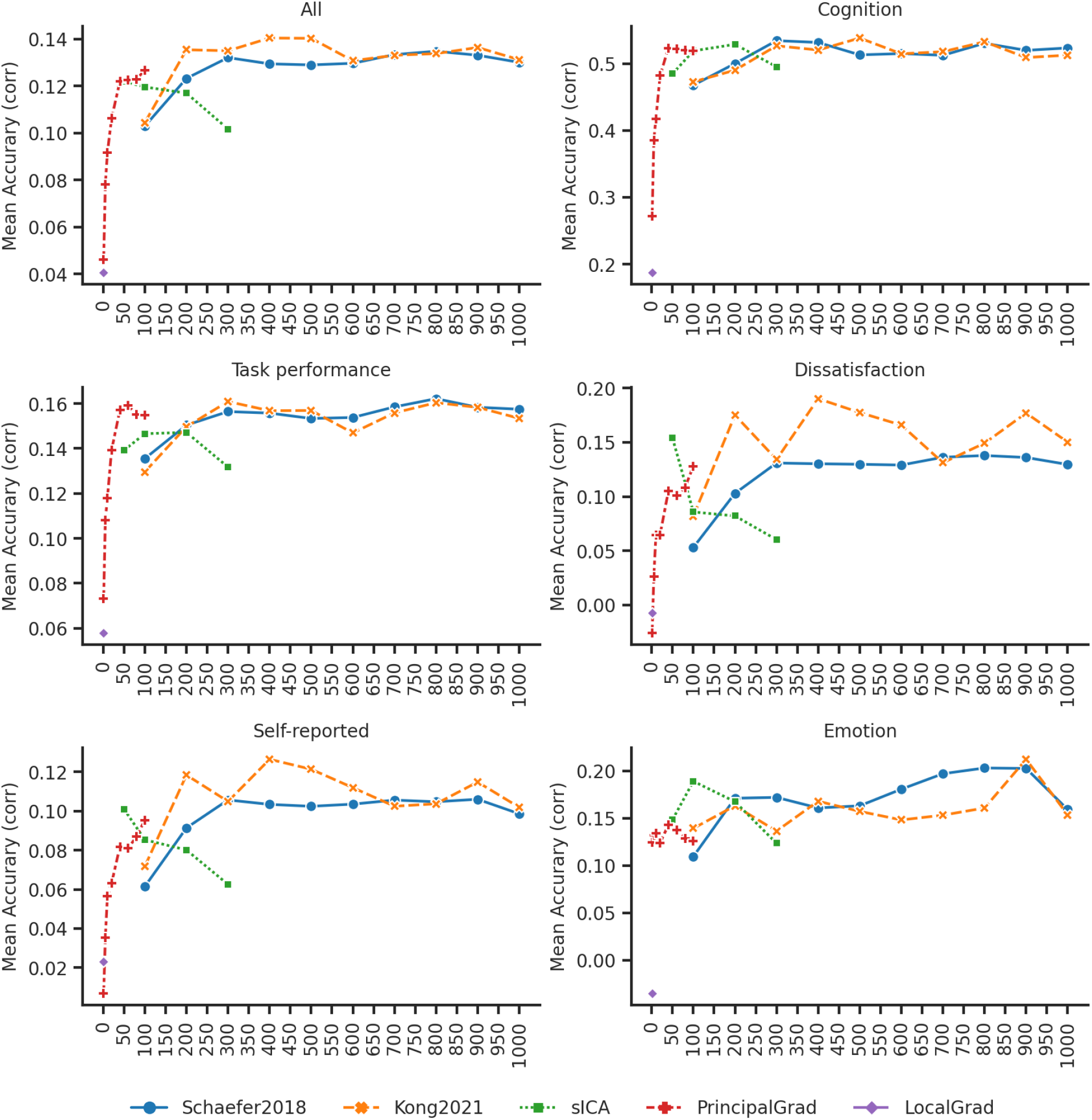
Prediction accuracies (Pearson’s correlation) vary across resolutions for gradient and parcellation approaches using KRR in the HCP dataset.

**Figure S40.**
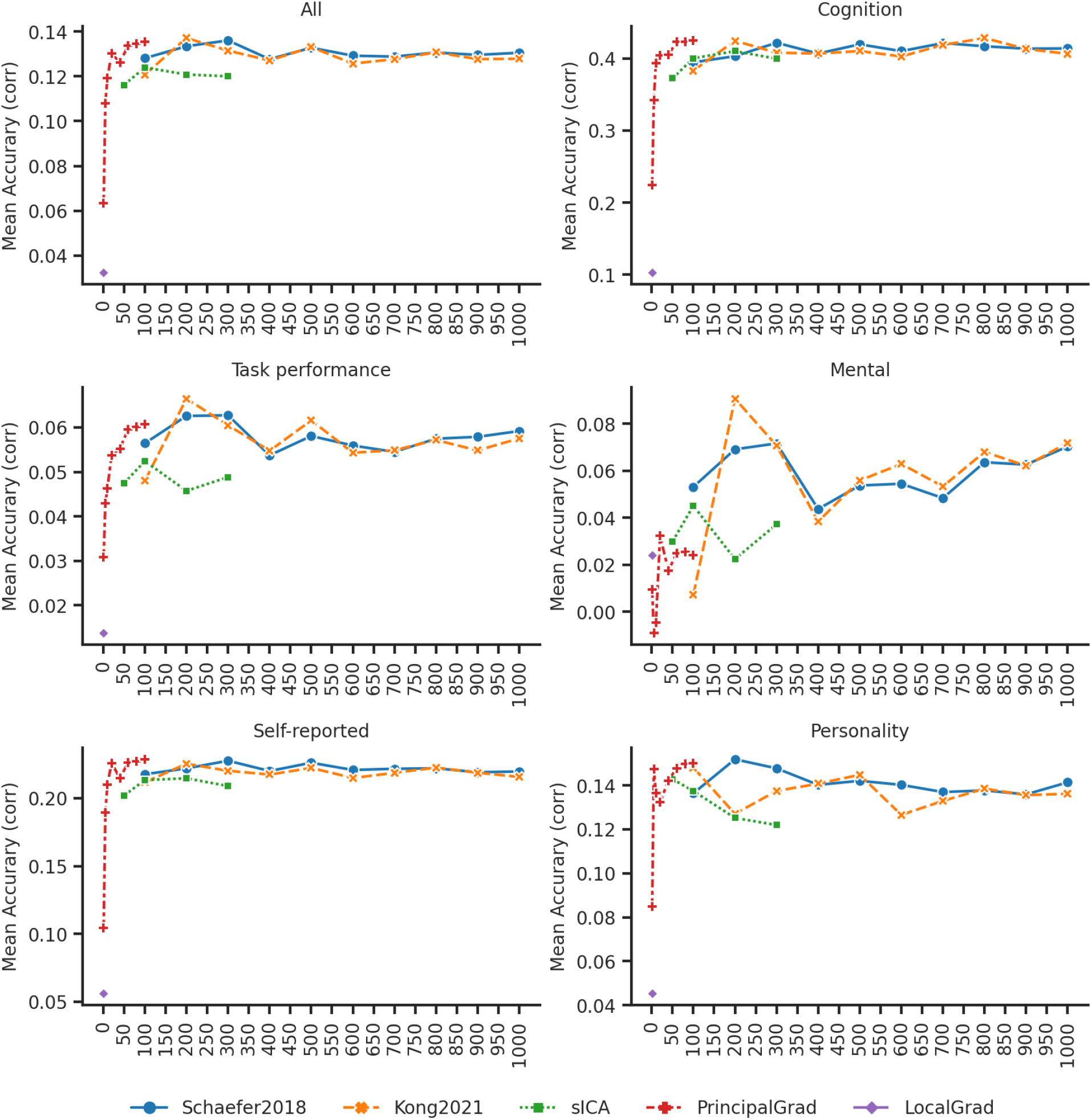
Prediction accuracies (Pearson’s correlation) vary across resolutions for gradient and parcellation approaches using KRR in the ABCD dataset.

**Figure S41.**
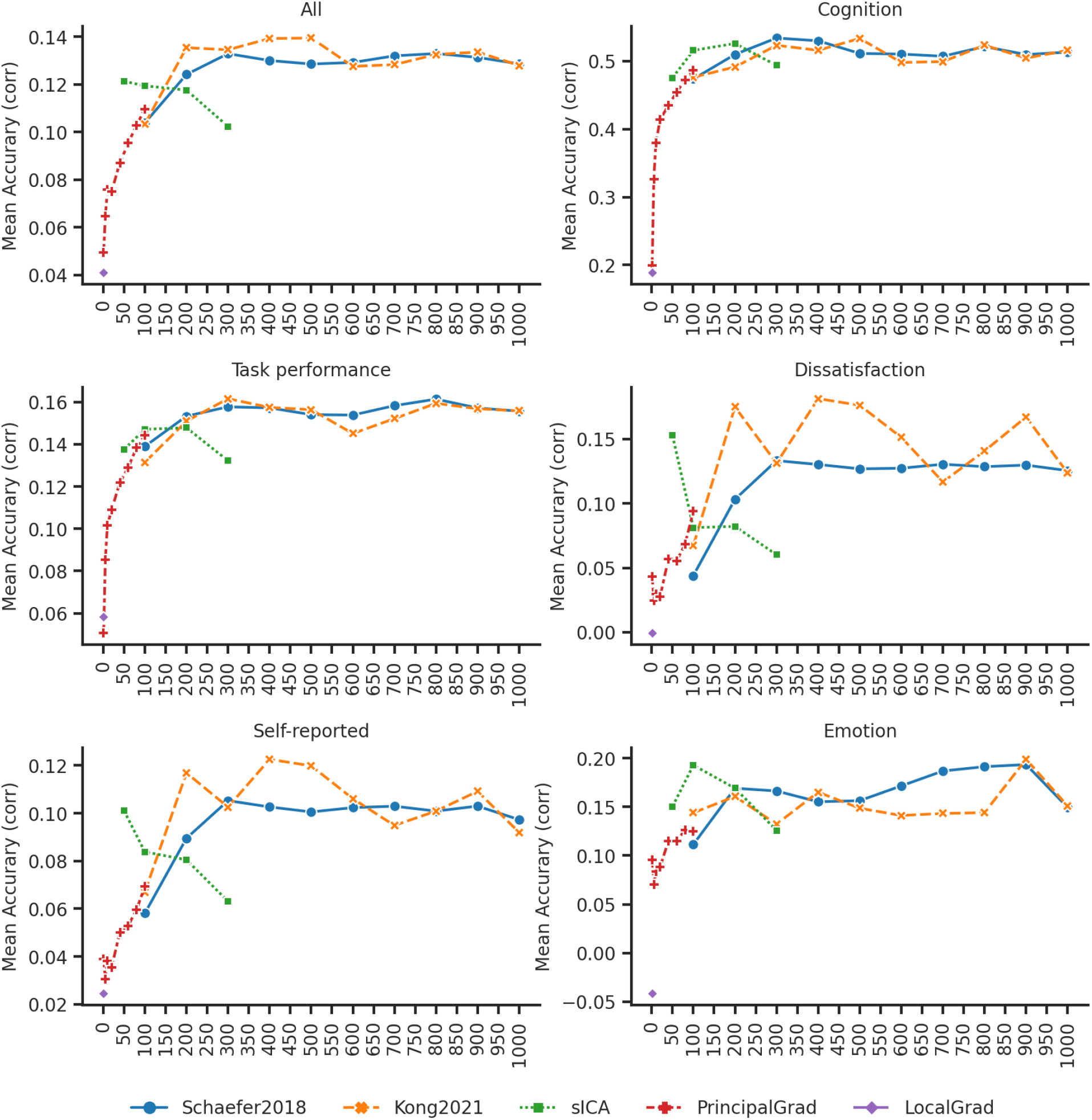
Prediction accuracies (Pearson’s correlation) vary across resolutions for gradient and parcellation approaches using LRR in the HCP dataset.

**Figure S42.**
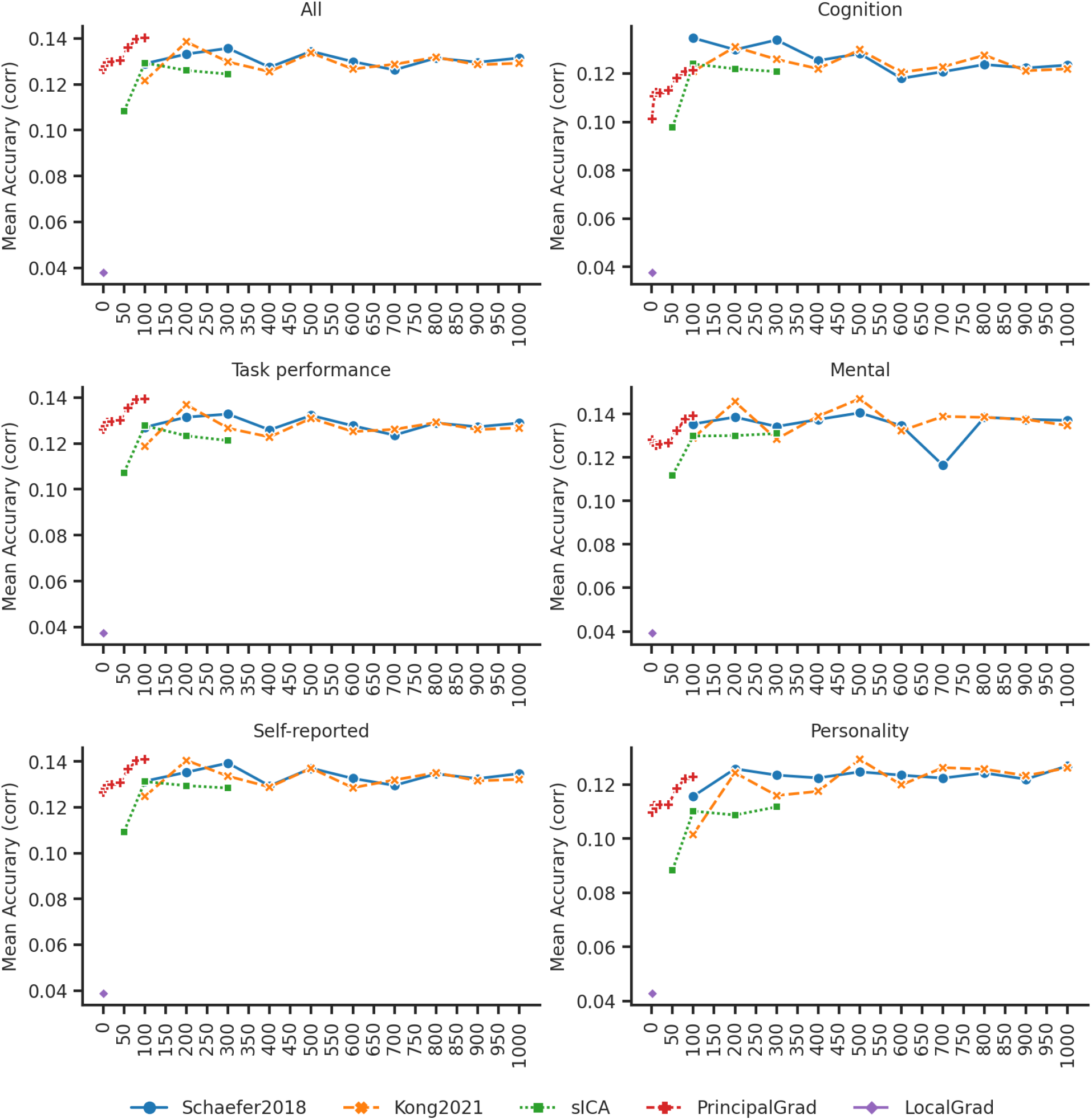
Prediction accuracies (Pearson’s correlation) vary across resolutions for gradient and parcellation approaches using LRR in the ABCD dataset.

